# Sex-Specific Genes Identified in Sea Urchin Gonads are Expressed Prior to Metamorphosis

**DOI:** 10.1101/2022.01.26.477926

**Authors:** Cosmo Pieplow, Gary M. Wessel

## Abstract

A great collaboration between the germline and somatic cells of an organism is the creation of a functional gonad. Here we begin to test this mechanism in a sea urchin by use of RNA-seq, and quantitative mRNA measurements throughout the life cycle of the sea urchin, *Lytechinus variegatus (Lv)*. We found through de novo analyses that the ovary and testis of this echinoderm contains the many transcripts predicted for gamete morphology and function, but we also discovered many uncharacterized gene products unique to each gonad. We found that transcripts involved in glycolysis are highly enriched in the ovary compared to the testis, and that *Lv* has an expansion of nanos genes not observed in other sequenced echinoderms. We developed a pipeline integrating timepoint RNA-seq data throughout development to identify hallmark gene expression in gonads. We found activation of meiotic genes surprisingly early in development, and observed that candidate genes involved in sex determination are first expressed during larval growth, well before metamorphosis. We further discovered that individual larvae express varying levels of male- or female-hallmark genes before metamorphosis, including factors for the germ line, oocyte, sperm, and meiosis related genes. Together these data support the hypothesis that sex determination in the sea urchin is initiated prior to metamorphosis, from a shared profile of male and female factors, and that the many uncharacterized genes unique to the gonads may reveal unique pathways and mechanisms in echinoderm reproduction.

**Highlight:**

- Testis and Ovary of the sea urchin, *Lytechinus variegatus*, have distinct transcriptome profiles
- *L. variegatus* gonad RNA-seq reveals the expression of 9 nanos genes
- The ovary in the sea urchin uniquely expresses genes of the glycolytic pathway, including an ovary-specific transcript of GAPDH
- Genes involved in meiosis and in sex determination are expressed early in larval development
- From gene expression profiles we conclude here that sex determination is initiated prior to metamorphosis in this echinoderm

## Introduction

The cyclical nature of sexual reproduction requires distinct gametes, the reproductive units of the male and female, to meet, fuse, and initiate development of an embryo (Hertwig, 1875; Just, 1930, 1936). Within each sexually reproductive organism, the cellular lineage giving rise to gametes, called the germ line, usually originates early in development with the partitioning of primordial germ cells (PGCs) from the rest of the embryo, herein referred to as the soma (Simonelig, 2012; Weismann, 1892; Wilson, 1904; Zou, 2015). PGCs usually form in one region of the embryo, whereas the somatic cells that give rise to gonadal tissues, referred to as a “bipotential primordium,” develop elsewhere (Bendel-Stenzel, Anderson, Heasman, & Wylie, 1998; Capel, 2017; Chiquoine, 1954; Raz, 2004). The PGCs then migrate to the primordium sites where they invade the nascent, bi-potential gonad, and proliferate to become the replicative stem cells for gametogenesis (Anderson, Copeland, Scholer, Heasman, & Wylie, 2000; Nicholls et al., 2019).

Following the arrival of PGCs to the bipotential primordium, the gonad differentiates to become either male, female, or both, before it is able to participate in gametogenesis (King, Gut, & Zenker, 2020; Lin & Capel, 2015; Lucas-Herald & Bashamboo, 2014; Nef, Stevant, & Greenfield, 2019; Weber & Capel, 2018). While gametes are front and center in the event of sexual reproduction, the somatic cells that make up the gonad are essential for germline differentiation into egg or sperm (Jost, 1948; Rey, Josso, & Racine, 2000). Importantly, the somatic cells of the gonad are essential for sex differentiation, maintenance, and the continued fertility of the organism (Jost, 1970). Somatic cells of the testes are responsible for the production of the (often trillions of) spermatozoa an adult male will make during his reproductive lifetime (Bujan et al., 1989). The somatic cells of the ovaries govern development of the egg, the largest cell of the adult body, by sequestration of nutrients, mRNAs, and accumulation of organelles within the growing oocytes (Hikabe et al., 2016; Just, 1930; Wessel et al., 2014). Therefore, the soma of the testis and ovary, although derived from the same bipotential primordia, have distinct functions and developmental trajectories able to propagate gametes following the establishment of their sexual identity (Capel, 2017).

In the sea urchin, *Lytechinus variegatus* (*Lv*), adult animals are gonochoristic, meaning they are either male or female, not both, and the adult animals contain either ovaries or testes based on their determined sex. While much of early embryonic development in the sea urchin is under intense investigation (McClay, 2011), the origin and composition of the adult organs and how they arrive at either sex, is poorly understood. Here we present a study diving into the biology and functional transcription of ovarian-and testes genes, their timing, and expression in the life cycle of this classically used sea urchin.

## Results

### Sex-enriched and Sex-specific transcripts align with Gonad Function

Results of RNA-seq differential gene expression (DEG) analysis reveal testis- and ovary-unique and enriched transcripts in the sea urchin, *Lytechinus variegatus* (*Lv*). Deseq2 analysis (Love, Huber, & Anders, 2014) returned 101 unique and 1,437 significantly enriched genes among *Lv* ovaries (N=3), and 361 unique 1,066 significantly enriched genes among *Lv* testes (N=3), respectively (Figure 2A and B). Interestingly, when the gonad transcripts unique to each sex are examined, as many as 50% had no annotated transcript name or predicted orthology (Supplemental Figures 1&2). To aid in our understanding of possible sex-specific transcripts, the top 16 of these unknown transcripts were annotated using protein domain predictions, 8 male, and 8 female (Supplemental Data). The female unique novel genes were quite variable, including a cell-surface signaling protein, a glycosyltransferase, and a methyltransferase, among others. The male specific novel genes included many transmembrane proteins, likely orphan GPCR’s (Supplemental Data).

**Figure 1.**
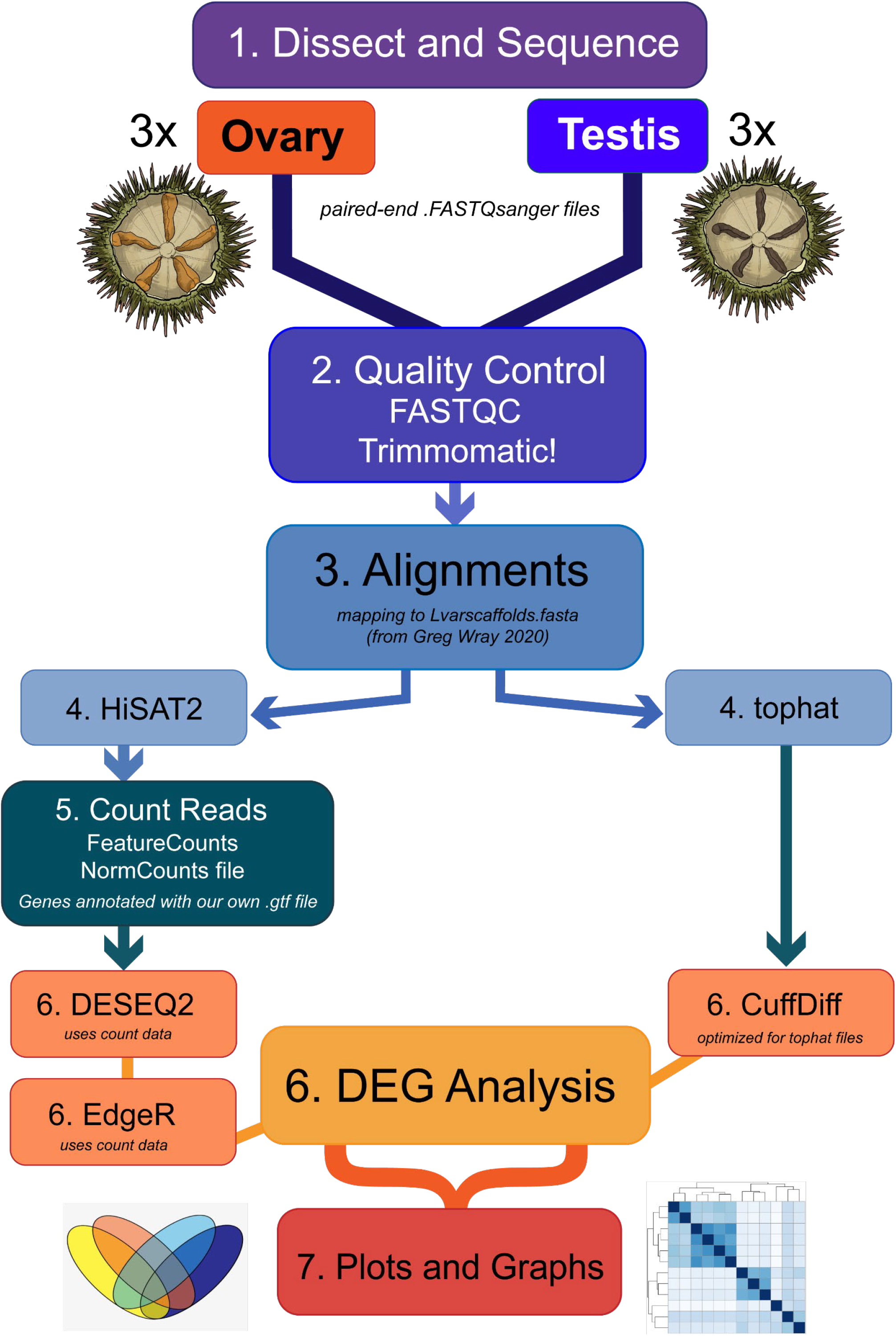
Workflow of Analysis. An overview of the methods used to obtain and analyze RNA-seq data from ovaries and testes of *Lytechinus variegatus.* 1. Three replicates of each sex were obtained by dissecting and preparing total RNA isolates from Ovaries and Testes of live *L.variegatus* sea urchins. 2. Once .FASTQ sequences were returned, six were trimmed using Trimmomatic! Software, And Quality Control reports were obtained using FastQC. 3. Trimmed and cleaned .FASTQ files for all gonads were aligned using two pipelines, separately, and further compared downstream. 4. HiSAT2 was used for alignments to obtain count data, and tophat was used for alignments meant for CuffDiff analysis. 5. Featurecounts was used to produce count files for DESEQ2 and EdgeR statistical analysis platforms. 6. Differential gene expression was obtained and compared using three unique statistical packages. 7. Result files and gene matrices were then plotted and presented respectively.

**Figure 2.**
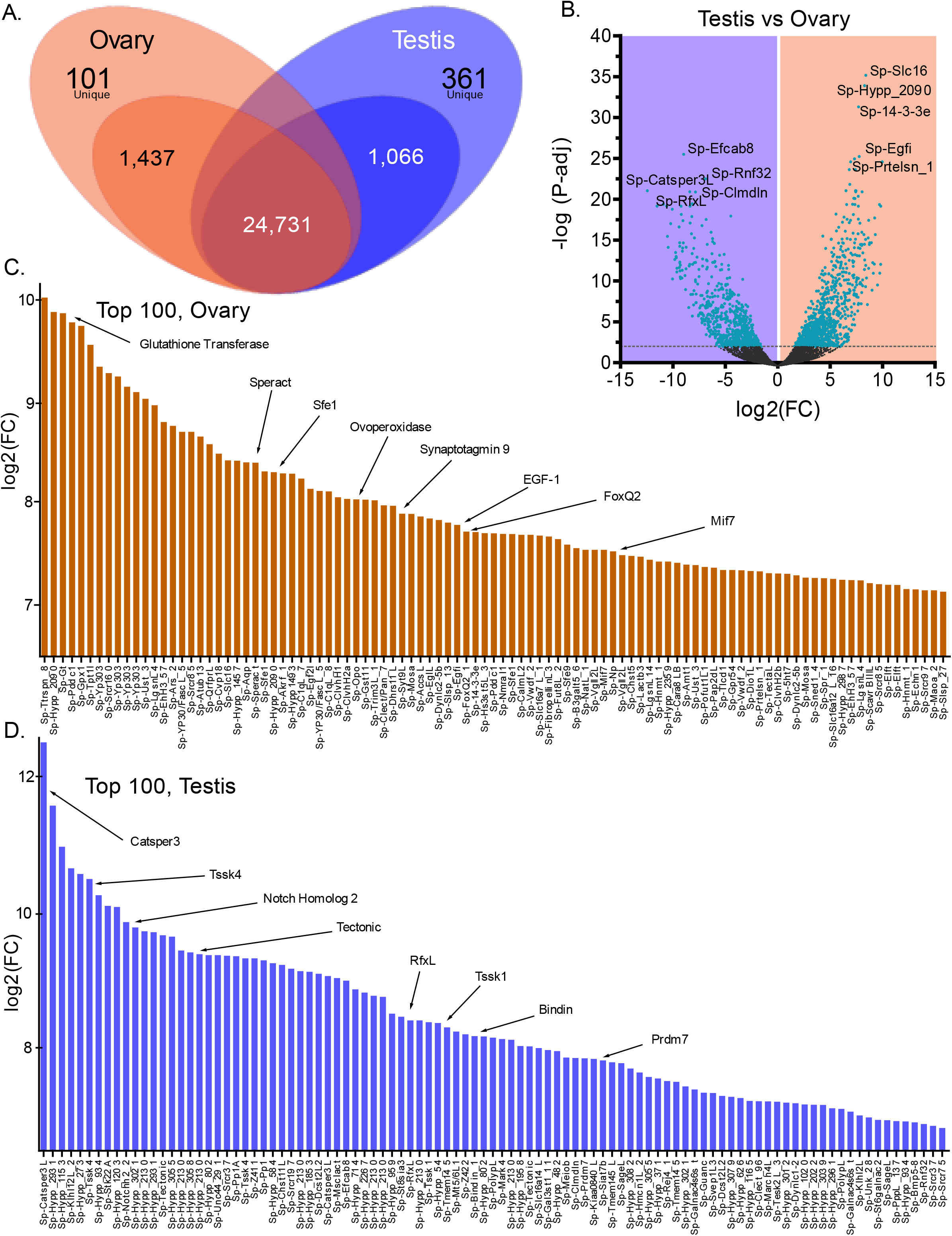
Results of Ovary versus Testis RNA-seq. A. Venn diagram summarizing total unique genes for each sex/gonad, significantly enriched genes for each sex, and shared/non-significant genes for all six gonad samples analyzed. Non-significant genes were omitted from any further downstream analyses. B. Volcano plot of all significantly enriched genes, Y-axis is P-value (significance) and X-axis is log(FC) expression values for each gene (plot points). Left-side genes are those enriched in testes samples, and right-side genes are those most significantly enriched in the ovary. C. Histogram of relative expression for top-100 most significant ovary enriched genes, genes of interest are highlighted with small arrows. D. Histogram of relative expression for top-100 most significant testis enriched genes.

When Deseq2 results are plotted with p-values as the y-axis, several transcripts emerge as being the most significantly enriched. Among the top 100 most significantly enriched transcripts in the ovary: speract, an egg secreted protein (Hansbrough & Garbers, 1980), ovoperoxidase, an egg-specific enzyme (Foerder & Shapiro, 1977), and EGF, a key ovarian follicle regulator (Bennett, Osathanondh, & Yeh, 1996) are among the most expressed, indicating that the clustering of ovary-enriched transcripts support the functional transcription expected for the resultant eggs (Figure 2C). Similarly, in the testis: Catsper (Hildebrand, Avenarius, & Smith, 1993; Xia, Reigada, Mitchell, & Ren, 2007), a sperm specific ion channel, Bindin a sperm protein (Glabe & Vacquier, 1977), and Testis-specific serine-threonine kinases (Hao et al., 2004; Zuercher, Rohrbach, Andres, & Ziemiecki, 2000) are among the top enriched transcripts (Figure 2D). This initial scan of ovary- and testis-specific gene expression supported authentication of gonad gene expression for further analyses.

Our initial results of DEG analyses provide support to the hypothesis that the testes transcribe massive amounts of cytoskeletal transcripts needed to primarily produce flagellated spermatozoa (Bannai, Yoshimura, Takahashi, & Shingyoji, 2000; Schatten & Schatten, 1981). Testis specific serine-threonine kinases (TSSK’s) are also present, though these genes are of unknown functions in echinoderms. Additionally, calcium-signaling and meiotic genes are also incredibly abundant, their functions all in-line with a sperm producing organ.

We found that ovaries express transcripts associated with RNA-binding and metabolism, likely involved in their role of generating huge volumes of oocytes, with RNA-binding, translation, metabolism, and ribosomal function (Figure 3A & 4A). Each of these pathways ties directly into the primary role of the ovary: the production of RNA-rich oocytes with maternally-loaded transcripts, capable of driving early development (Just, 1930; Simonelig, 2012; Tadros & Lipshitz, 2009)

**Figure 3.**
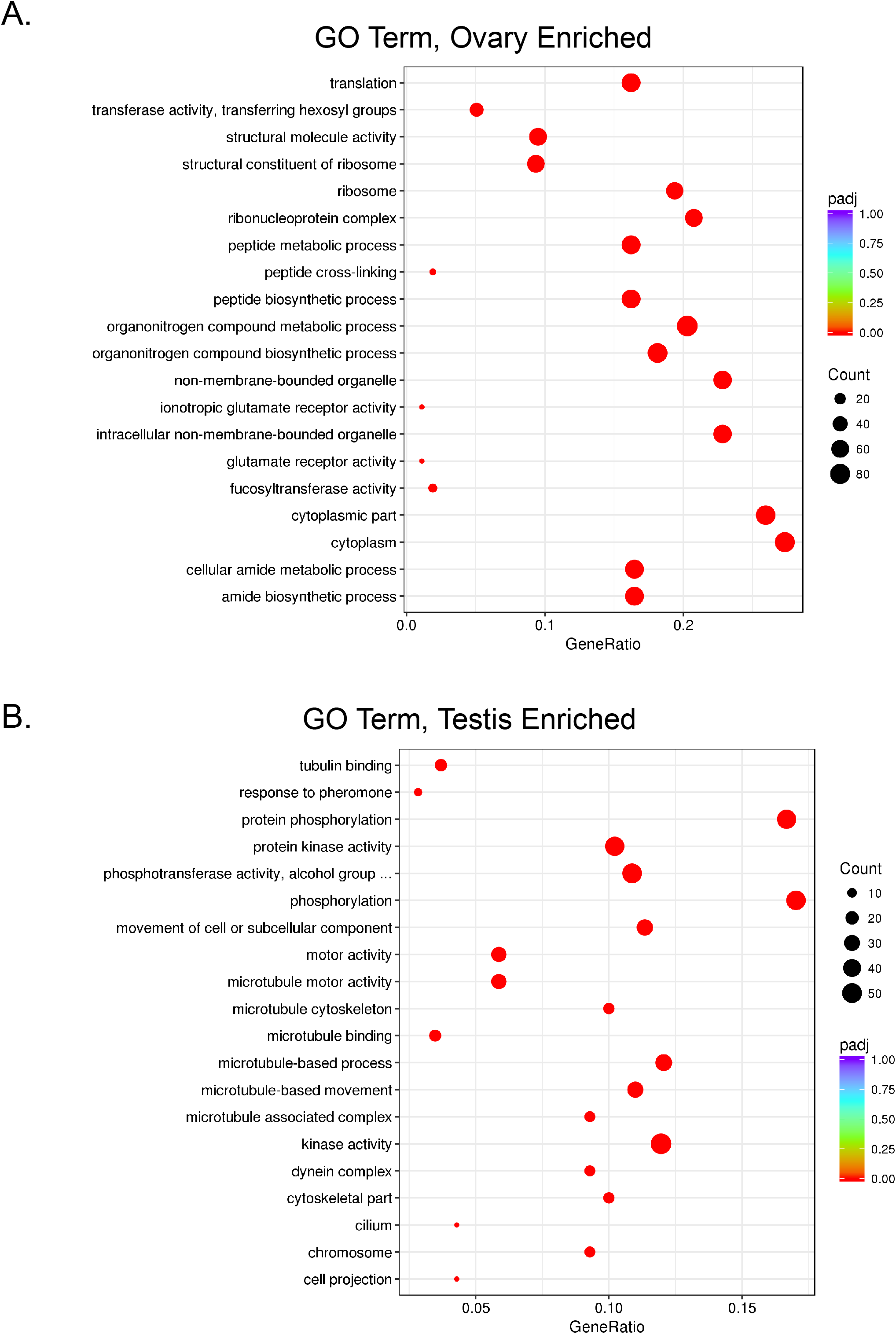
Ovary and Testis Gene Ontology Enrichment Analysis. A. Gene Ontology Bubble plot of GO terms significantly enriched among the three ovary samples. Color represents P-value (significance) while size of the bubble represents number of transcripts detected above threshold for each GO term. B. Gene Ontology Enrichment Bubble plot of GO terms significantly enriched among the three testes samples.

**Figure 4.**
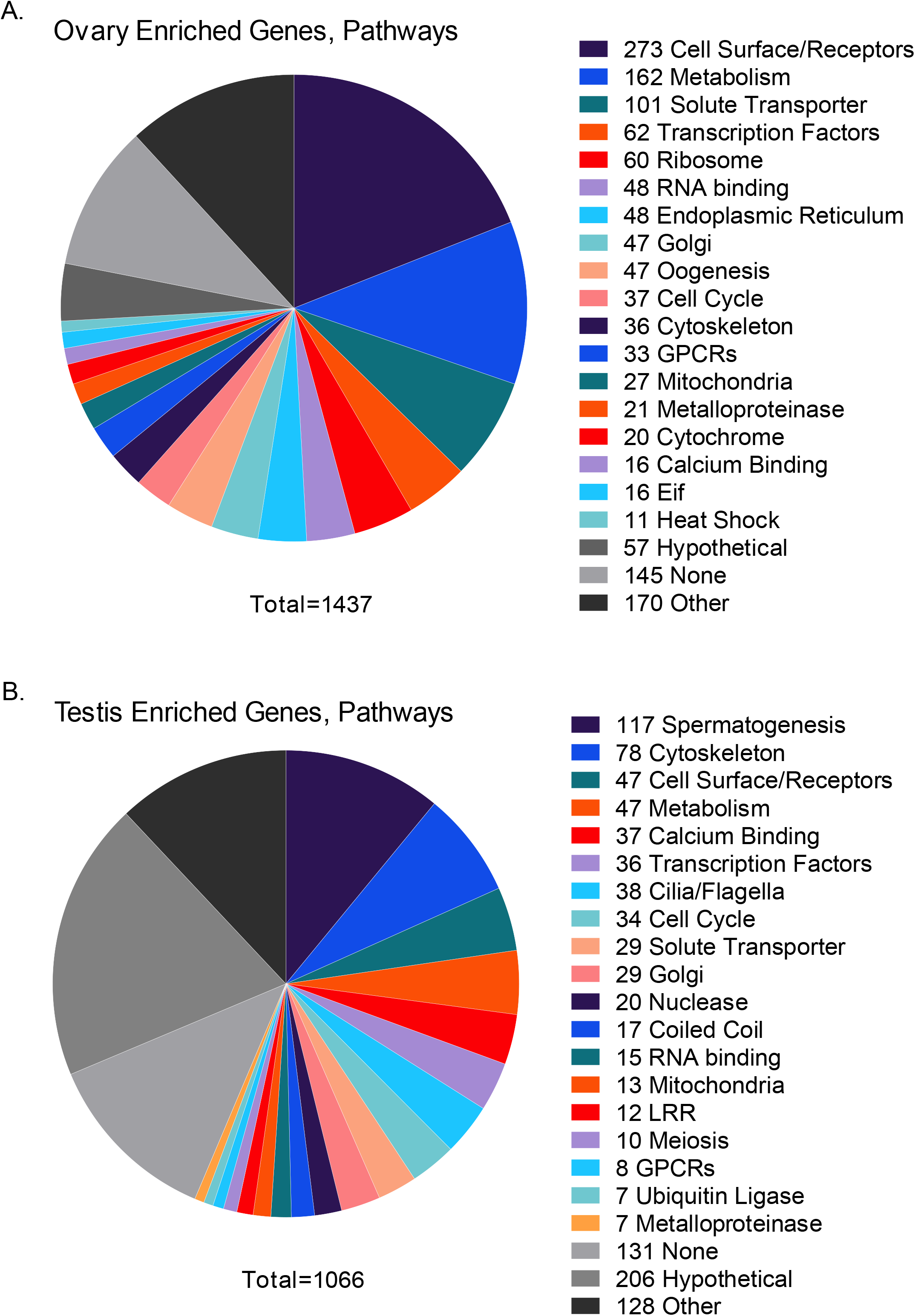
Ovary and Testis Enriched Genes and Pathways. A. Pie Chart of Gene Groups and pathways enriched in the ovary, only significantly enriched genes were used in this analysis, and pathways are summarized by each color to the right side of the chart. Ovaries were most enriched for transcripts involved in cell surface receptors, metabolism, solute transport, and RNA-binding. B. Pie Chart of Gene Groups and pathways enriched in the testis. Testis pathways most enriched included genes involved in spermatogenesis, cytoskeleton/microtubule binding, and calcium signaling.

Similarly, for the testis, enriched pathways included tubulin binding, cytoskeleton, microtubule function, and calcium binding (Figure 3B & 4B). Each of these pathways is implicated in the process of spermatogenesis, and reveals genes in the production of flagellated sperm (Goldsmith et al., 1991). We conclude from these data that we have performed a robust DEG analysis, whereby sex-specific transcripts have been successfully filtered by their expression according to male- or female-sex in each gonad.

### Glycolysis is an enriched pathway of the ovary

Glycolysis is an essential metabolic pathway for normal tissue growth and homeostasis. Much to our initial surprise, our analyses revealed a marked enrichment of glycolytic pathway transcripts in *Lv* ovaries (Figure 5A). Importantly, several transcripts for glycolytic and gluconeogenic enzymes, as well as the IR (insulin receptor), are both highly enriched in ovaries, and not detectable in the testes of *Lv*. This finding supports the hypothesis that glucose metabolism via the insulin receptor is involved in oocyte production and oocyte maturation in this organism (Sutton-McDowall, Gilchrist, & Thompson, 2010) (Figure 5C).Glyceraldehyde 3-phosphate dehydrogenase (GAPDH) is an essential metabolic enzyme, present in nearly every species (Lima et al., 2009), and it catalyzes the 6^th^ step in the canonical pathway of glycolysis. Historically, GAPDH has been used as a loading control for protein abundance in Western Blot analyses, and as an mRNA transcript abundance control in many RNA-based assays (Rimarachin, Norcross, Szabo, & Weksler, 1992; Slagboom, de Leeuw, & Vijg, 1990). However, this RNA-seq analysis of gonads in Lv has revealed that GAPDH, in addition to nearly every canonical glycolytic enzyme transcript, is either unique to, or highly enriched, in the ovaries of the sea urchin (Figure 5A&B). Like most metazoans, Lv has multiple transcripts for the peptides making up GAPDH subunits (Baalmann, Scheibe, Cerff, & Martin, 1996), totaling four transcripts in this species of sea urchin. More striking, is that of the four GAPDH transcripts, three of the four are not detected at all in the testes, but are expressed abundantly by the ovary, and the fourth is detectable in the testis, but remains markedly enriched in the ovary (Figure 5B).

**Figure 5.**
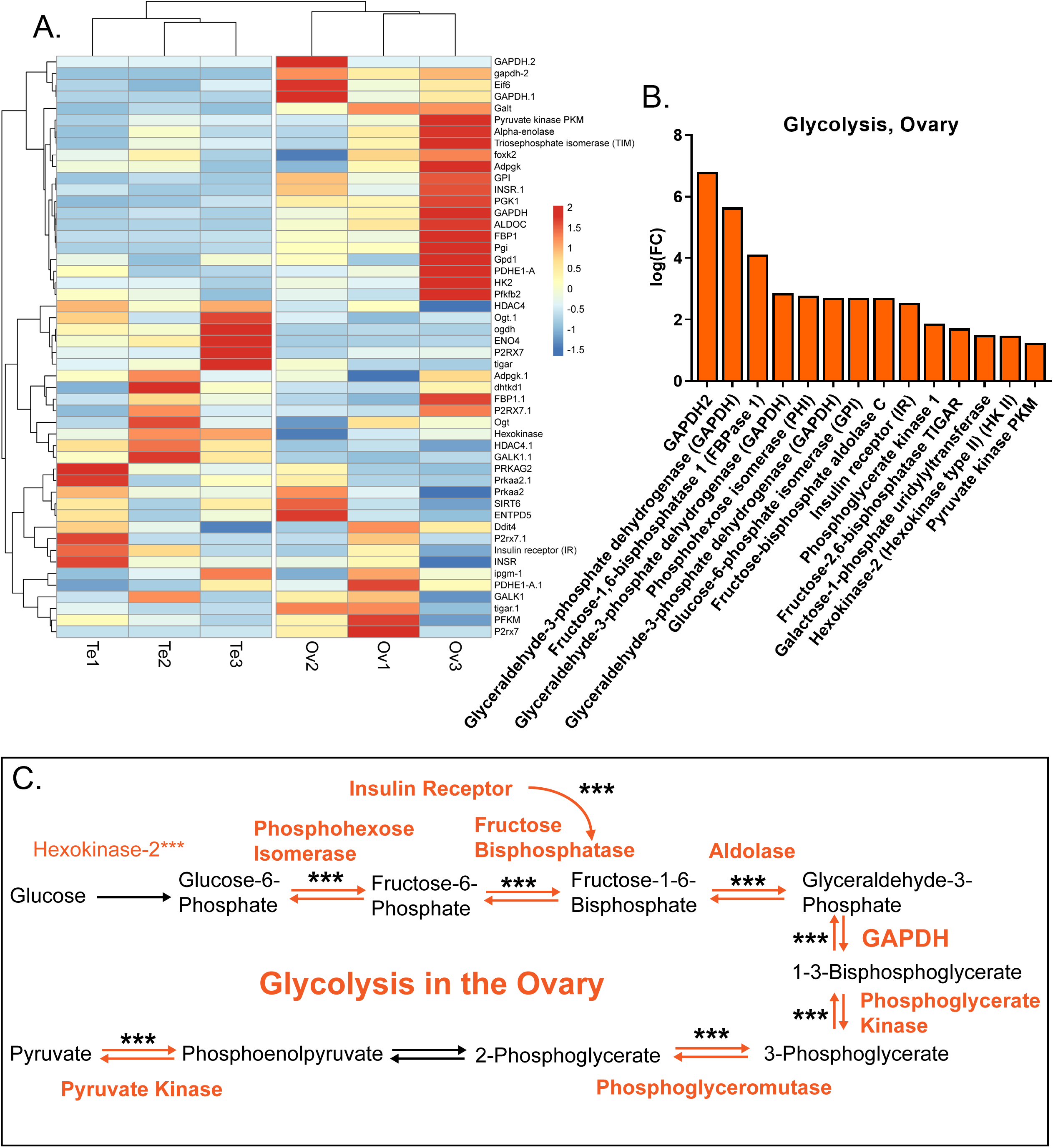
Glycolysis is a Pathway of the Ovary. A. Heatmap showing sex-specific expression of glycolysis pathway genes in gonads of both sexes. Color hue and intensity represent relative FC values for each gene in each gonad. B. Fourteen glycolysis-specific genes were significantly enriched (adj. p-val<0.0001, FC>1.5) in the ovary when compared to the testis, while none of the genes involved in the canonical glycolytic pathway were significantly enriched in any of the testes samples. C. A summarized schematic of the canonical pathway of glycolysis, with ovarian enriched or unique genes highlighted in orange. Significance is denoted with asterisks above each enzyme gene in the pathway. Nearly all major enzymes utilized in glycolysis is either significantly enriched in, or unique to the ovary in *L.variegatus*.

The glycolytic and gluconeogenic enzymes appear to be a “hallmark” of ovarian expression in this species of sea urchin. The enrichment of IR in the ovary supports the hypothesis that it is not one strict pathway over another, but rather the regulation of glycolysis and glycogen production that is needed for normal oocyte development. This data supports the consensus of glycolysis being an abundant pathway of the ovary, as it is observed in other animals, including mammals (Biggers, Whittingham, & Donahue, 1967).

### Testes and Ovary express unique collections of germline genes

In addition to metabolism, we sought to parse out transcripts related to germ cell fate and germ cell maintenance. To address this question, we simply filtered all differentially expressed transcripts by GO term (GO= germ cells) and plotted expression values for all major germ cell specific genes detected by RNA-seq.

With an initial look at heat maps, it is striking that male-biased (testis) expression of germline genes tend to cluster tightly, with most germline genes expressed at similar abundance. In contrast to the testes generally similar expression of the germline genes, ovaries have widely variable expression between individuals. Strikingly, while germline gene expression is widely variable in female Lv, the functions of these groups of ovarian - enriched genes are similar (Figure 6A).

**Figure 6.**
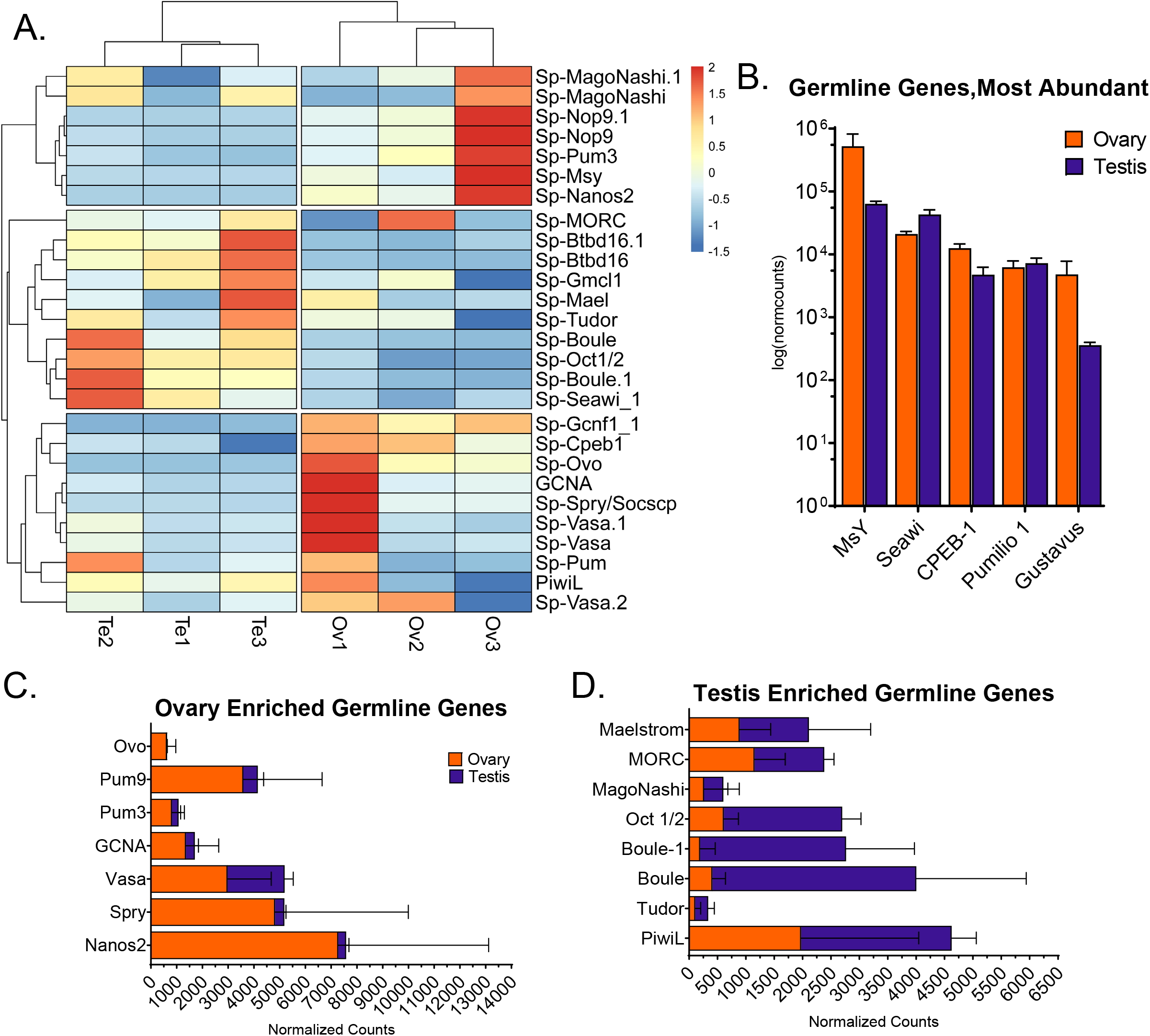
Germline Genes Cluster based on Sex or Abundance. A. Heatmap providing an overview of all germline gene expression within gonads. B. Logarithmic-scale expression bar chart of germline genes from gonad data. The five most abundant germline genes observed in the gonads were MsY, Seawi, CPEB-1, Pumilio1 and Gustavus. Note sex-specific biasing of expression levels: MsY and Gustavus are 10 times more abundant in ovary, while Seawi is only 2 times more abundant in testes. C. Bar charts of germline genes expressed by the ovary. Error bars represent standard error on expression values for each sex. Ovary values = orange, Testis =purple. D. Germline genes significantly enriched in, or unique to testes. Error bars represent standard error on expression values for each sex. Ovary values = orange, Testis =purple.

Ovaries are enriched for the germline genes nanos, vasa, germ cell nuclear antigen (GCNA) (Carmell et al., 2016), and sprouty (Spry) (Casci, Vinos, & Freeman, 1999), among others. It is known that nanos and pumilio work together (Parisi & Lin, 2000) for the essential translational repression observed in female germ cells, and it was not surprising to find multiple pumilio transcripts enriched in the ovary as well. Importantly, however, many of the ovary enriched germline transcripts are maternally loaded mRNAs important for mature oocyte function (Sugimori, Kumata, & Kobayashi, 2018).

Testes, on the other hand, are enriched for Boule, essential for normal meiotic progression (Sekine, Furusawa, & Hatakeyama, 2015; Xu et al., 2003) and Oct1 (Williams, Cai, & Clore, 2004) which is likely a regulator of spermatogenic stem cell fate in the testis of *Lv* (Figure 6C).

All gonads, regardless of sex, express incredibly high levels of the transcripts for Gustavus (Gustafson, Yajima, Juliano, & Wessel, 2011), Seawi (Rodriguez et al., 2005), and Pumilio, (Swartz & Wessel, 2015; Wessel et al., 2014) among others (Figure 6B). This analysis yielded a list of functionally relevant germline genes that may bear essential roles in gamete production in the gonads of adult *Lv* (Ponz-Segrelles, Bleidorn, & Aguado, 2018). Importantly, these findings now also allow us to segregate germline transcripts by sex.

### *L. variegatus* has an expansion of Nanos genes not observed in other Echinoderms

It is well known that Nanos (Bhat, 1999; Kobayashi, Yamada, Asaoka, & Kitamura, 1996) is an essential translational regulator gene needed for germline stem cell maintenance and germ cell identity (Juliano, Yajima, & Wessel, 2010; Oulhen & Wessel, 2016). While pursuing our pathway analyses of germline gene expression in *Lv* gonads, we were puzzled that ten nanos transcripts returned after filtering. From our RNA-seq analysis, we identified ten transcripts, all annotated as “Zinc Finger Nanos,” with nine of the ten returning a bona fide Nanos protein after translation and protein structure prediction software (Figure 7A). While it was surprising that there are nine nanos transcripts in *Lv*, most interesting of all, however, was the tenth transcript: Lv_19334, which returned a previously uncharacterized transposase (Figure 7A). We decided that further analyses were needed, so pursued an evolutionary analysis using the well-characterized nanos transcripts in the closely related sea urchin species, *S.purpuratus (Sp)* for comparison.

**Figure 7.**
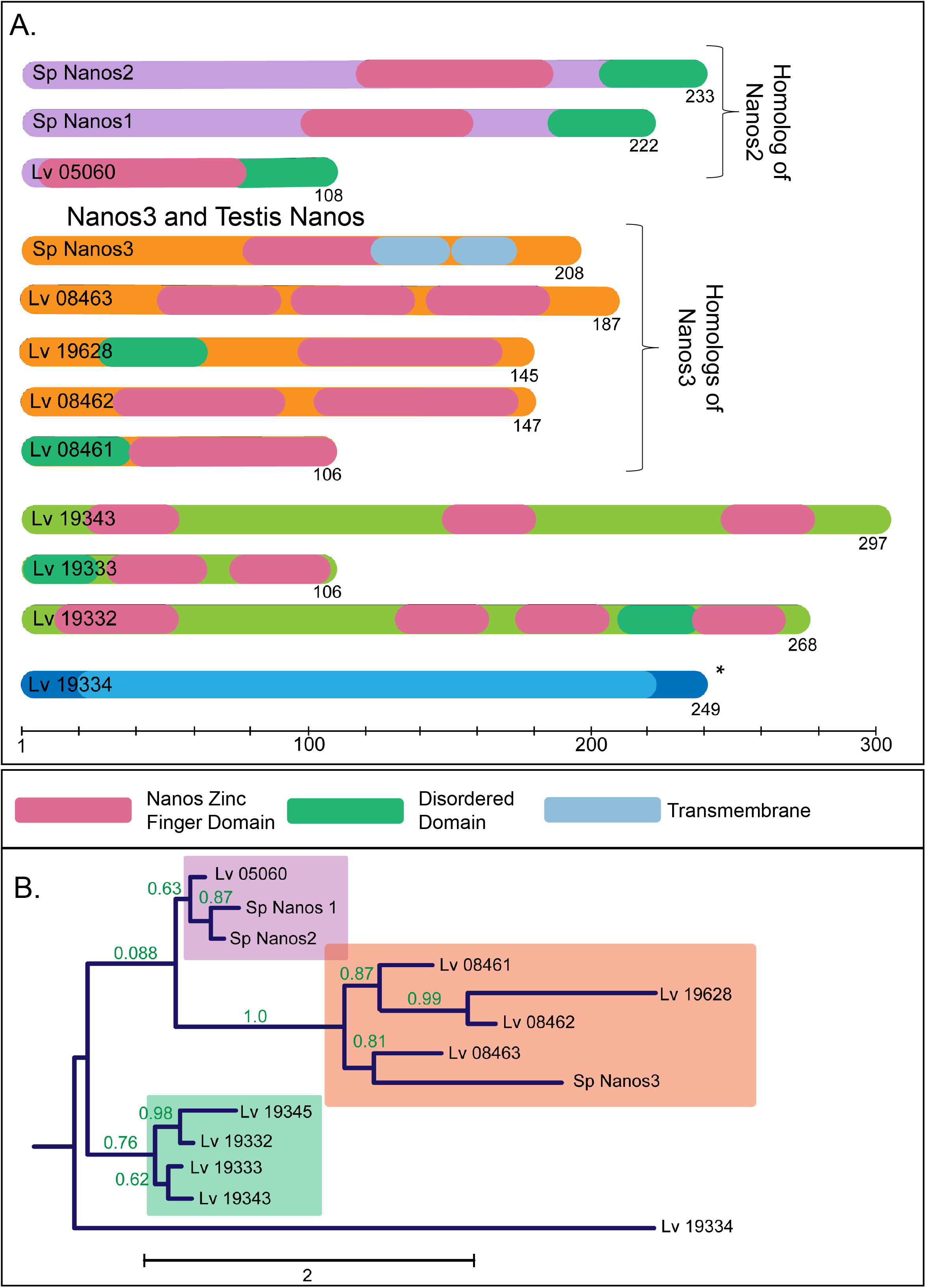
Identification of 10 Nanos Transcripts from RNA-seq Data. A. An evolutionary tree including major extant groups of echinoderms. Nanos genes for each class were obtained with assembled genome and transcriptome data. *L.variegatus* is highlighted as it represents a unique expansion of Nanos gene transcripts, totaling nine nanos transcripts, when compared to other Echinoids. B. Chromosomal map view of Nanos genomic loci in *Lytechinus variegatus.* Grey represents an inferred centromere and rounded/colored ends represent inferred telomere sequences on each respective chromosome. Nanos transcripts identified through RNA-seq each mapped to unique chromosomal loci, on chromosomes 2, 4 and 12. Interestingly, Transcript 19334 and 19343 map to the same genomic location, however, transcript 19343 is transcribed antisense, and encodes a transposase, not a Nanos protein. C. Heatmap of sex-biased expression of the 10 nanos transcripts identified through RNA-seq. Note messy sex-biased expression of the nanos transcripts, with several being expressed preferentially by testes. D. qPCR validation of nanos transcripts 08461 and 19628. Plot shows mRNA expression over developmental time, using nanos2 (05060) as a reference. Transcript 08461 mimics Nanos2 expression but is not expressed abundantly in the ovary. E. qPCR analysis of nanos transcripts 19332, 19333, 19334 and 19345 mRNA expression over time, using nanos2 as a reference. Transcripts in this group are expressed variably, though later in developmental time. However, Transcript 19334 mimics Nanos2 expression with a slight developmental delay.

For ease of reference, these peptide alignments and sequence identity comparisons are shown with the structures of well-characterized *Sp* Nanos proteins (Juliano et al., 2010) (Figure 7A). These alignments produced several previously undescribed clusters of nanos genes in Lv (Figure 7A). The first, containing Lv_05060, includes the direct Lv homolog of Sp Nanos2, the essential germline regulator in ovaries and early embryos (Figure 7B). Another cluster, *Lv* transcripts sharing the most sequence identity with Sp Nanos3, a Nanos gene that is detected in the testes of *Sp*. These Nanos3 genes, Lv_08463, Lv_19628, Lv_08462, and Lv_08461 are also all detected in the testes, as would be expected for Nanos3 genes, as there is evidence they may play a role in the identity of spermatogonial stem cells (Lolicato et al., 2008). The third cluster contains nanos transcripts that share significant sequence similarity with one another, but not with any *Sp* homologs. Notice that the outgroup, Lv_19334, the uncharacterized transposase, does not share significant sequence similarity with any of the nine bona fide nanos transcripts in Lv (Figure 7B).

After interrogating the sequence homology of the mRNA and predicted protein structures, we pursued further characterization of the nine nanos transcripts in *Lv*. It is known currently that all extant echinoderm species that have been sequenced, have three, at most four, nanos genes (Figure 8A). The question remains as to whether these nine transcripts in *Lv* are functional nanos proteins, involved in germline establishment or maintenance, or simply gene duplications that are the result of this transposon. It can be noted that in some vertebrate species, a diversification of functional nanos genes has been observed, and is essential for germline establishment (Sun et al., 2017).

**Figure 8.**
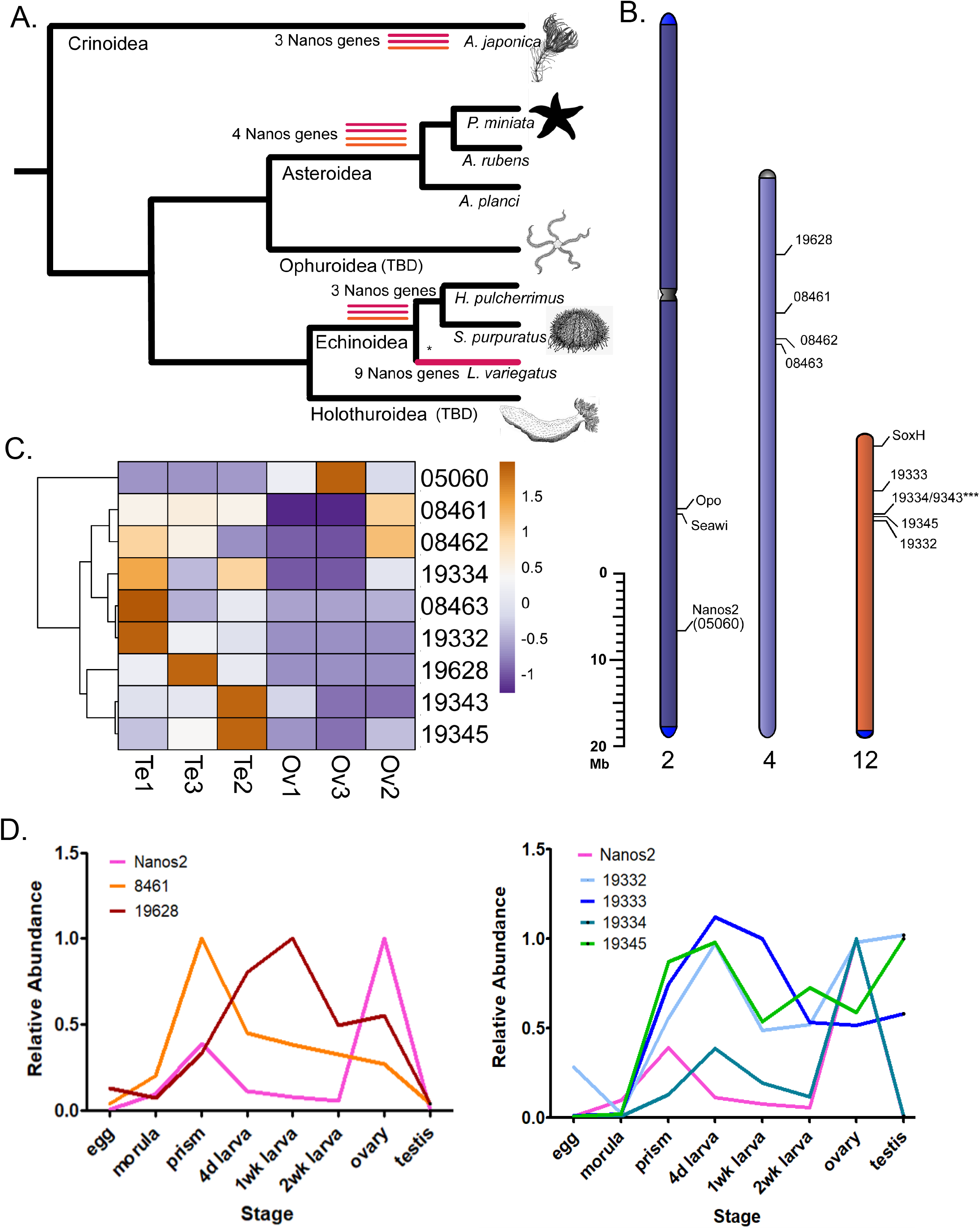
A unique transposase is the possible explanation for Nanos gene expansion in *variegatus*. A. Schematics of all predicted peptide sequences obtained from all nanos transcripts detected in the *L.variegatus* RNA-seq data. Schematics are grouped based on their homology to *S. purpuratus* Nanos proteins, with Nanos2 at the top, Nanos2 (an ovary-specific transcript) belonging to the same group, and Nanos3 as a separate group. The nanos transcripts associated with Nanos3 are typically expressed in the testes. The Nanos genes in the third group (green) share sequence identity with all nanos transcripts, but cluster independently of *S.purpuratus* homologs. Important domains are highlighted, pink = nanos zinc finger domain, and teal = predicted disordered domain. The final nanos transcript occupies the same genomic locus as transcript 19334, but is transcribed antisense, and encodes a unique transposase with no known homology. B. Sequence homology tree of the 10 predicted peptides for each of the Nanos genes in *L.variegatus*, compared to the three known *S.purpuratus* Nanos genes for sequence homology. Note that the transposase is a distinct outgroup, with no sequence identity to the other 9 Nanos genes.

To characterize these genes, we also examined the genome, and found that several of the nanos transcripts occupy the same locus within chromosome 12, and the novel transposase actually occupies the same locus as the transcript Lv_19343, just in an antisense orientation (Figure 8B). For the sake of clarity, Lv_05060 has been named “nanos2” as it is the bona fide homolog of Sp Nanos2 and is expressed at the same times in the same tissues. Nanos2 occupies a unique locus on chromosome 2, which also contains the nearby genomic loci of Ovoperoxidase and Seawi (Figure 8B).

Finally, the “male” Nanos genes, those *Lv* homologs of Sp Nanos3, each occupy unique positions on chromosome four (figure 8B). This revealed that, except for the strange transposase and its “partnered” nanos on chromosome 12, each Nanos gene in *Lv* occupies a unique genomic locus, and expresses unique transcripts. Most important of all, the genomic locus of the transposase (Lv_19334) and it’s “partnered” nanos gene (Lv_19343), implicates evidence of a possible gene duplication through transposable element activity on the chromosomes of *Lv*, allowing our hypothesis of a transposase generating an expansion of Nanos transcripts not observed in other echinoderms (Figure 8A).

The final question remained, are these nanos transcripts expressed in embryos, or only in the respective gonad tissues of *L.variegatus?* Through RNA-seq analysis, we observed many of these transcripts being expressed by either testes or ovary, with the Nanos3 homologs being expressed most abundantly in the testis, as expected (Figure 8C). Using qPCR analyses of distinct developmental timepoints, we validated the embryonic and larval expression of all the *Lv* nanos transcripts. Interestingly, the temporal expression of Lv_08461, a Nanos3 homolog, matches the temporal expression of Lv Nanos2 (Lv_05060), most closely during embryonic development (Figure 8D). Other Nanos3 homologs are expressed more abundantly than nanos2 during embryonic development, but are most abundantly expressed in the testes, as expected (Figure 8E). Together, these data paint a picture of nanos gene duplication in *Lv*, with an increase in the number of testes-specific Nanos genes. We hypothesize that this duplication was due to this associated transposase (Lv_19334) present antisense to the nanos gene Lv_19343, present on chromosome 12.

### Gonads exhibit Sex-Specific Expression of Transcription Factors

After our initial DEG analysis and identification of novel germline genes, we sought to identify any sex-biasing of transcription factor expression. It is known that certain transcription factors, namely FoxL2 (Huang, Ye, & Chen, 2017) in ovaries, and Sox9 or Sox30 (Bishop et al., 2000; Feng et al., 2017; Jakob & Lovell-Badge, 2011; Polanco, Wilhelm, Davidson, Knight, & Koopman, 2010; Zhang et al., 2018) in testes, are essential for gonad function and normal gamete production in mammals (Bai et al., 2018; She & Yang, 2017). Echinoderms have a FoxL2 ortholog and have a single Sox transcription factor representative that is orthologous to each Sox family in vertebrates. SoxE encodes the echinoderm ortholog of vertebrate Sox9, Sox8 and Sox10 (Tai et al., 2016; Weider & Wegner, 2017). Sox9 is an essential regulator of male fate, and in conjunction with Sox10, Sry, and Sox3, make up an ancient and essential pathway for testis differentiation (Bergstrom, Young, Albrecht, & Eicher, 2000). Echinoderm SoxH encodes the ortholog of Sox30, a key spermatogenesis regulator essential for normal meiosis in vertebrate testes (Bai et al., 2018).

To test the hypothesis that transcription factors for vertebrate sex determination would be conserved in echinoderms, we assayed all forkhead and Sox box family transcription factors as they were of greatest interest and plotted their expression patterns in the gonads of either sex. We found clean segregation in Sox box transcript expression, where testes express SoxE and SoxH abundantly, while ovaries are enriched for the genes SoxB1, SoxB2, SoxD and SoxC (Figure 9A). These data supported the hypothesis that SoxH and SoxE are important regulators of male gonad function (Bagheri-Fam, Sinclair, Koopman, & Harley, 2010; Lucas-Herald & Bashamboo, 2014). Testing this result with qPCR (data not shown) and in-situ hybridization supports the conclusion that SoxE and SoxH are male-specific transcripts and are expressed abundantly in the testes of *Lv.* Surprisingly, we found a low level of SoxH expression very early in embryonic development, at the mesenchyme blastula/early gastrula stage (Figure 9B&C). The Forkhead family of transcription factors were also assayed using DEG analyses, and the apparent lack of segregation of expression by sex was rather surprising. Most interesting of all was the lack of female biasing in FoxL2 expression; FoxL2 appears to be a factor expressed by both gonads. It is important to consider that while there is variable expression of all Forkhead family genes in the ovary, this expression is perhaps reflective of the variable status of overall oogenesis depending on the prevalence of oocyte stages in the ovary (Figure 9D).

**Figure 9.**
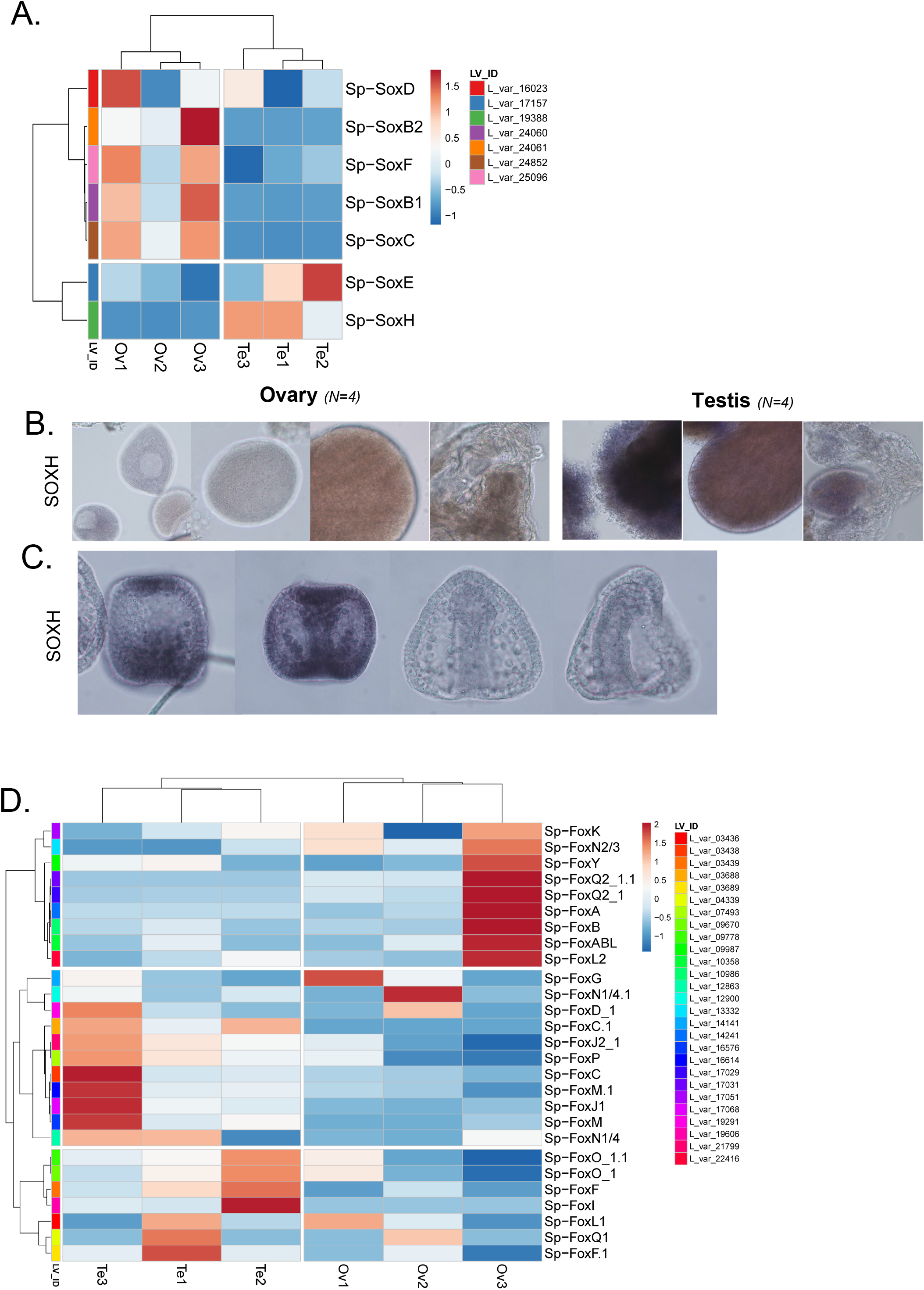
Sex Specific Expression of Transcription Factors. A. Heatmap summarizing gonad expression of Soxbox genes expression by sex. Color hue and intensity represent average relative fold change values. There is clear binary distinction in expression of Sox genes by each sex, with SoxH and SoxE observed only in testes, and SoxB genes highly expressed by ovaries. B. Validation of SoxH mRNA expression over developmental time by qPCR. SoxH represents a possible male sex factor, and is expressed exclusively by the testis, but is also expressed also early in development. C. Validation of sex-specific spatial expression of the transcription factor, SoxH. SoxH is only detected in the testis by in situ hybridization of whole-mount gonads. SoxH is also transiently expressd in early gastrulation, as also validated by in situ hybridization. D. Heatmap of Forkhead box transcription factors across gonads. Sex-Specific biasing of expression of Forkhead boxes is also observed.

### Identification of Gonad “Hallmark” Genes

We next attempted to determine which genes expressed by the gonads were actually unique to these tissues, or whether they were simply expressed broadly during embryonic development and only in adults become sex-biased. To dissect this question, we developed a pipeline to eliminate transcripts used during embryonic development and focused our lens on those present only in the gonads in adults. We first segregated data into two groups: embryonic transcriptomes immediately following zygotic genome activation (ZGA), and gonad transcriptomes of both sexes. To obtain transcripts most relevant to gonad function and biology in the adult organism, we selected those unique to- and enriched in-gonads, that were depleted in the post-ZGA embryo timepoint datasets (Figure 10).

**Figure 10.**
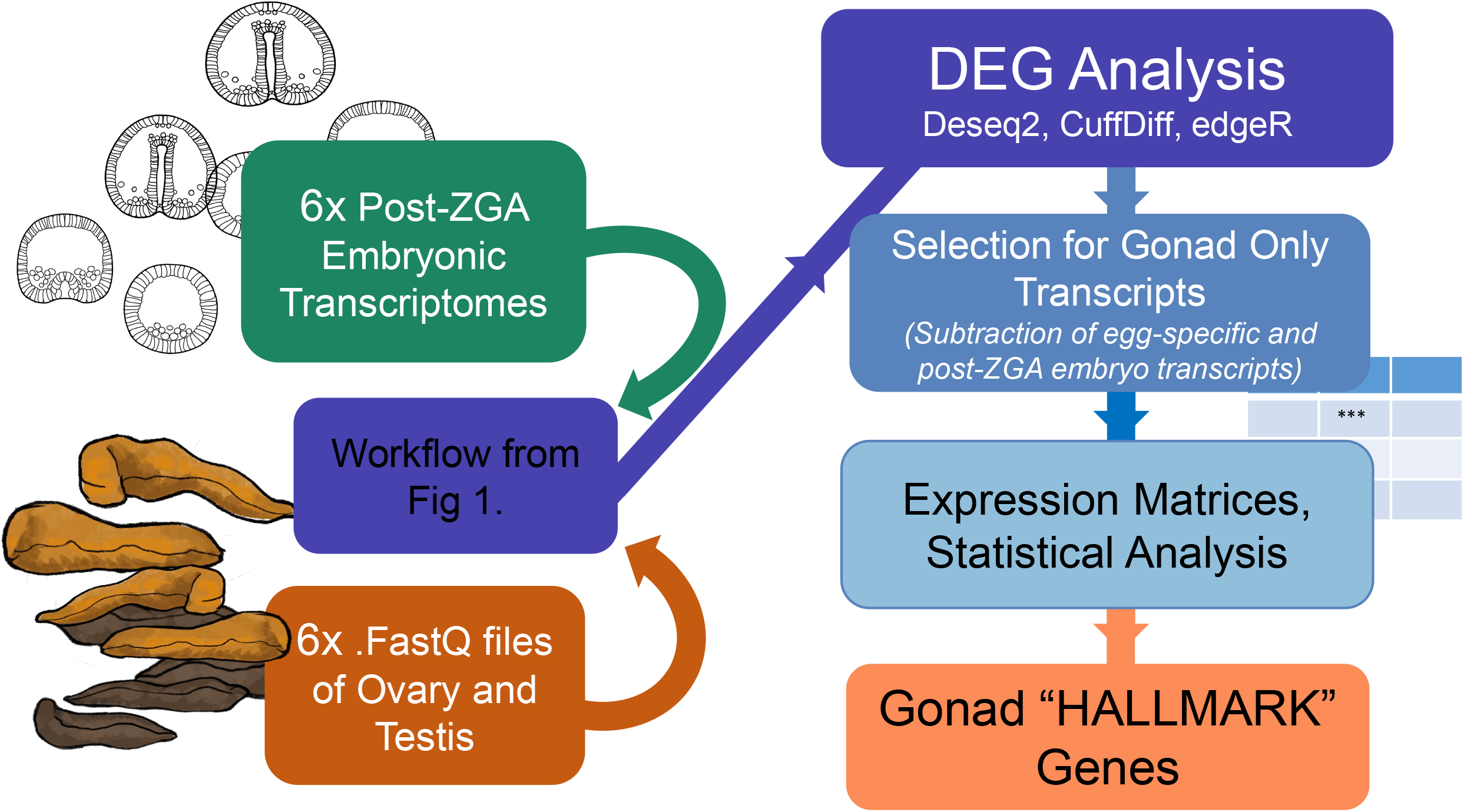
A six-way analysis to Identify Gonad Hallmark Genes. Flowchart for analysis conducted after Gonad-specific genes were obtained. In this 6-way analysis, six gonads (testes and ovaries) and six post-zygotic genome activation stage embryonic transcriptomes were put through the same pipeline in figure 1. Post-ZGA embryos were selected to not include egg-specific transcripts, and thus focus on genes expressed specifically by embryos in the late blastula to gastrula stage. Once expression matrices were obtained, non-significant genes were omitted as noise.

Following this “gonad hallmark” DEG analysis, we obtained 3,363 enriched gonad hallmark transcripts not expressed, or significantly depleted in embryo transcriptomes (Figure 11A). When sorted by lowest p-values, the genes most enriched in gonads included selenoproteins and major yolk proteins (MYP’s) (Figure 10B). The most enriched transcripts in the gonads of *Lv* included bindin, catsper, and testis specific serine-threonine kinases (Tssk’s) (Figure 11C). We also used this analysis to eliminate many genes from the gonad set that are expressed abundantly by developing embryos, including polyketide synthetase (PKS2), Wnt8, and Endo16e! (Figure 11D).

**Figure 11.**
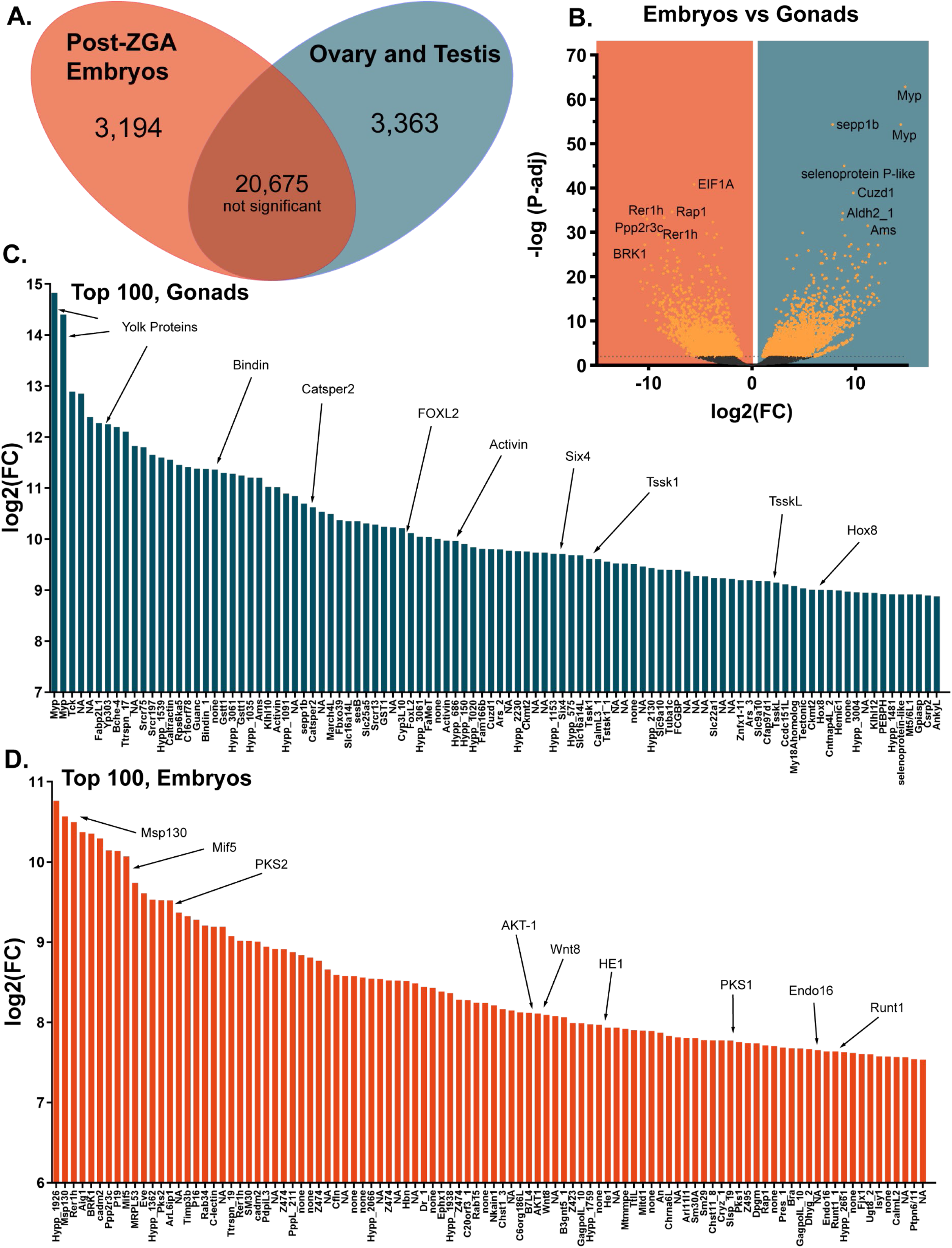
Results of Gonad Hallmarks Analysis. A. Venn diagram summarizing significantly enriched genes and non-significant genes from the 6-way embryos versus gonads analyses. B. Volcano plot of all significantly enriched genes for post-ZGA embryos (left) and gonads of both sex (right) C. Histogram of relative expression for top-100 most significant gonad genes. These were termed “gonad hallmarks” as they represent genes not detected in embryonic development, but rather abundant in gonads of either sex. D. Histogram of relative expression for top-100 most significant embryo-specific genes.

Most interesting were the genes forming the gonad hallmark. Those genes were upregulated in the gonads over post-ZGA embryos in the thousands, and our Deseq2 hallmark analysis returned 3,363 genes highly enriched in gonads. This analysis allowed us to eliminate over 20,675 transcripts shared by both embryos and the gonads (Figure 11A). Surprisingly, FoxL2 was not enriched in nor unique to ovaries as predicted (Ottolenghi et al., 2007; Uhlenhaut et al., 2009), but was very enriched in gonads of both sexes (Figure 9D), while not expressed at all in embryonic and larval stages (Figure 11C). The signaling molecule Activin, which is only first expressed upon the advent of rudiment formation in *Lv* prior to metamorphosis, was also a striking finding (Figure 11C). Lastly, gonad hallmark analysis also yielded the transcription factor, Six4, which has no currently known functional role in echinoderms, but is known to play a role in gonad primordium cell fate in the vertebrate embryo (Lin & Capel, 2015), making it an additional candidate for sex determination and gonadogenesis in echinoderms (Figure 11C). Taken together, these “gonad hallmarks” may each prove to have an essential role in sex determination or gonad function and are tantalizing gene targets for further studies.

### Gonad Hallmarks across developmental time

After performing gonad hallmark analysis, the entire developmental trajectory of Lv, including entire adults and whole gonads, were further analyzed to supplement identified gonad hallmarks with a temporal component. Strikingly, when plotted just by gene expression differences, a clear segregation of transcriptomes based on sex- or gonad-like expression is seen across developmental time (Figure 12A). Gonads occupy the lower right quadrant of the PCA matrix, while embryos develop towards the left quadrant of the PCA matrix with time (Figure 12A). Strikingly, GO term filtering these data over time revealed for the first time, that the expression of many genes involved in gametogenesis begins at the late larval stage (Figure 12B).

**Figure 12.**
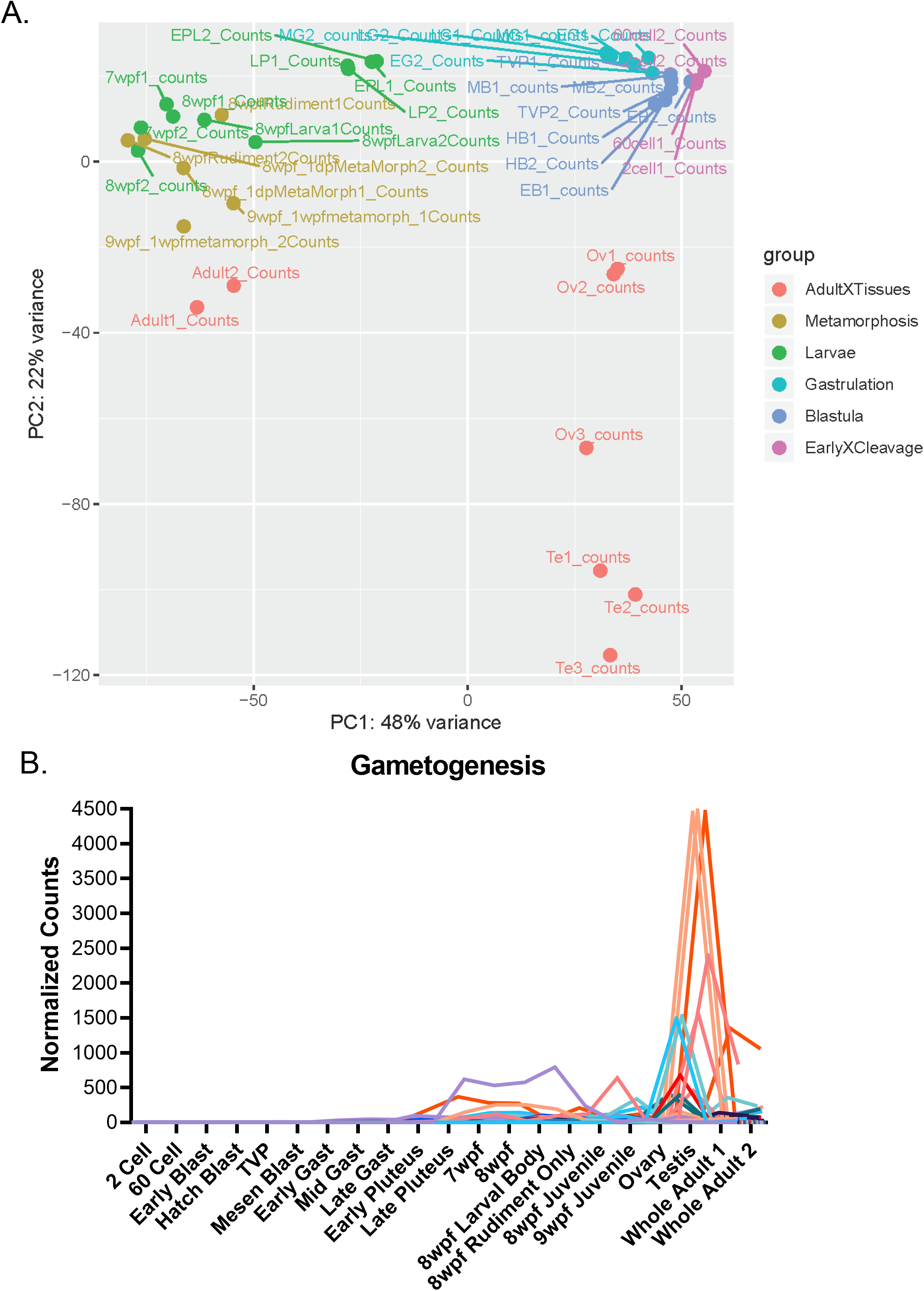
Gonad Hallmark Genes are expressed early in Development. A. Principal component analysis of 42 *L.variegatus* transcriptomes over developmental time, whole adults, as well as gonads. During Deseq2 analysis, transcriptomes were grouped based on transcriptional similarity, groups are denoted with colors. Red are whole adults and gonads, Yellow are metamorphosis and juveniles, Green are late larval growth stages, Cyan are early-late gastrula, Blue are blastula, and Pink are zygote/early cleavage before ZGA. B. Expression graph of GO term = Gametogenesis filtered expression matrix. Genes thought to be unique to gonads in this pathway are detected in late larval development. The developmental timepoints of pluteus-juvenile stages are when small, but detectable amounts of gametogenesis genes are expressed.

### Meiotic genes are active activate well prior to metamorphosis

Meiosis is a unique feature of the germline in reproductive species, regardless of sex (Cleveland, 1947; Darlington & La Cour, 1946). A marker of functional germ cells is whether they can undergo meiosis, an essential process required to produce haploid gametes (Havekes, Jong, & Heyting, 1997), and thus, a “hallmark” of a functioning gonad (Nicholls et al., 2019). To our surprise, meiotic transcripts are active even earlier than those involved in gonadogenesis. Genes such as MeiD, Msx1, and MeioB are observed as early as gastrulation in *Lv* (Figure 13A). To validate sex-specific expression of these genes, in situ hybridizations followed, confirming sex-biasing of meiotic genes such as SYCP3 (Miyamoto et al., 2003) and Msx1 (Simon et al., 1995) among others (Figure 13B).

**Figure 13.**
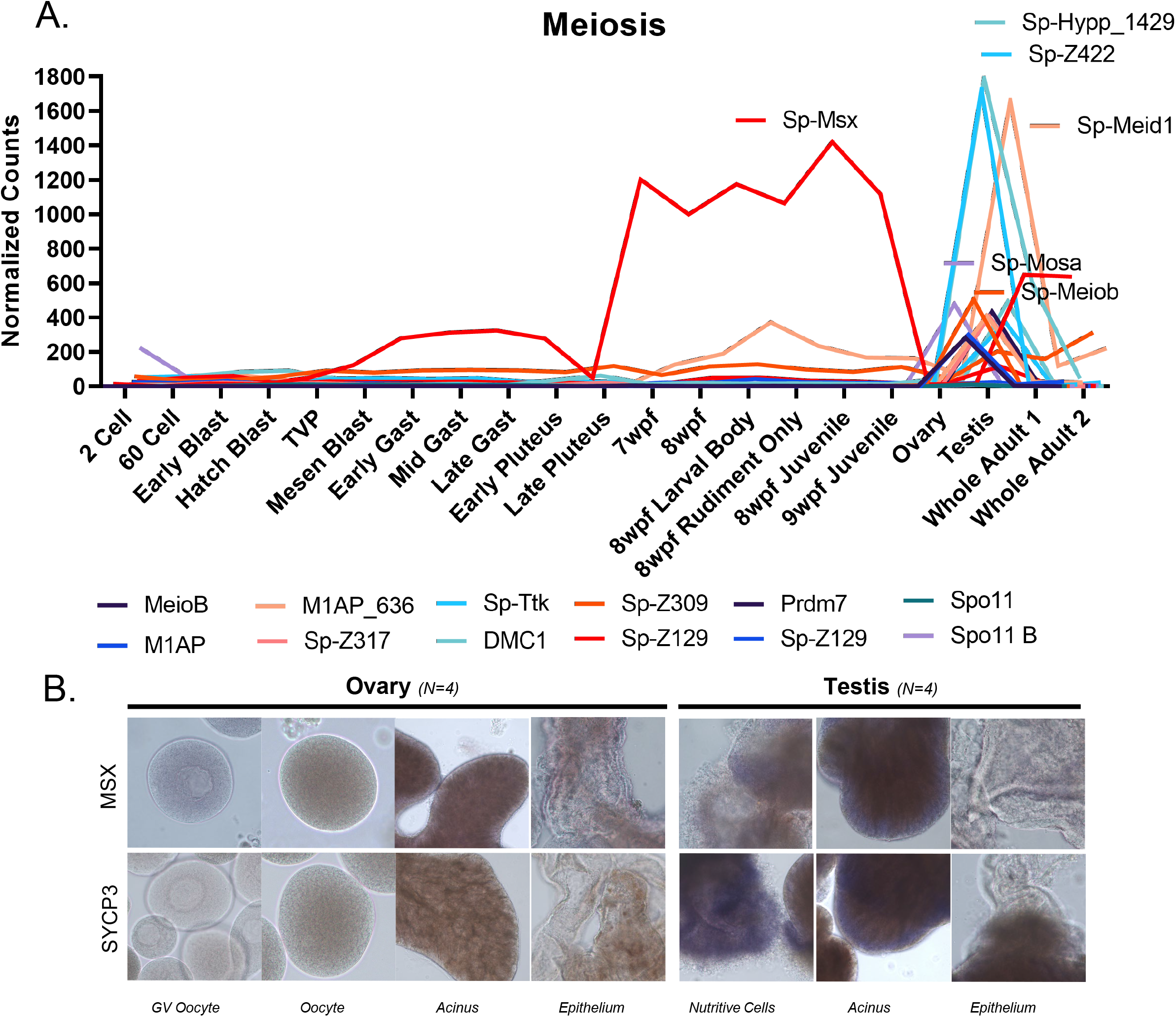
Meiotic Genes are expressed early, and become Male-Specific. A. Meiosis genes expression matrix plotted over developmental time. Many of the genes involved in meiosis are expressed rather early in development, some as early as gastrulation. B. Msx expression in the gonads does not appear to have a strong bias, while SYCP3 appears to have a very strong testis-only signal by whole mount in situ RNA hybridization.

### Sex-determination candidates and gonad hallmarks are expressed prior to metamorphosis

When and where are sex-specific hallmark genes first transcribed? The initial DEG analyses produced a list of gonad-unique transcripts that were unique to each sex, and further gonad hallmark analysis allowed us to narrow them down to those not expressed abundantly by early embryos (Figure10). To supplement the findings of our filtered novel gene lists, we further studied a cohort of sex-determination genes on a candidate-based approach (Figure 14A). SoxH presents a tantalizing male sex determining candidate, as its transcription is activated before even Dmrt genes relative to developmental time (Figure 14A). Most interesting of all, however, is the switching of TGF-B signaling molecules just prior to metamorphosis. From the hallmark analyses, Activin was a candidate female sex determinant, however, when plotted across developmental time with respect to Nodal-a clear binary switch from Nodal-signaling by embryos, to Activin signaling by late larvae just before metamorphosis (Figure 14B).

**Figure 14.**
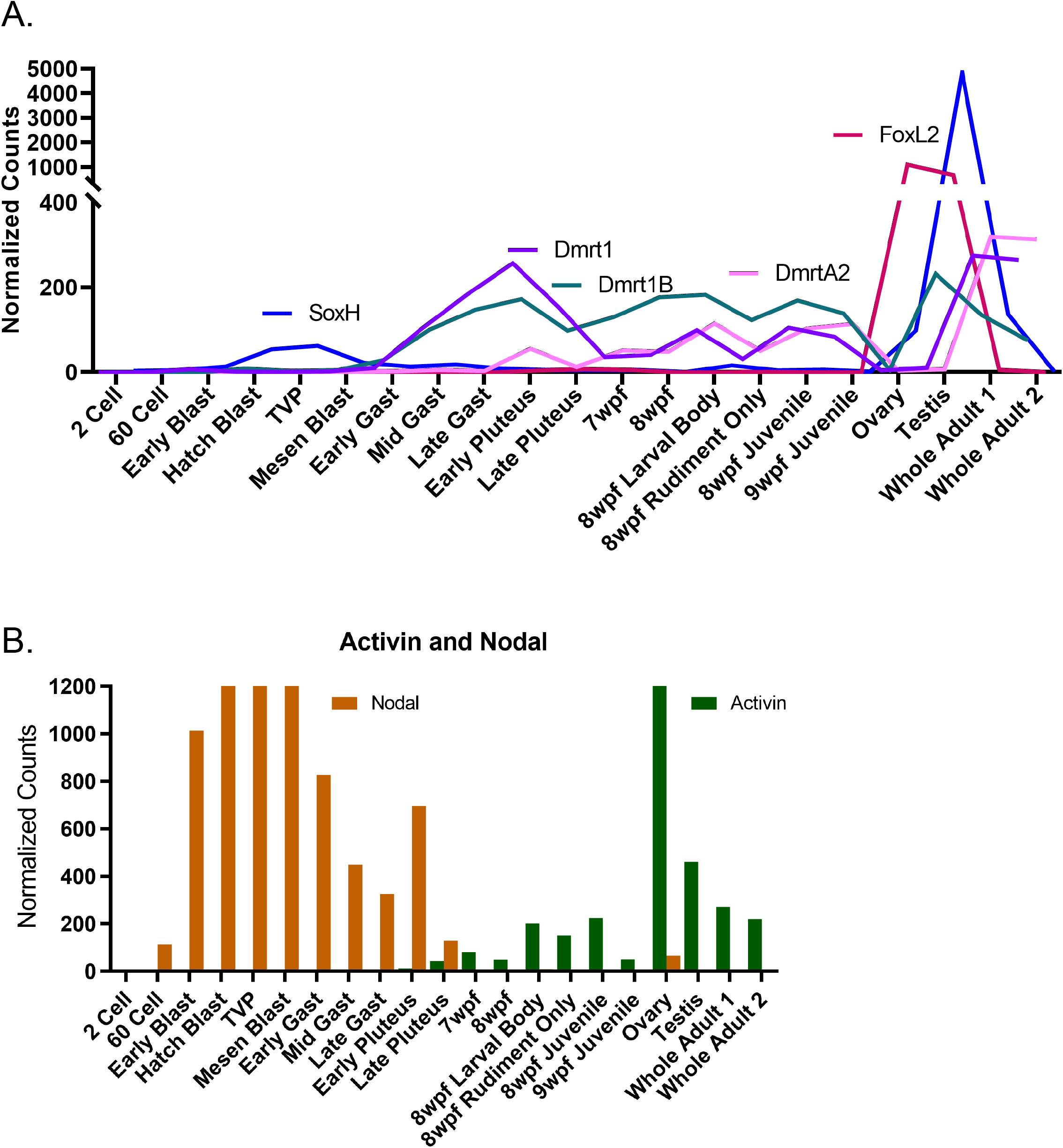
Candidate-Based Sex Determinant Genes. **A.** Gene expression of candidate sex-determinants based on orthology to known vertebrate sex determinants are plotted in developmental time. The majority of our validated candidate genes are those thought to be involved in male sex determination. Note activation of SoxH very early in development, followed by activation of the Dmrt genes. **B.** A histogram of nodal and activin mRNA expression profiles in Lv. Before metamorphosis, expression of nodal diminishes, and activin is expressed, reaching higher and higher abundances until it is most abundantly expressed in the ovary.

As observed with GO filtering for gametogenesis transcripts (Figure 12B), sex-specific markers are transcribed much earlier in development than anticipated. Just as gametogenesis genes activate around late larvae stage, male-specific transcripts appear to activate at the same time (Figure 15A). These transcripts include genes such as Catspers and Tssk’s, thought to be essential only during spermatogenesis, and expressed most abundantly by functional testes in adults (Figure 15A).

**Figure 15.**
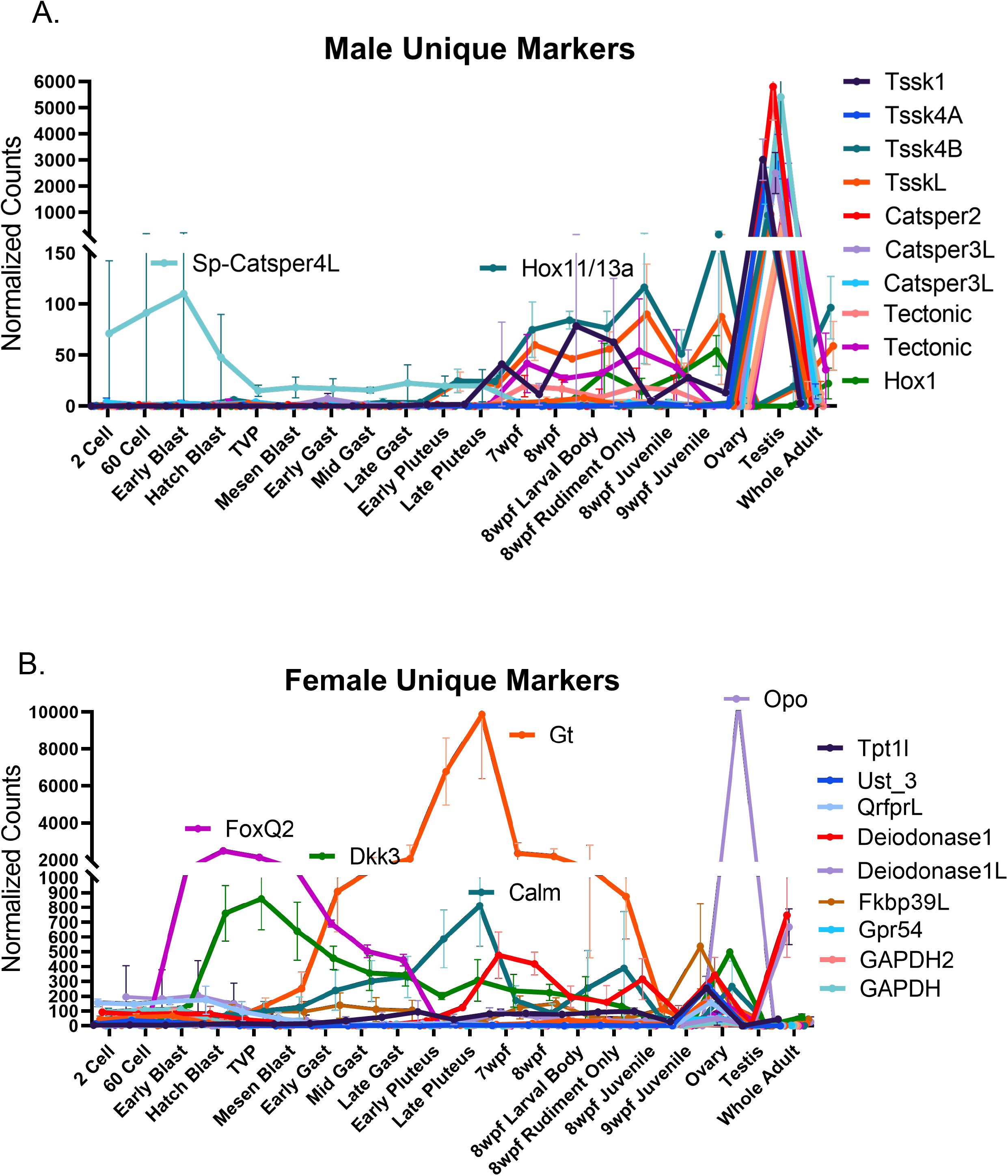
Male- and Female-specific gonad marker genes are expressed early in development. A. A selection of male-specific transcripts from analyses in Fig.1 are plotted here across developmental time from the analyses in Fig.12. Testis transcripts are generally detectable at late larval developmental stages, prior to metamorphosis, apart from the calcium ion channel Catsper-4L which appears to be maternally loaded. Error bars represent standard error, and all values were significantly enriched (not detected in ovary samples). B. A selection of female-specific transcripts from analyses in Fig.1 are plotted here across developmental time from the analyses in Fig.12. Ovary-specific transcripts are not only detectable at late larval development, but many of them also appear to be expressed rather early in development at blastula and gastrula stages.

More striking are the female-specific genes plotted across developmental time. When trying to parse out germline-oocyte-versus somatic transcripts in the female, most transcripts are those used by the oocyte as it develops. Because of the overlap between oocyte and female sex specific transcripts, many transcripts returned during our initial DEG analyses were lost when filtered for those not used in oocytes, and still more filtered out during gonad hallmark analysis. For ease of plotting, the top female-specific transcripts are simply plotted here (Figure 15B). Many of these ovarian genes, including calmodulin and dickkopf-4 (Tulac et al., 2006; J. Wang, Shou, & Chen, 2000), are expressed very early in development, around gastrulation, or earlier (Figure 15B). In line with male sex candidates, however, Gt, Deiodonases, and calmodulin genes are transcribed in similar abundance at the late larval/pre-metamorphic stages of development (Figure 15B).

A group of sex determination candidates is the Doublesex gene family. Doublesex (Dmrt) genes include a battery of DNA-binding or splice-isoform regulating master switches of sex determination in nearly any species of metazoan examined (Camara, Whitworth, Dove, & Van Doren, 2019; Jeng et al., 2019) While the means of transcription and basis of function of Dmrt genes are variable-one factor remains the same: their role in male sex identity (Matsushita, Oshima, & Nakamura, 2007; Tang et al., 2019; Webster et al., 2017). *Lytechinus variegatus* has five Doublesex genes present in its genome. Of these five genes, one encodes a gonad-specific transcript with almost no known functional domains, thus named: doublesex-like (DsxL). Two more encode the homologs of Sp Dmrt: Dmrt1 and Dmrt1B, and interestingly, both are implicated in early embryonic development as well as testis function. It is likely that Dmrt1B is a duplication of Dmrt1 due to sequence similarity, yet Dmrt1B encodes a transcript with unique Ankyrin repeat domains (Figure 16A). DmrtA2 represents a novel Dmrt gene, it shares sequence identity with other Dmrt genes, however it encodes a transcript with nuclear hormone receptor domains. Interestingly, DmrtA2 is only expressed during embryonic and larval development, and is not expressed in the testes of the adult organism (Figure 16B). Finally, Dmrt2 is transcribed at incredibly low abundance, and its function remains rather elusive. Dmrt2 in Lv has two bona fide DNA-binding functional domains like that of vertebrate doublesex, as well as a DM domain like Dmrt1 and Dmrt1B (Figure 15A). We confirmed that Dmrt1 and DsxL are expressed in the testes, while DmrtA2 appears to be exclusive to embryos (data not shown). The expression of the doublesex genes in gonads support a male-specific role for Doublesex genes in sea urchins, though their functional role in sex determination, whether early or late, warrants further studies.

**Figure 16.**
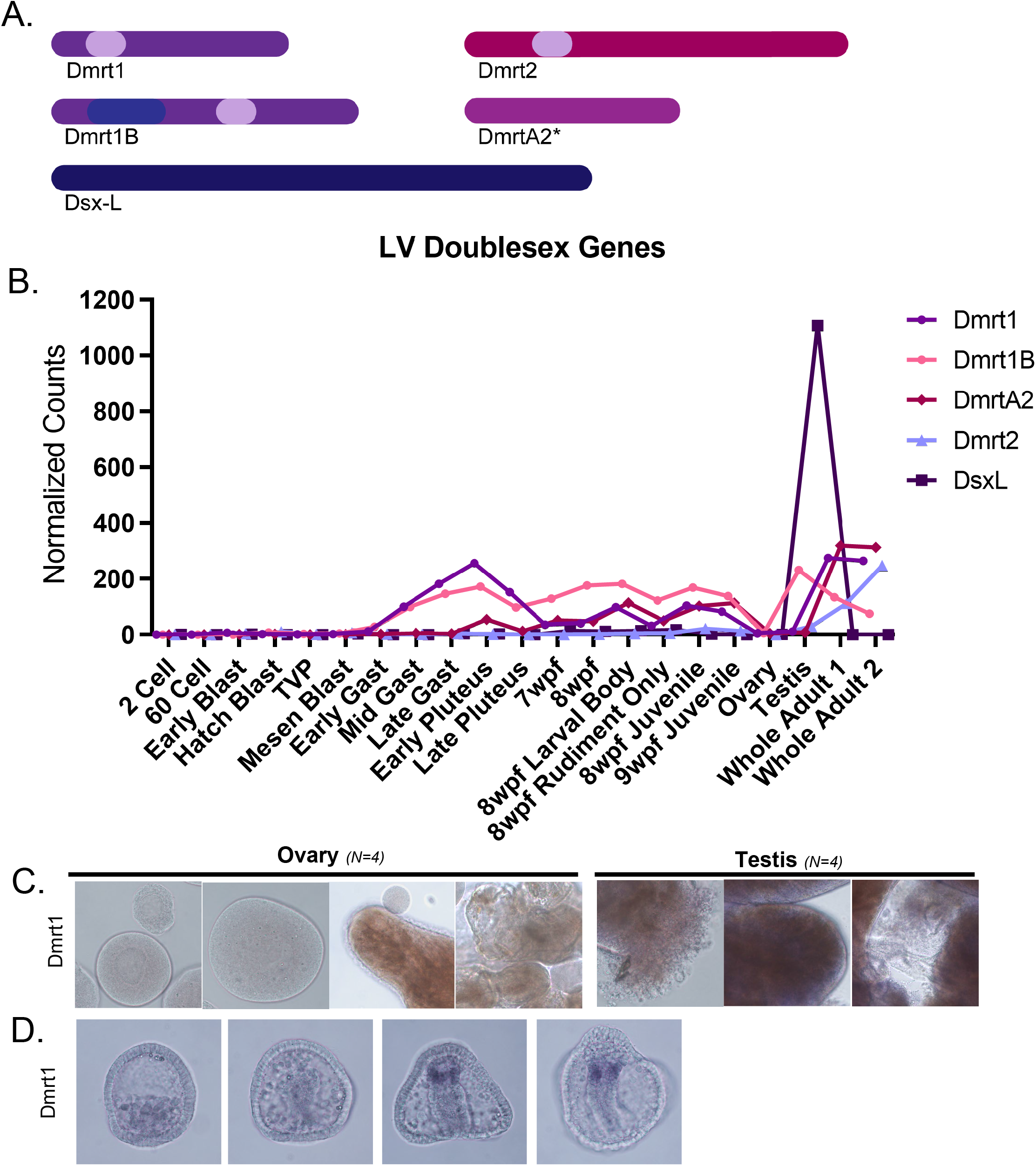
Five Lv Doublesex Genes and their Expression Patterns. A. Outline of the five Dmrt transcripts identified and characterized in Lv. The colored bars represent peptide length, and light-colored regions represent predicted DM domains. Dmrt1B’s predicted peptide has an ankyrin repeat domain (purple) not observed in the other Dmrt1 gene. B. Summary and expression graph for the five Doublesex gene orthologs in *L.variegatus*. Each doublesex transcript has a unique expression pattern across developmental time. DmrtA2 and Dmrt1 are not expressed abundantly in gonads of either sex, but are expressed abundantly, transiently early in development at the gastrula stage. Doublesex-Like (DsxL), however, appears to be a testis-only transcript not expressed embryonically. C. Validation of Doublesex gene expression by whole-mount in situ in gonads D. In situ RNA hybridization of Dmrt1 in embryos shows very clear and spatially restricted expression at late gastrula stage.

### Individual late-stage larvae express sex-specific transcripts

Following the whole transcriptome-based analysis and identification of novel transcripts implicated in gonad function and sex biology, we obtained a shortlist of possible gene targets implicated in sex determination or gonadogenesis. Because so many were expressed prior to metamorphosis (Figures 12-16) we sought to test whether the larvae of Lv urchins do express sex-specific mRNAs. For these analyses, individual larvae leading up to, and just prior to metamorphosis were individually assayed via qPCR for the expression of sex-specific mRNAs. The gene expression results varied widely depending on the stage in development of the larvae in question.

For analysis of the individual larvae, we selected male- or female-specific transcripts with no question of their role in gonad function. Strikingly, there emerge two patterns of temporal expression: first, and more abundantly, male; followed by later, and in lower abundance, female sex marker expression (Figure 17C). In 8-armed larvae with no large rudiment present, individual Lv express sex specific transcripts in one of three patterns: a low but consistent number of male-specific transcripts, a composite of male-with some female-transcripts, or no sex specific transcripts at all (Figure 17A). Later in development, at the competent rudiment stage, however, individual larvae express much more abundant sex-specific transcripts with a clear biasing towards one sex profile: male, or expression of sex-specific transcripts of both sexes-(Figure 17B). When plotted using multiple regression, it is more obvious that these larvae follow distinct gene expression profiles at these stages (Figure 18A&B). Further, it can be noted that the tendency to express “male with some female,” transcripts, or “neither” is characteristic of late larvae, while sex-specific gene expression or male-only sex transcript expression is a characteristic of those with a competent rudiment (Figure 17D).

**Figure 17.**
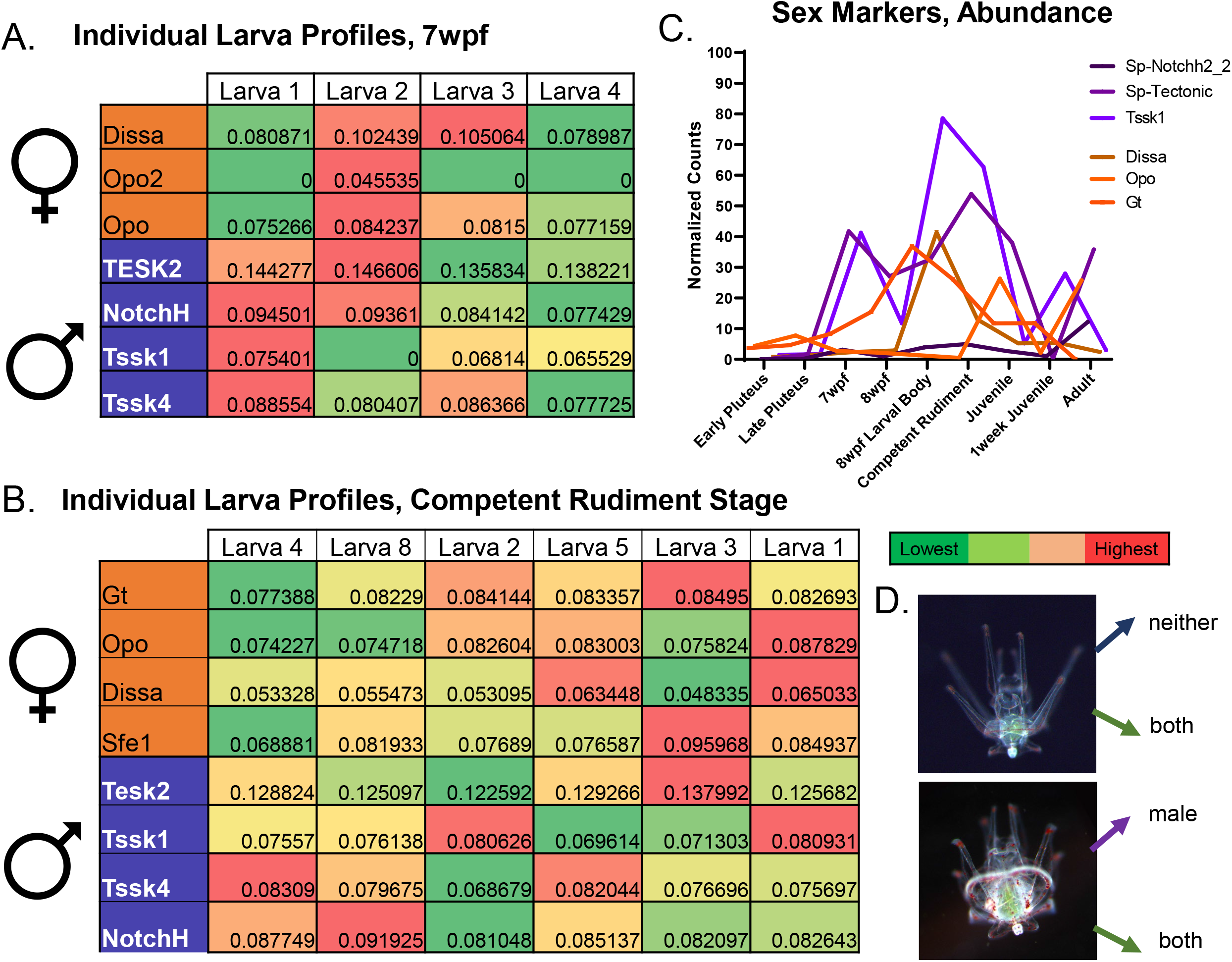
Individual larvae express sex-specific transcripts before metamorphosis. A. Heatmap summarizing individual larvae qPCR data at the late larval (8 arm) stage. A key for heatmap colors can be found below and to the right side. Each column represents and individual larvae, and each row represents a male- or female-specific transcript. Some individual larvae at the 8-arm stage show biased male-only gene expression, while others express both male- and female-markers, or do not express sex-specific markers in detectable amounts. Numbers represent normalized expression values for each gene. B. Heatmap summarizing individual larvae qPCR data for competent rudiment stage larvae. The sex-specific transcripts become far more abundant at this stage in larval development, before metamorphosis. C. Summary of the expression of the gene markers used for this analysis. qPCR primers were generated and validated for each of these genes. Note that female sex markers are expressed just after, and at lower abundance than the male sex markers at this stage of development. D. Summary of gene expression findings from individual larva qPCR. At the 8 arm stage, larvae express one of two profiles: neither, or male with some female gene expression as well (both). At the competent rudiment stage, larvae express male-only sex profiles, or a collection of both sex marker genes.

**Figure 18.**
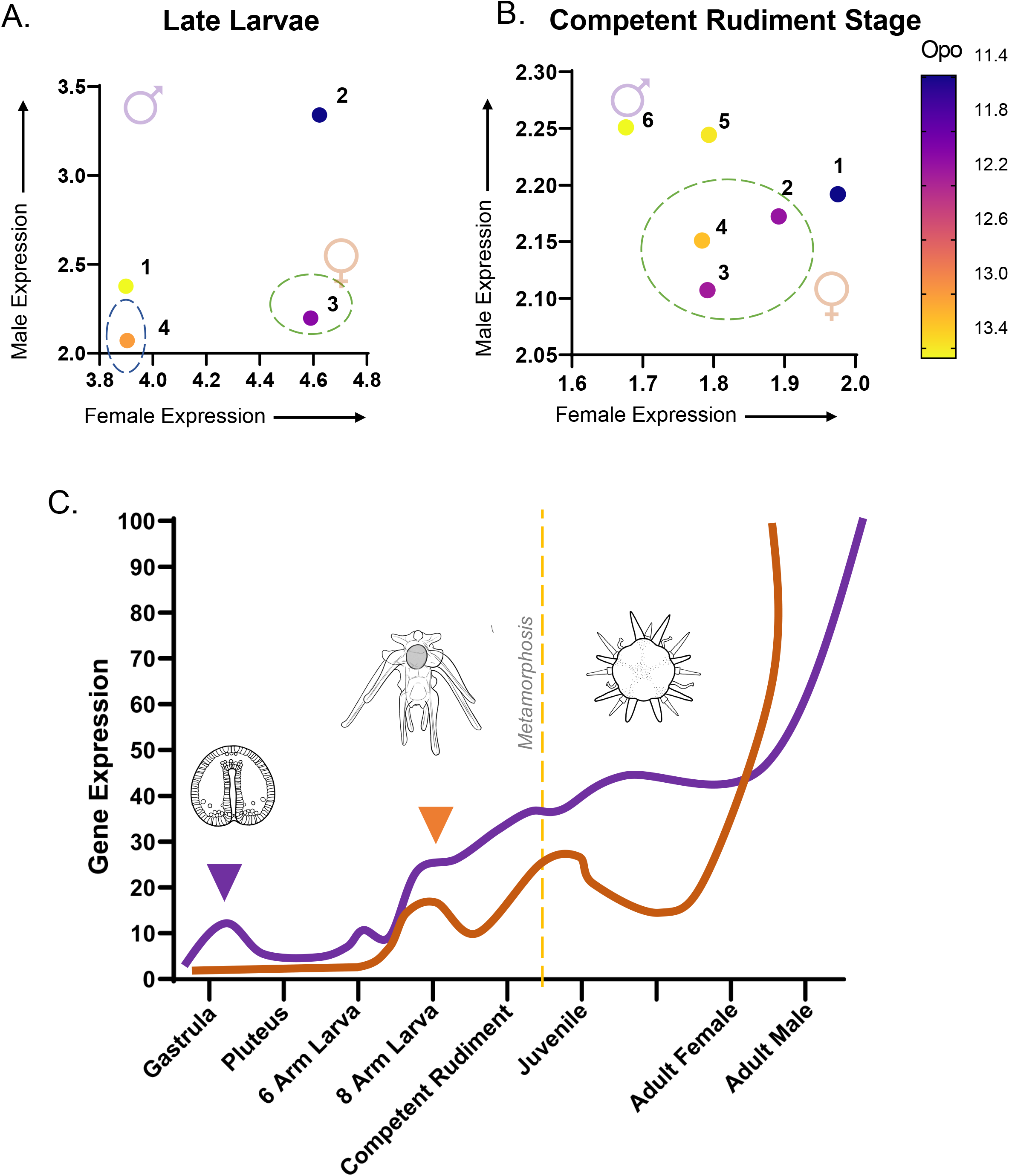
A New Hypothesis for Sea Urchin Sex Determination. A. Multiple regression bubble plot of gene expression matrices from Figure 17 at the 8 arm stage. With female gene expression (arbitrary units, averages of the group of genes) on the X-axis, a clear distinction is made where larvae 1 and 4 express neither sex profiles (dark blue circle), and larvae 3 expresses both sets of sex markers (green circle) B. Bubble plot at the competent rudiment stage demonstrates the marked increase in sex marker expression at this stage in development. Importantly, 3 out of 6 larvae cluster in the center of the plot, indicating a gene expression profile where both male- and female-markers are expressed. Importantly, at this stage, a clear delineation of male only sex marker expression becomes apparent as in larvae 5 and 6. C. Summary of findings. A hypothesis for expression of sex determination genes in the sea urchin. Before larval growth, male candidate sex determinants are expressed (purple arrowhead, see SoxH and Dmrt genes). Later, before metamorphosis, there is an activation of male sex markers first, closely followed by a marked increase of female sex marker expression (orange arrowhead). The expression of male sex markers becomes clearly delineated just before metamorphosis (dotted line). We do not know the trajectory of female sexual differentiation, but following metamorphosis to a reproductive adult, the differentiation into a functional ovary likely takes place.

## Discussion

We propose a model for the transcriptional activation of sea urchin sex determination pathways (Figure 19C). First, even before larval growth and development, we observed the activation of possible male sex determinants such as SoxH or the Dmrt genes. Next, at late larvae stage, before metamorphosis, we observed activation and relatively stable expression of male-specific sex transcripts. This was closely followed with shaky expression of some female transcripts as late larvae and then as competent rudiment stage larvae. Prior to metamorphosis, we observed that many individual larvae still express both male- and female-specific transcripts, leading us to wonder whether an activation of the female pathway and differentiation into an ovary occurs later in development, and if this is a product of inhibition by the continued, or stabilized expression of the male pathway. If there is indeed a bipotential gonad in this species of sea urchin, it is the deliberate switching of gene expression by activation of the male pathway that appears to be the ticket to sexual differentiation.

As observed in other organisms, maintenance of glycolytic and gluconeogenic pathways are essential for the development of healthy oocytes (Sutton-McDowall et al., 2010) It is known that in vertebrates, it is an essential function of the soma of the ovary to transfer sufficient glycolytic products to developing oocytes (Ou et al., 2012; Q. Wang et al., 2009) Our findings support a hypothesis that it is the sheer volume of oocytes produced by the female sea urchin that warrant the expression of a unique GAPDH transcript for ovaries in order to provide sufficient glycolytic activity (Biggers et al., 1967). Additionally, the sea urchin egg is a giant, developmentally primed cell, virtually bursting with mitochondria. Therefore, it may be that the production of sufficiently mitochondria-loaded eggs is the reason for a glycolysis reliant ovary in this animal (Jansen & de Boer, 1998). Because it is already established that glycolytic products are transferred to the oocyte itself in mammalian oogenesis, further studies seek to characterize, and establish, exactly which supporting cells within echinoderm ovaries are responsible for the high glycolytic activity, and further, how these products of glycolysis are transferred to maturing oocytes.

An important observation is the wide variation observed in the expression of germline, oocyte, and somatic sex genes in the ovaries of this species of sea urchin. While the testis whole transcriptome profiles cluster rather closely (Figures 5&6), the ovaries remain widely variable, with one entire transcriptome clustering somewhere between the collection of ovary and testis transcriptomes (Figure 12). While it cannot be explained as the sole reason for this variation, it can be noted that depending on the gravidity of ovaries (Stage I-IV), the proportion of oocytes to somatic cells, as well as yolk content appears to be highly variable. To this end, we have supplied representative images of Stage III oocytes, those most like the gonads used for these analyses (Supplemental Figure 1).

PGCs are an essential unit of reproduction, and their fate following embryogenesis is crucial to study, as we still do not know whether sea urchins have a bona fide bipotential gonad, nor if the PGCs themselves have any influence on the fate of the bipotential primordium. It is known in other organisms, that germline genes are often under sex-specific control (Camara et al., 2019; Zhao, Svingen, Ng, & Koopman, 2015) or, as in zebrafish-the germ cells may even influence the sex identity of the somatic gonad itself (Dranow, Tucker, & Draper, 2013; Uchida, Yamashita, Kitano, & Iguchi, 2002). Upon assaying germline genes, we parsed out very sex-specific expression patterns of critical germline genes such as Nanos and Pumilio in ovaries. It is still unknown if sex differentiation in the sea urchin is what influences the expression of these genes, or vice-versa. Perhaps most interestingly of all, we found and characterized nine nanos transcripts unique to *Lv*. Their existence, especially that the majority of them are Nanos3 homologs expressed by the testes, implies a strange role of a novel transposase, perhaps in maintenance of spermatogonial stem cells.

A bipotential primordium is a unique and common feature of embryos across many phyla (Nef et al., 2019), with conserved gene programs directing this bipotential gonad down a trajectory of differentiation into one sex or another (Garcia-Moreno, Plebanek, & Capel, 2018; Peng et al., 2021; Rey et al., 2000). Taken with our new findings in *Lv* larvae, this implies a shared mechanism of gonad development, or deeply conserved mechanism in which the gonad primordium is influenced by a balance of sex determination markers, almost as if it senses the outcome of earlier gene expression, and ultimately responds, albeit slowly, to the developmental tilting towards one sex or another. If so, one can now imagine how slight changes in gene expression earlier in development may influence the balance and change the outcome. One might also imagine how environmental factors, whether they be physical or biological, can also influence a gradual shifting of the gonad’s outcome.

While the complex pathways governing sex determination in this animal have yet to be dissected, important new findings are presented here: the male fate factors Dmrt and SoxH are expressed early, meiosis initiates early, and gonad-specific transcripts involved in egg and sperm biology are first activated before rudiment formation in the larvae of this sea urchin. Based on previous findings in other organisms (Bagheri-Fam et al., 2010; Tang et al., 2019), a common thread through sex determination pathways is the mutual agonism-and competitive antagonism of sex identity by the male pathway (Ottolenghi et al., 2007; Rey et al., 2000). By finding clean distinction of male sex genes in the late larvae of *Lv*, we have new support for a hypothesis of genetic sex determination mechanism in the sea urchin. To dissect this pathway in further detail, we propose to use CRISPR-Cas9 generation of mutant juvenile *Lv* sea urchins. We know now it is possible to validate and assay individual larvae for the expression of male- or female-specific transcripts, but we now must follow mutant animals through juvenile growth to their adult life stage, when they produce functional gametes. An essential consideration is whether there is full sex reversal, or a partial sex reversal following the knockout of sex-determination candidates early in development. Additionally, it should be noted that cases of hermaphroditic individuals in *Lv* and closely related gonochoristic sea urchins are incredibly rare, based on scant reports (Boolootian & Moore, 1959). Twins raised from the same zygote (separated as blastomeres) develop into adults of the same sex (Cameron, Leahy, & Davidson, 1996) arguing strongly for a genetic basis of sex determination. Our findings provide additional evidence for a genetic mechanism.

## Materials and Methods

### Gonad Collection and RNA-seq

Freshly collected Lv were sexed by opening the test. Ovaries and testis were identified, rinsed briefly in filtered seawater, and immediately lysed by douncing in Trizol. The samples were then frozen at −80C and sent on dry ice for library construction and sequencing (Novogene).

### Gonad DEG Analyses

Each sequencing file as a FASTQ.gz was downloaded via FTP client to usegalaxy.org for subsequent DEG analyses. First, FASTQC, and Trimmomatic! Were used to prune low quality reads and adapter sequences from each FASTQ file (Bolger, Lohse, & Usadel, 2014; Chen et al., 2017)For alignments, HiSAT2 (Kim, Langmead, & Salzberg, 2015) used to align reads to the Lvar_Scaffolds.fasta genome assembly (Davidson et al., 2020)Following alignments, featurecounts was used to generate gene count tables for each alignment file. A proprietary gene annotation file: modified_GREG.GTF generated by our lab was used as the transcript annotation reference, it may be found in the supplemental data.

To grant further confidence to our DEG analyses, resultant count files were also run through the EdgeR platform.Finally, the original FASTQ files were also concurrently analyzed using a classic RNA-seq pipeline which utilizes tophat (Trapnell, Pachter, & Salzberg, 2009), bowtie (Langmead, Trapnell, Pop, & Salzberg, 2009) and cuffdiff (Trapnell et al., 2010) to produce DEG analyses without the use of count tables. These results are summarized in supplemental figure 3.

### Novel Transcript Identification

Following DEG analyses, fifty gonad-enriched or gonad-sex-specific transcripts with no predicted name or orthology were selected based on both transcript abundance and those with lowest P-adj values. To characterize each transcript, mRNA was translated into peptides using Expasy translate, and protein structure predictions were then generated using interproscan-version 5.45-80.0 software (Jones et al., 2014). A comprehensive list of L_Var ID’s with their associated protein structure prediction file can be found in the supplemental data.

### Evolutionary analyses of *L.variegatus* nanos orthologs

Ten nanos transcripts were returned from gonad germline gene expression analyses. All ten transcripts were compared to the three characterized nanos orthologs in *S.purpuratus*, first using PRALINE multiple sequence alignment to determine the peptide identity, and further using Clustal OMEGA and phylogenetic tree analysis (http://www.phylogeny.fr/documentation.cgi). For all ten nanos transcripts, genomic location and position was identified using the genome build from (Davidson et al., 2020) and local blast on BioEdit software. Chromosome maps were reconstructed from chromosome maps provided in (Davidson et al., 2020).

### Lv Gonad Hallmarks Analysis

All additional developmental timepoint .FastQ.gz files were accessed from the RNA seq study published in (Hogan et al., 2020; Li et al., 2020) which may be accessed at: https://www.ebi.ac.uk/ena/browser/view/PRJNA554218. To generate a DEG analysis and expression matrix, DEseq2 software was once again used, following the same pipeline as above (Liu et al., 2021; Love et al., 2014). Whole transcriptomes were compared across two groups “gonads”, versus “post-ZGA” to display only those transcripts unique to, or most abundantly expressed in the gonads (3,363), and eliminate those transcripts shared or unique to the post-ZGA embryos (>23,000 transcripts)

### LV Developmental Timepoint Analyses

All additional developmental timepoints as FastQ.gz files were accessed from the RNA seq study published in (Hogan et al., 2020; Li et al., 2020) which may be accessed at: https://www.ebi.ac.uk/ena/browser/view/PRJNA554218. To generate an across-development matrix for referencing transcript abundance normalized across all timepoints, DEseq2 software was once again used, following the same pipeline as above (Liu et al., 2021; Love et al., 2014). Differential gene expression analyses were not pursued, rather individualized groups across developmental time were compared broadly. These groups are herein denoted as “early cleavage” which corresponds to zygotes and pre-ZGA embryos, “blastula” which includes all hatched, swimming, and mesenchymal blastula stage embryos, “gastrulation” which includes gastrula stage embryos as they progress through gastrulation, “larvae” which includes the pluteus through late larval stage embryos, “metamorphosis” which includes competent rudiment stage and young juveniles, and finally “adult tissues” which includes two entire small adult *Lv* urchins as well as the gonads. DESEQ2 outputs were three normalized count tables, standard, VST-normalized, and log2 normalized, of all gene expression across development.

### Primer design

For in-situ hybridization analyses, transcripts from *Lv* transcriptome from (Davidson et al., 2020)were selected and two sets of primers designed for each using NCBI PrimerBLAST software. All PCR products selected range between 700-1000bp. A T7 promoter is added to the reverse primer to generate antisense probes. For qPCR, transcripts were used as template in Primer3Plus software to generate two sets of primers for each transcript of interest. A comprehensive list of primers used in this publication may be found in the supplemental data.

### RNA In-situ Hybridization of Gonads and Embryos

All RNA in-situ hybridizations were performed as previously published (Arenas-Mena, Martinez, Cameron, & Davidson, 1998; Perillo, Paganos, Spurrell, Arnone, & Wessel, 2021). For whole gonad in situs referenced throughout the paper, respective negative controls may be found in the supplement (Supplemental Figure 1).

### Culture and individual qPCR of late-stage larvae

Lv larvae are cultured on benchtop aquaria at room temperature (25-27deg C) in 2uM filtered MBL water with a simple paddle rotator. Cultures are fed Filter Feeder Formula from Algae Research & Supply Company (https://algaeresearchsupply.com/) in mornings until satiated on odd days, with water change following feeding. Once reaching desired developmental timepoint, larvae were staged using darkfield microscopy to identify rudiment size and rudiment progression. Individual larvae were collected in PCR tubes and lysed in 250uL RLT buffer, then stored at −80deg until qPCR date. For qPCR analyses, individual larvae lysis were prepped using the Qiagen RNeasy micro kit (Qiagen, Cat#74004) and eluted in exactly 14uL RNAse free H2O. The 14uL RNA eluate is then immediately used for cDNA synthesis using Maxima First Strand cDNA synthesis kit (ThermoFisher **Cat#** K1641). cDNA is diluted 1:4 and used for subsequent qPCR analysis in triplicate.

## Supporting information

Pieplow Supplemental data

---

**Supplementary Figure 1.**
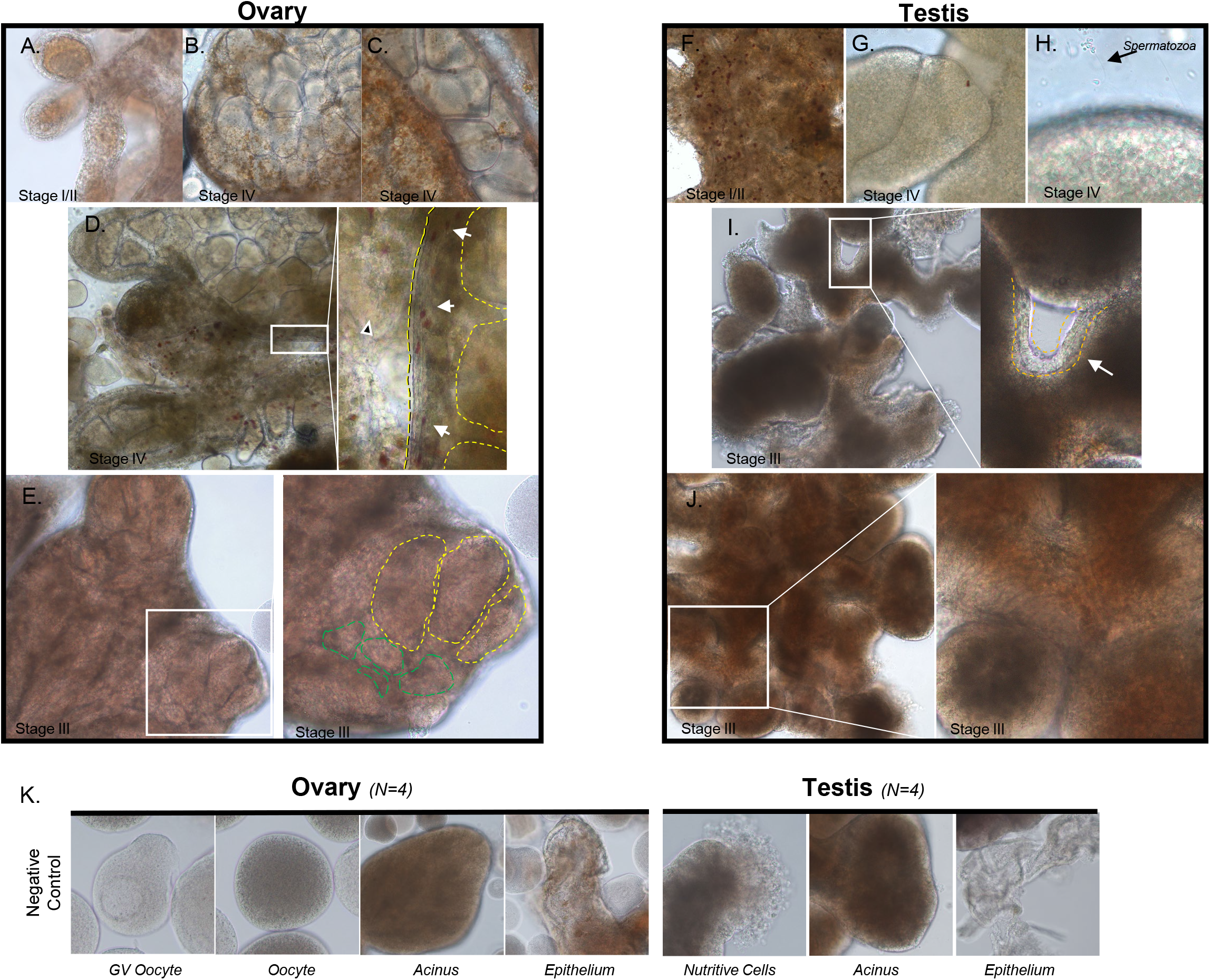
Gonad Images. A. Stage I/II ovary example, ovaries in this condition were not harvested for analysis. B. Mature, gravid, stage IV ovary example, note presence of large amounts of mature oocytes. C. Mature, gravid, stage IV ovary example, closeup of mature oocytes visible in a passageway between lobes. D. Light microscopy of ovarian structures, black arrowhead = nutritive cells, white arrows = colored phagocytes, yellow outlines = ovarian epithelium housing mature oocytes surrounded by nutritive cells. inset =10x Magnfication E. A Stage III ovary example, inset = immature oocytes in various stages. F. Stage I/II Testis Example G. Mature, Gravid, Stage IV Testis example image H. Live imaging inset shows swimming spermatozoa which are released upon agitation I. Stage III testis with inset showing epithelium (yellow) J. Structures of a Stage III testis include coiled tubules, epithelium and interstitial spaces filled with yolk and nutritive cells.

**Supplementary Figure 2.**
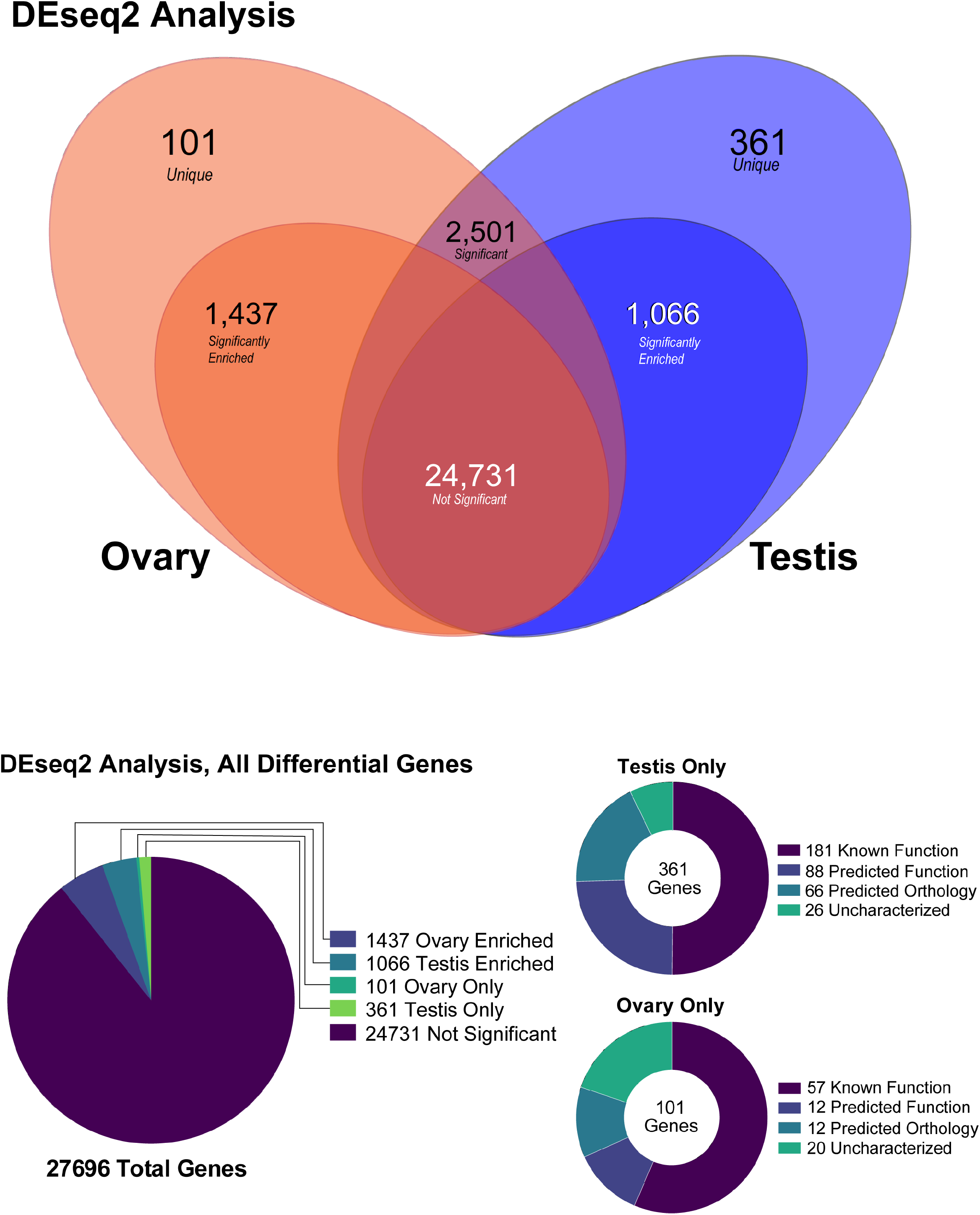
DeSeq2 Supplementary Results. A. Venn Diagram summarizing all genes from this analysis B. Pie Chart of all significant or unique genes as a fraction of the total C. Pie Chart of testis only genes by annotation status D. Pie Chart of ovary only genes by annotation status

**Supplementary Figure 3.**
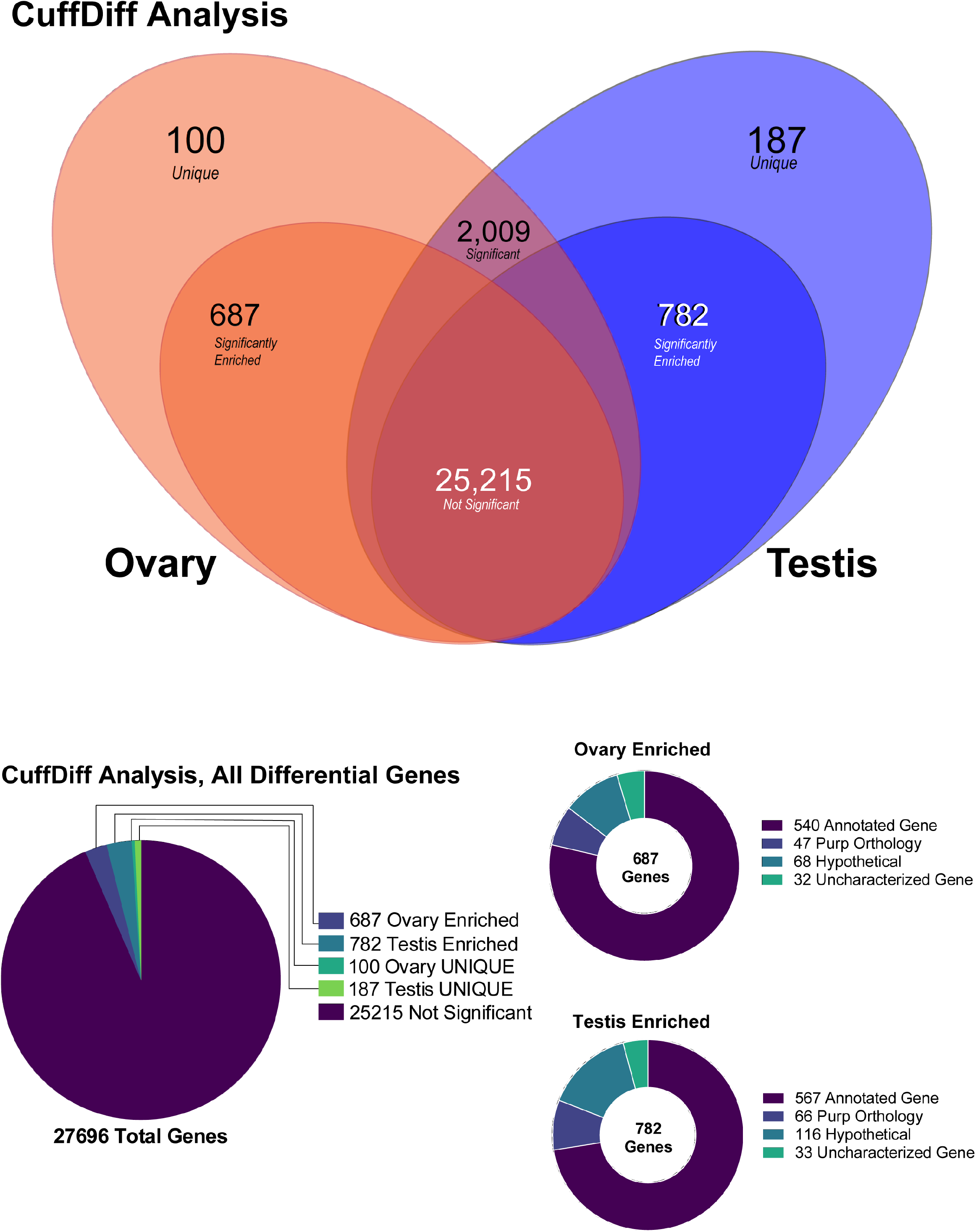
CuffDiff Supplementary Results. A. Venn Diagram summarizing all genes from this analysis B. Pie Chart of all genes as a fraction of the total C. Pie Chart of ovary enriched genes by annotation status D. Pie Chart of testis enriched genes by annotation status

**Supplementary Figure 4.**
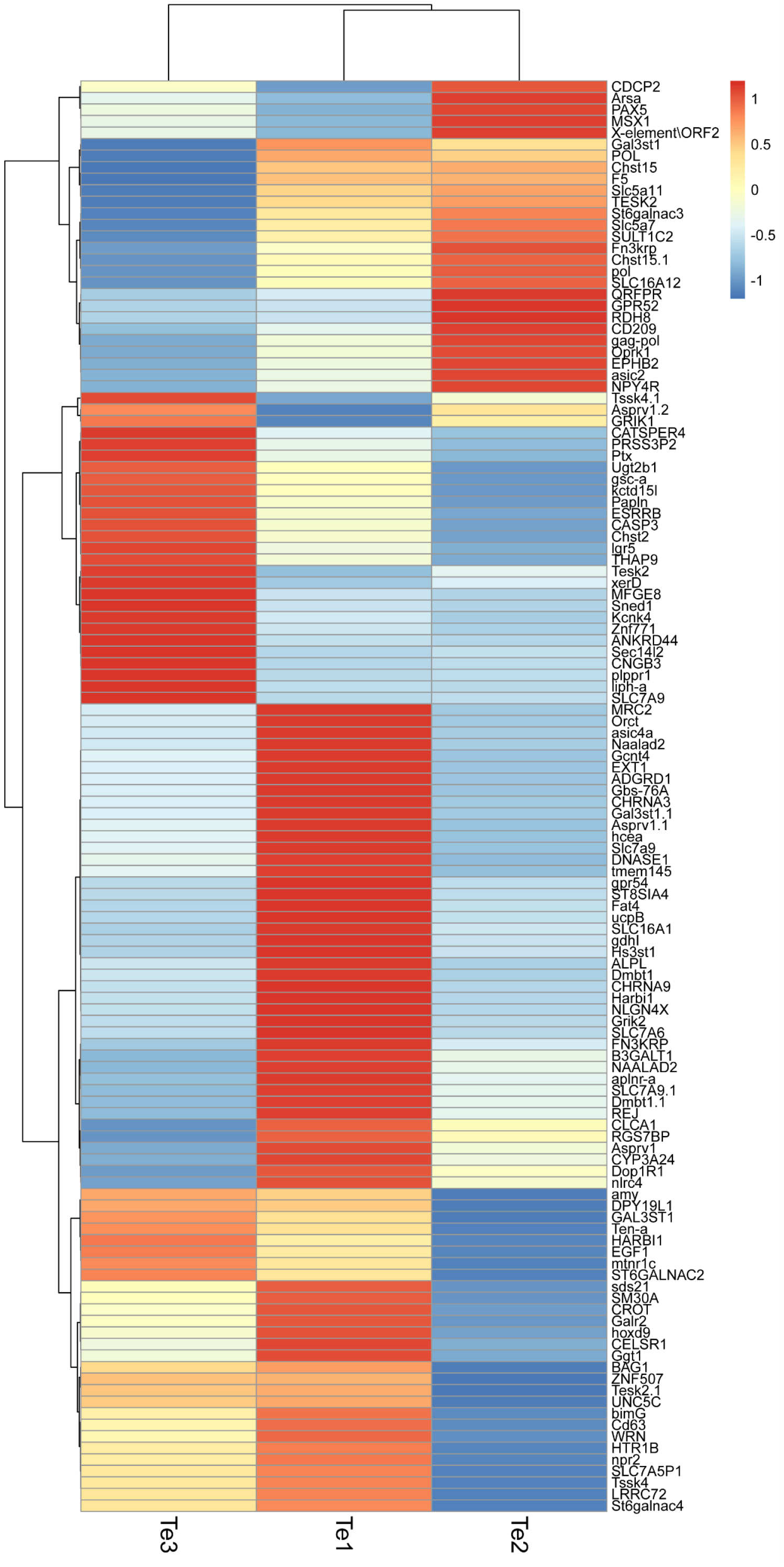
Male Specific Genes are expressed variably across individual gonads. Heatmap matrix of expression of testis-specific transcripts across the three testes replicates. Male specific genes identified through the pipeline in Fig.1 are not expressed in the same amount by all three testis replicates assayed in this study. There is a great variability in “testis-specific” gene expression by individual gonads.

**Supplementary Figure 5.**
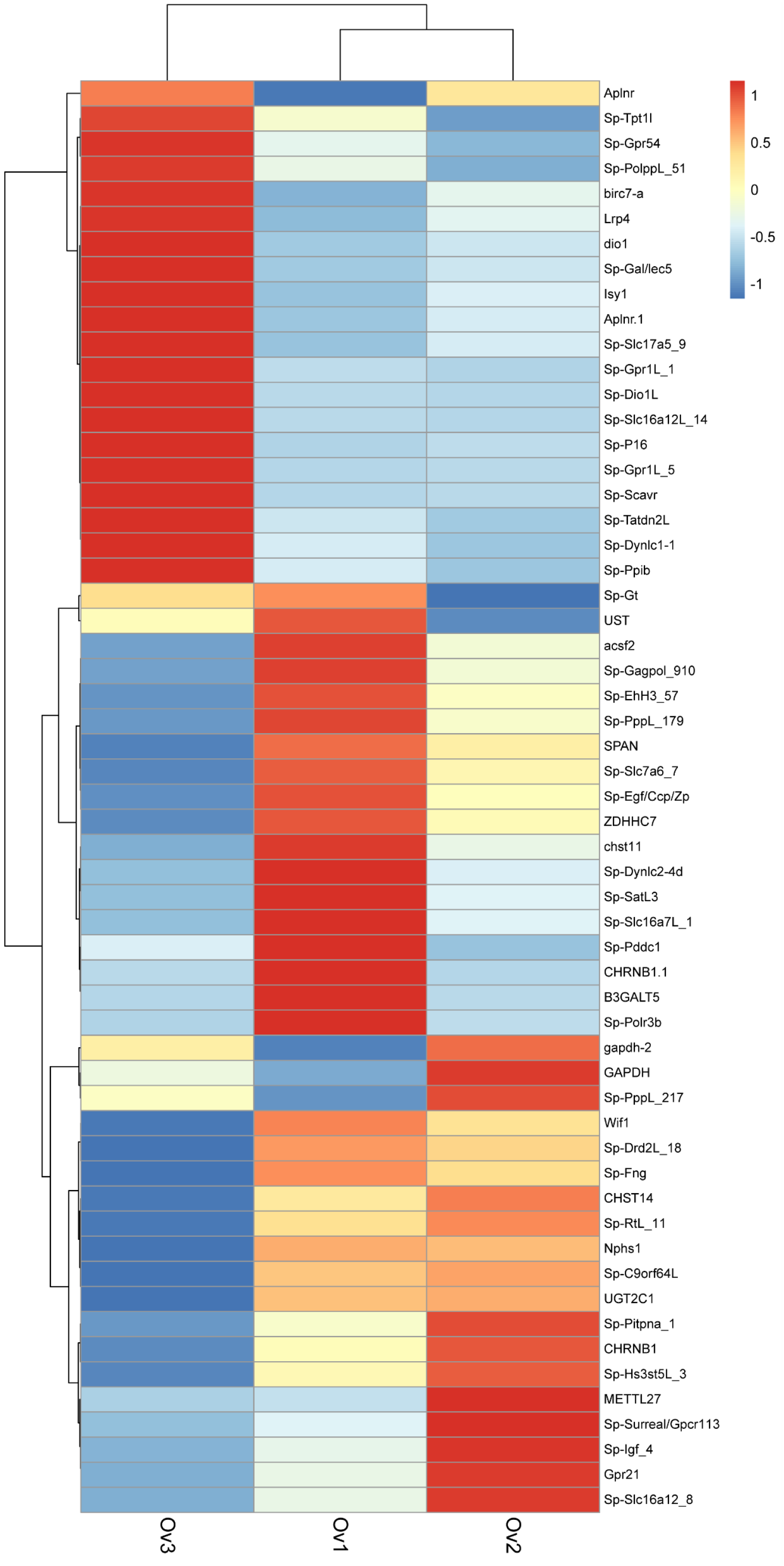
Female Specific Genes are expressed variably across individual gonads. Ovary specific genes identified through the pipeline in Fig.1 are not expressed at the same levels by all three ovary replicates assayed in this study. There is a great variability in “Ovary-specific” gene expression by individual gonads.

**Supplementary Figure 6.**
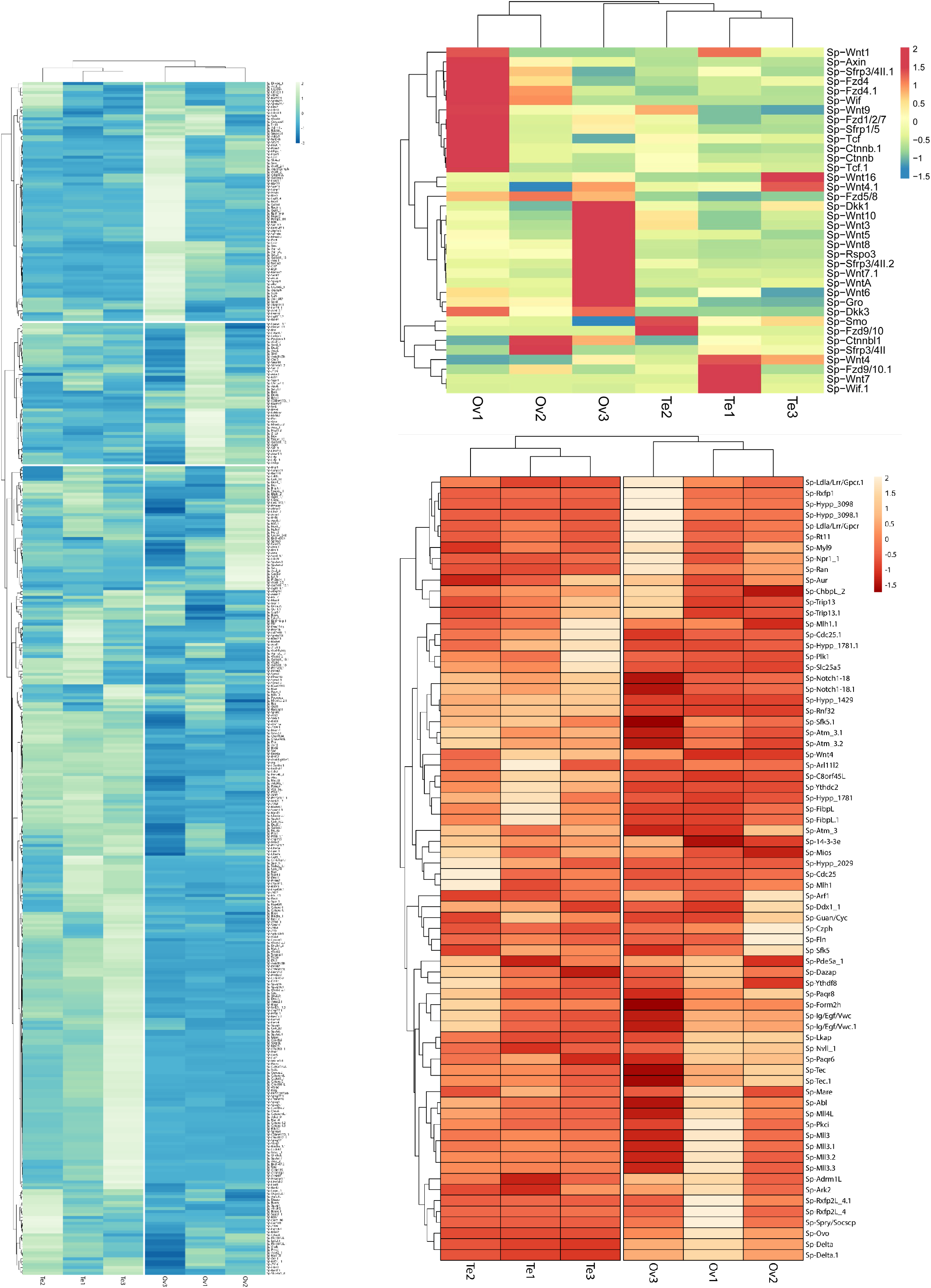
Heatmaps of Expression by Sex, gene groups. A. Spermatogenesis B. Wnt Signalling C. Oogenesis

**Supplementary Figure 7.**
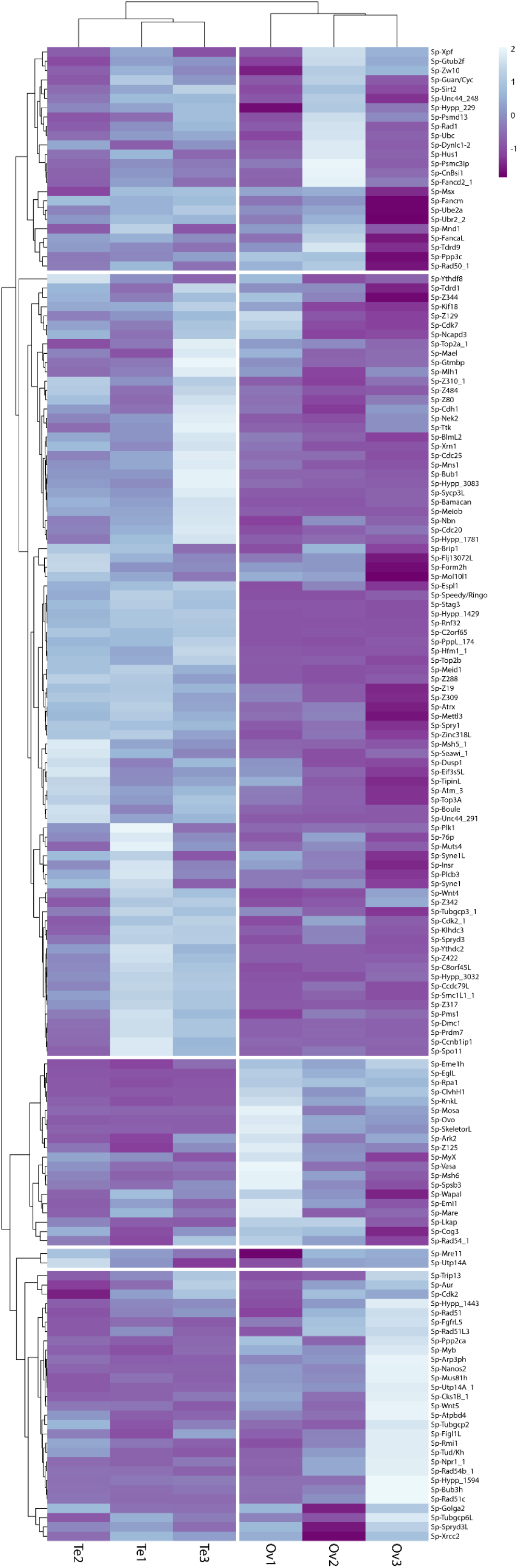
Meiotic gene expression by Sex. Heatmap of all GOterm= meiosis genes expression levels by gonads

**Supplementary Figure 8.**
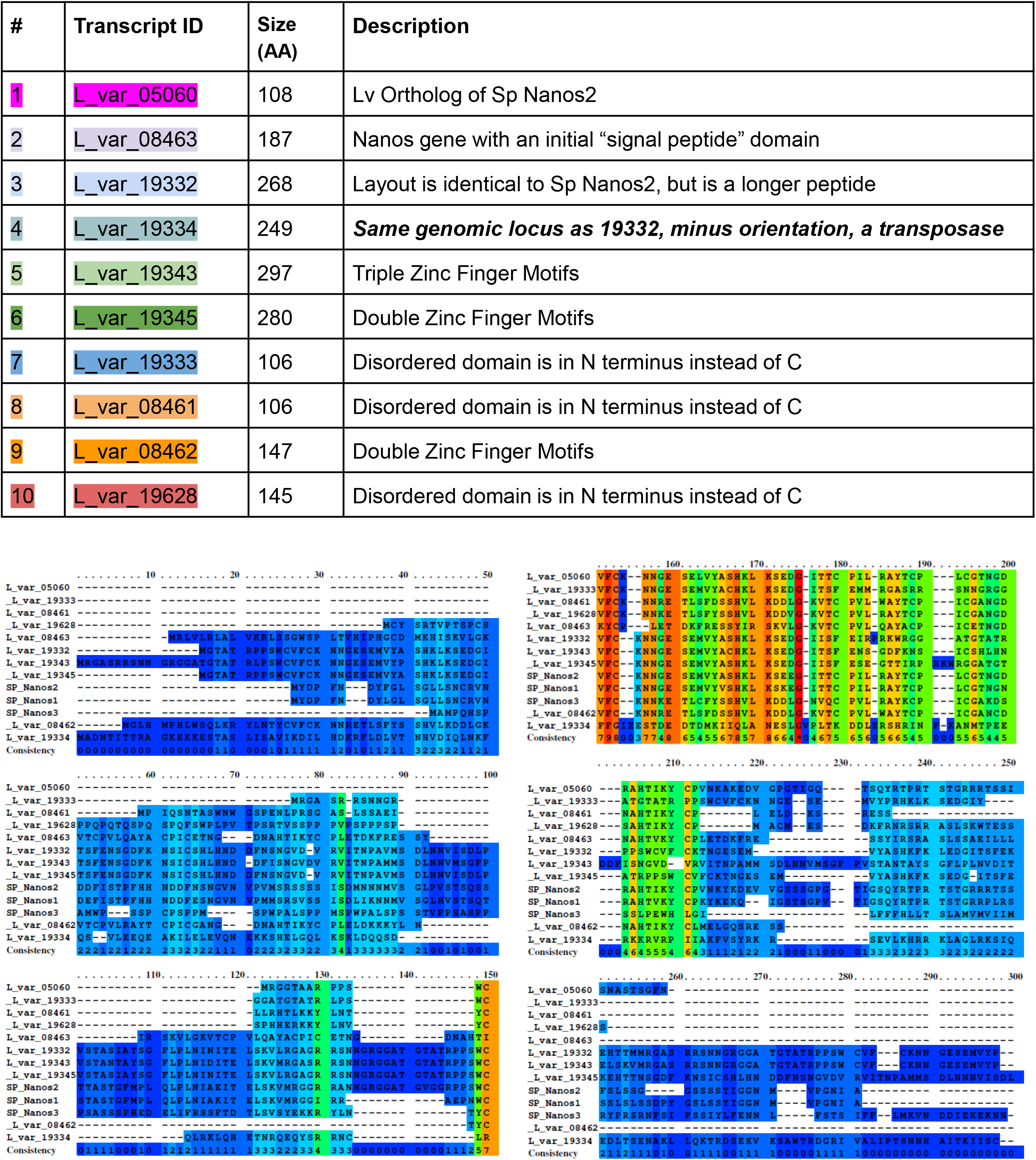
Summary of Nanos Genes. A. Table of all nanos transcripts identified in this RNA-seq approach, note that one is occupying the same genomic locus as L_var_19334, but is transcribed antisense and produces a transposase. B. Peptides from each nanos transcript were compared with peptides for each respective nanos homolog in *S.purpuratus. Alignments were generated using PRALINE MSA software, color represents amino acid identity*.

**Supplementary Figure 9.**
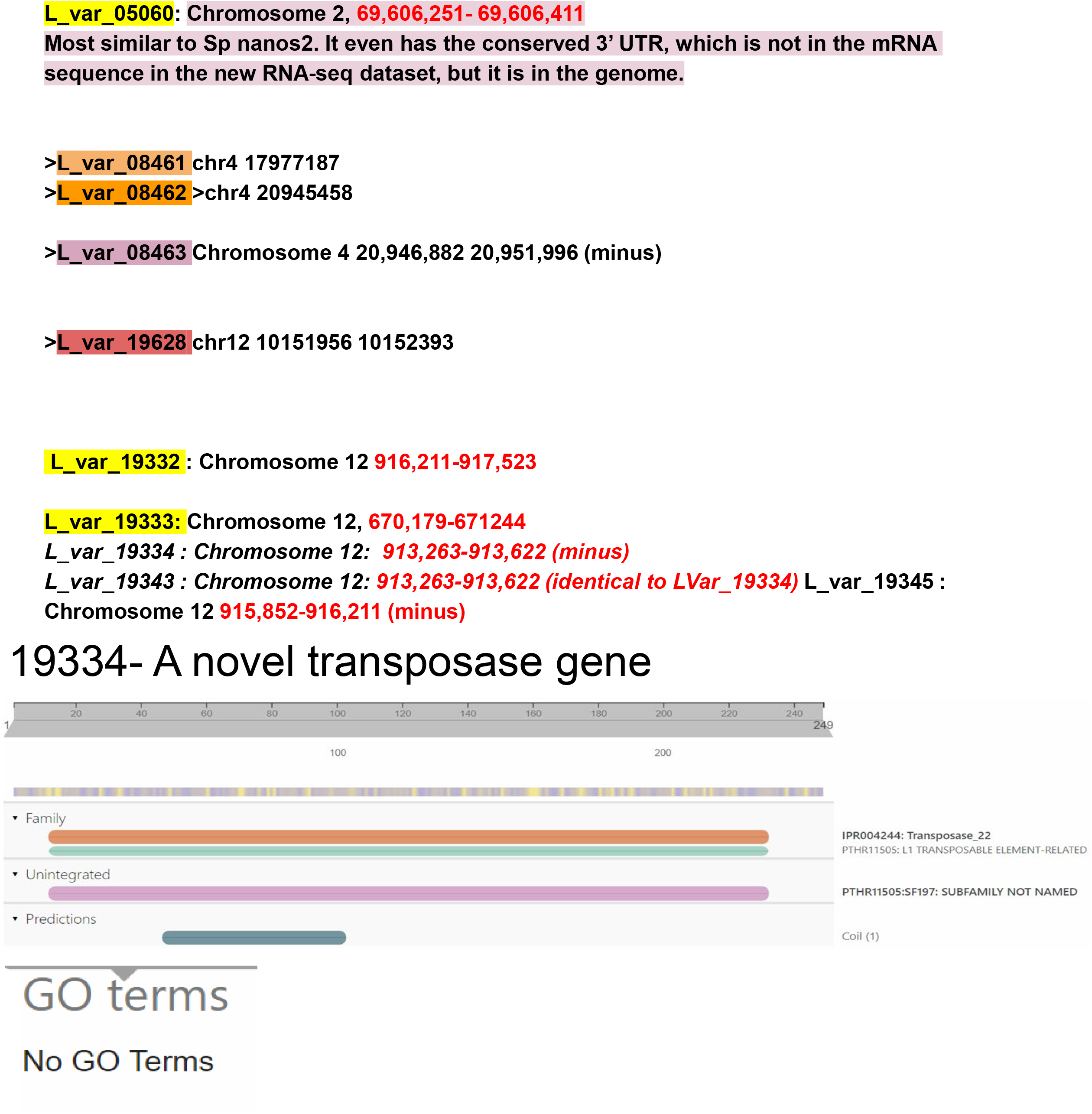
Nanos genes occupy unique genomic loci. A. Genomic loci for each nanos transcript in *L.variegatus*, used to generate positional information on chromosome maps. B. Structural information for Lv transcript 19334, which encodes a novel transposase. Note that there is no associated GO with this type of transposase.

**Supplementary Figure 10.**
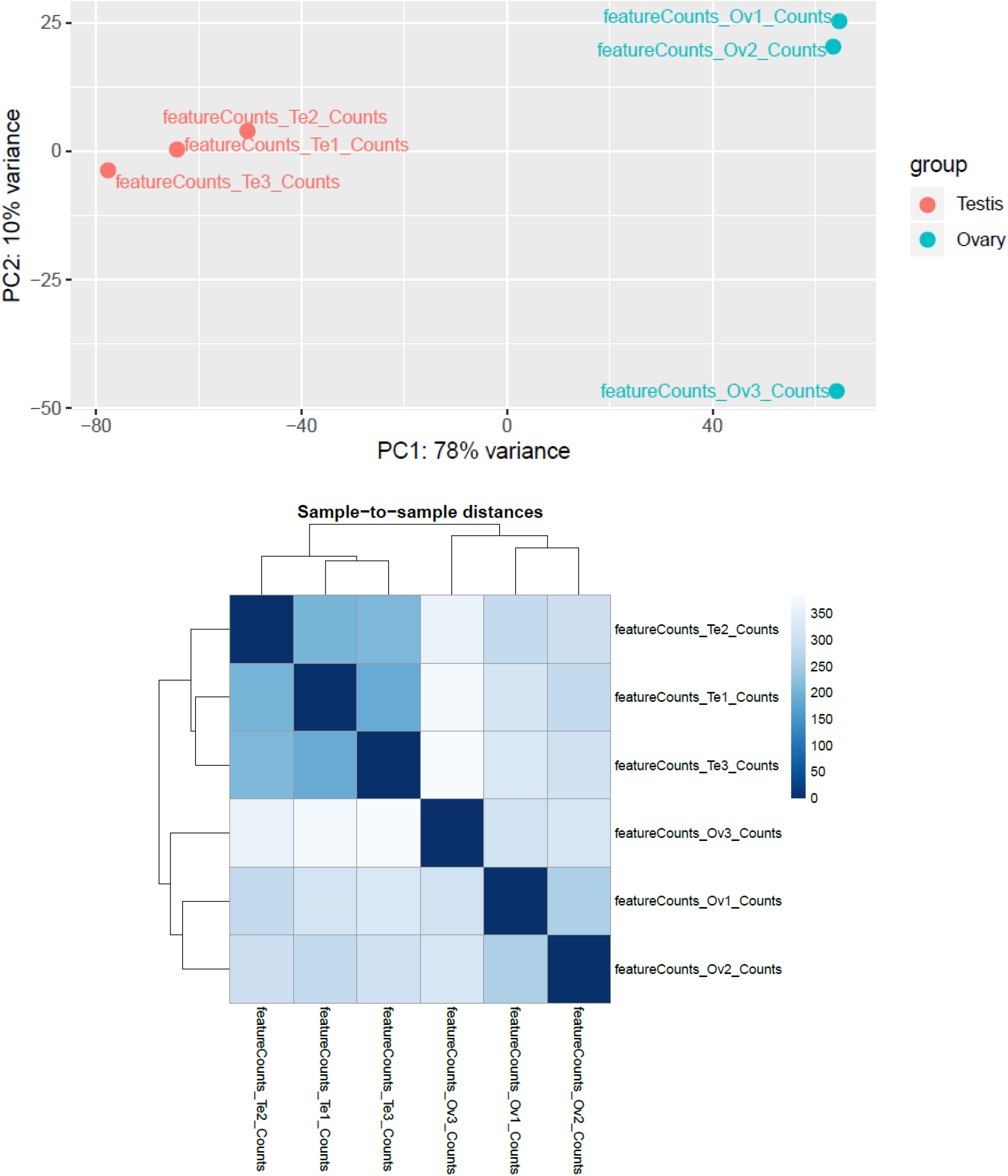
OvvTe Supplemental Information 1. A. PCA plot B. Matrix of sample-sample mean distances

**Supplementary Figure 11.**
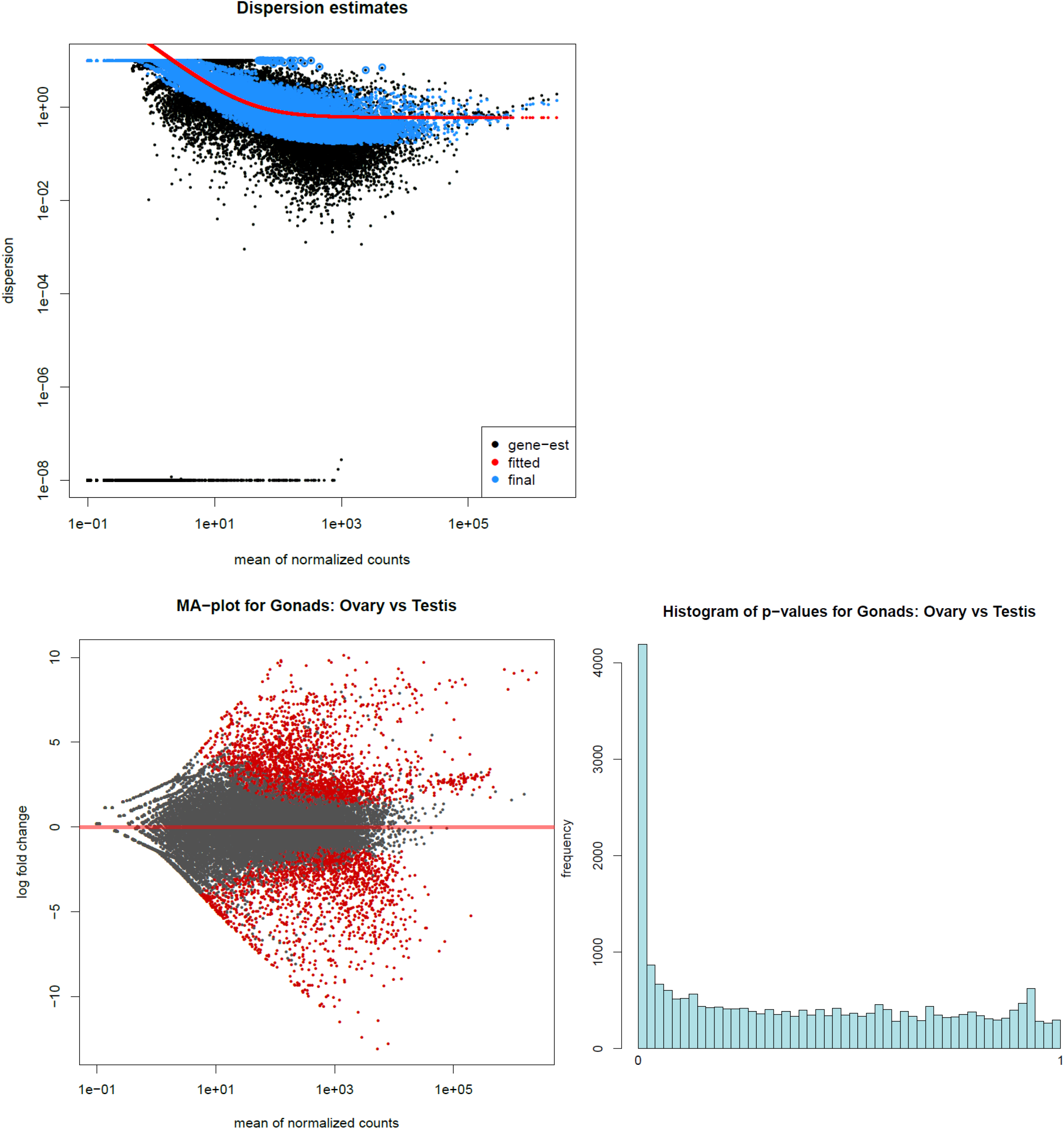
OvvTe Supplemental Information 2. A. Dispersion estimates for Deseq2 data B. MA-plot of mean relative to log-fold change values C. Histogram of p-value distributions for all data

**Supplementary Figure 12.**
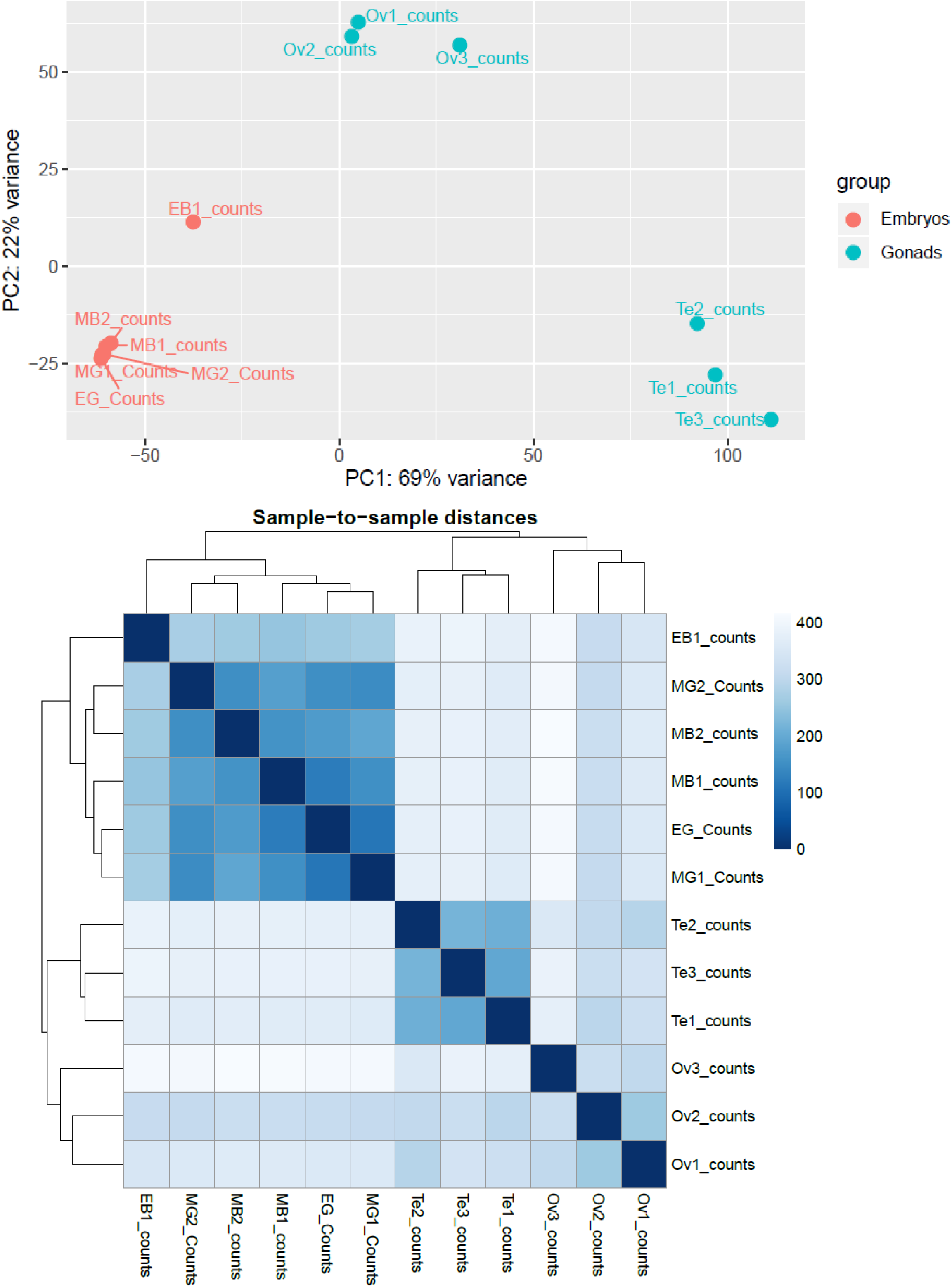
6-way analysis Supplemental Information 1. A. PCA plot B. Matrix of sample-sample mean distances

**Supplementary Figure 13.**
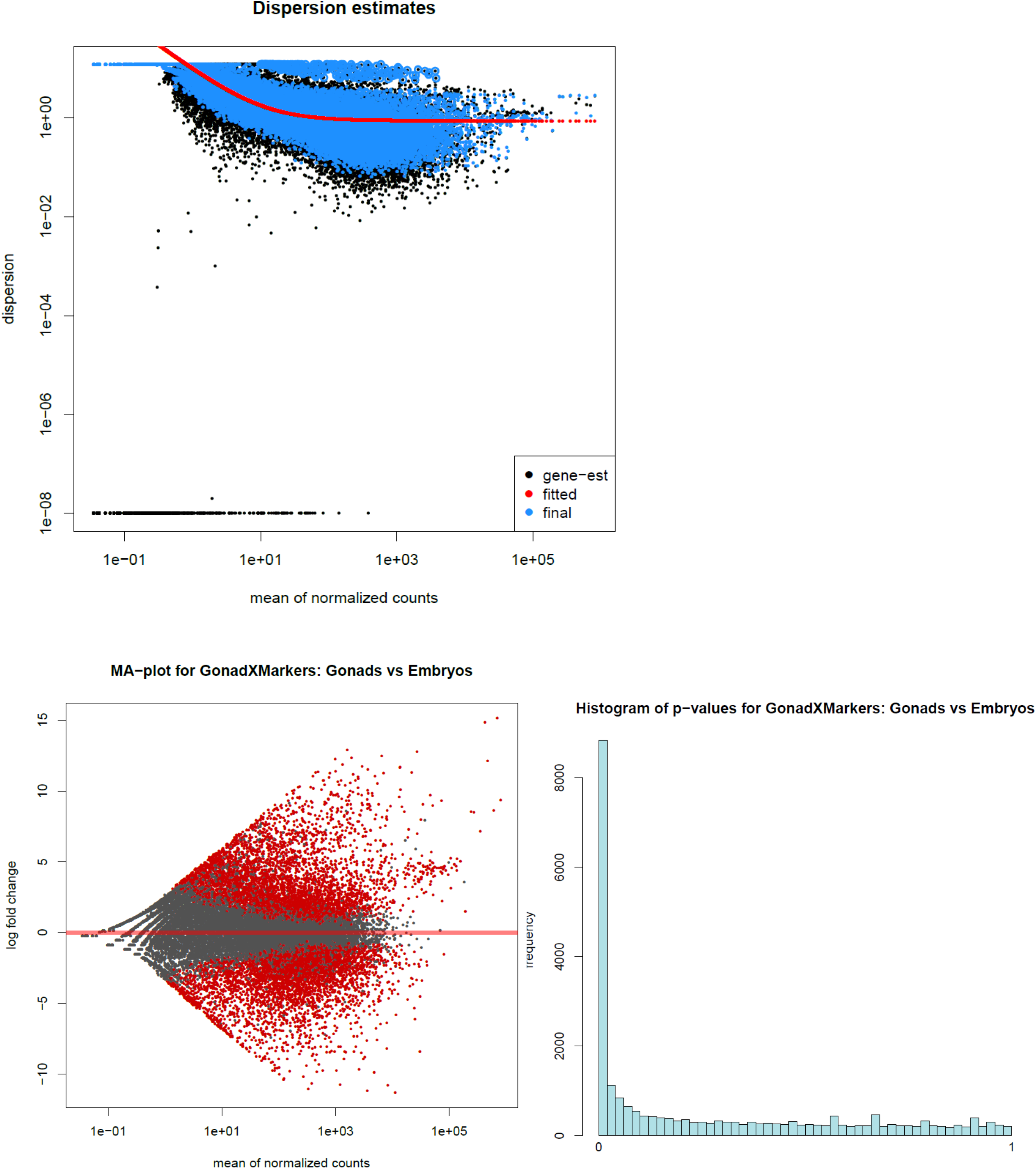
6-way analysis Supplemental Information 2. A. Dispersion estimates for Deseq2 data B. MA-plot of mean relative to log-fold change values C. Histogram of p-value distributions for all data

**Supplementary Figure 14.**
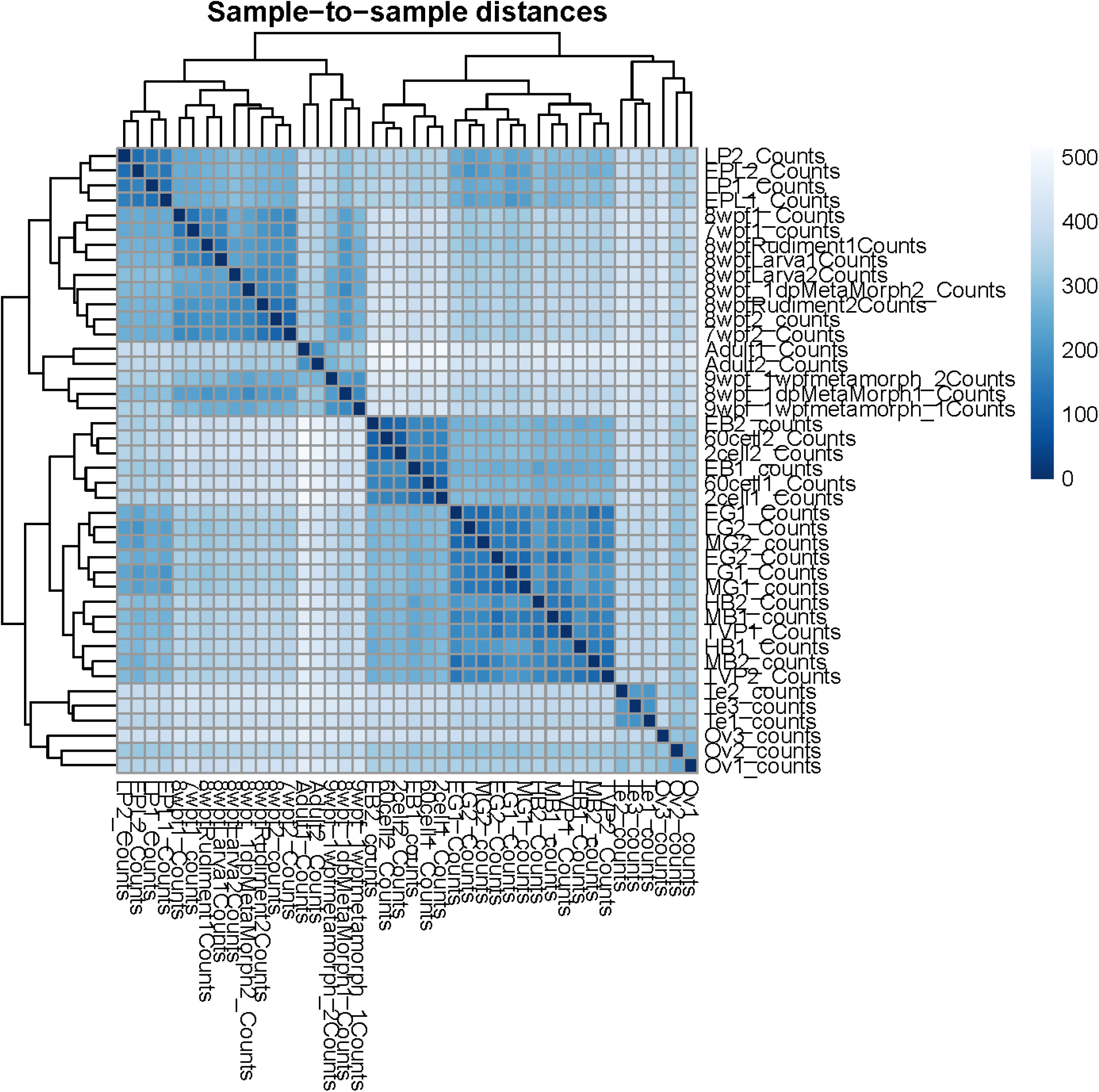
Timecourse Supplemental Information 1. A. Matrix of sample-sample mean distances

**Supplementary Figure 15.**
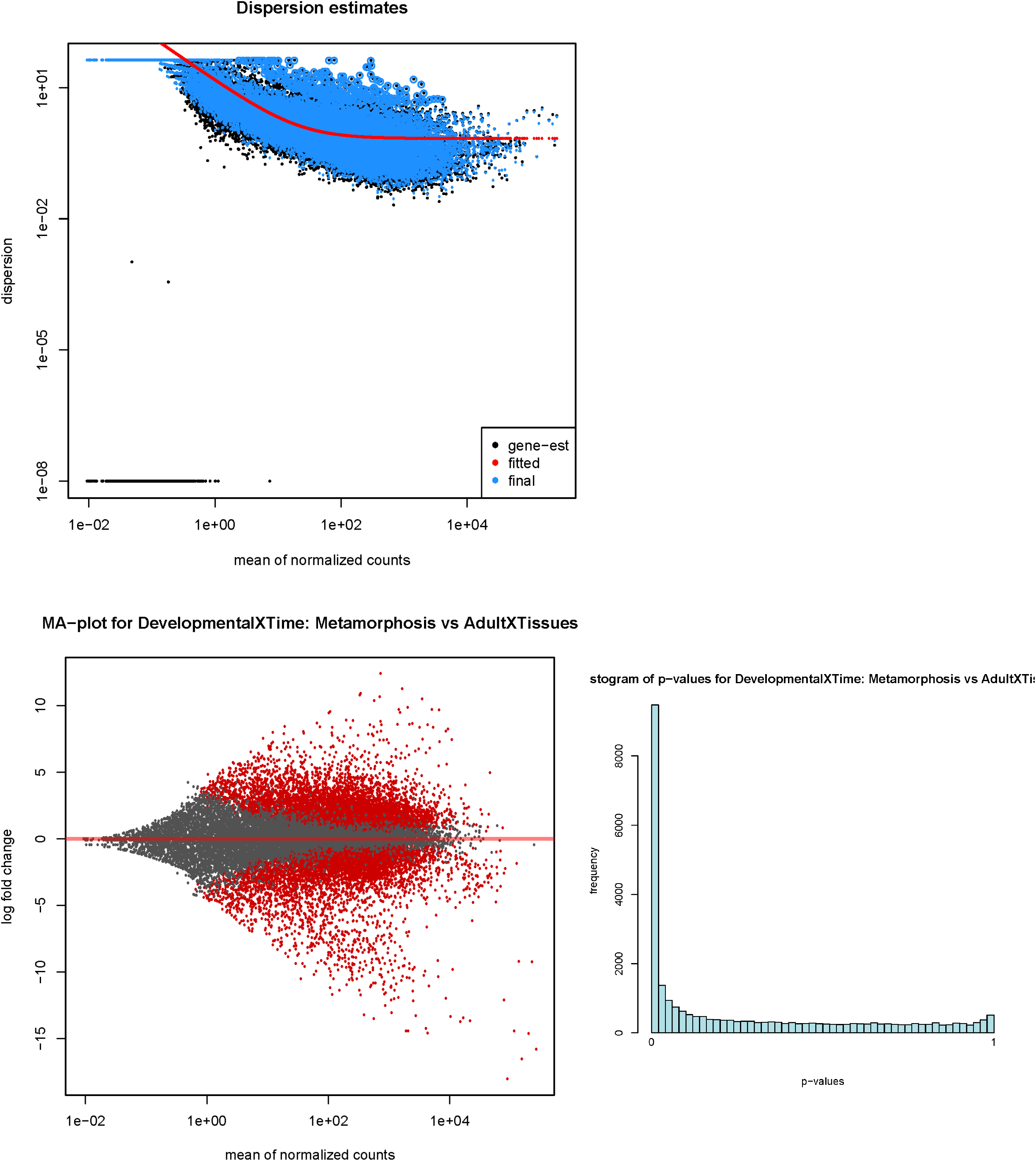
Timecourse Supplemental Information 2. A. Dispersion estimates for Deseq2 data B. MA-plot of mean relative to log-fold change values C. Histogram of p-value distributions for all data

