## Supplementary material for "Sex-Specific Genes Identified in Sea Urchin Gonads are Expressed Prior to Metamorphosis": Pieplow Supplemental data

### Supplemental Data Part 1: novel genes identified in Gonad RNA-seq

#### Novel Genes- Testis

#### L_var_05065_testis- extracellular transmembrane protein

```
##interproscan-version 5.45-80.0
##sequence-region VIRT-161451:53 1 172
VIRT-161451:53 .	polypeptide	1	172	.	+	.	ID=VIRT-
161451:53;md5=475c0002b80bf4dc140be200c1ea00e5
VIRT-161451:53 Phobius	protein_match	153	172	.	+	.	date=24-06-2020;Target=VIRT-161451:53
153 172;ID=match$1_153_172;signature_desc=Region of a membrane-bound protein predicted to be outside the
membrane, in the cytoplasm.;Name=CYTOPLASMIC_DOMAIN;status=T
VIRT-161451:53 Phobius	protein_match	133	152	.	+	.	date=24-06-2020;Target=VIRT-161451:53
133 152;ID=match$2_133_152;signature_desc=Region of a membrane-bound protein predicted to be embedded in the
membrane.;Name=TRANSMEMBRANE;status=T
VIRT-161451:53 MobiDBLite	protein_match	1	30	.	+	.	date=24-06-2020;Target=VIRT-
161451:53 1 30;ID=match$3_1_30;signature_desc=consensus disorder prediction;Name=mobidb-lite;status=T
VIRT-161451:53 Phobius	protein_match	1	132	.	+	.	date=24-06-2020;Target=VIRT-161451:53
1 132;ID=match$4_1_132;signature_desc=Region of a membrane-bound protein predicted to be outside the membrane,
in the extracellular region.;Name=NON_CYTOPLASMIC_DOMAIN;status=T
##FASTA
>VIRT-161451:53
MKGDESDETGNSTGVEKDLSMDVEKEVENETLRRPMAMGWVRRASKQVESAGEAFRKIGL
KVKLANKLGSKKKKKTETDEAPRLFTFDPNAIERGADICDLFETDEVEEDFNDVVDLDDE
RVEEVVSQLVDSMWFRGFILGVIVTNAILIGAQTNQELVRNLHRLFYMLMLF
>match$1_153_172
QTNQELVRNLHRLFYMLMLF
>match$2_133_152
MWFRGFILGVIVTNAILIGA
>match$3_1_30
MKGDESDETGNSTGVEKDLSMDVEKEVENE
>match$4_1_132
MKGDESDETGNSTGVEKDLSMDVEKEVENETLRRPMAMGWVRRASKQVESAGEAFRKIGL
KVKLANKLGSKKKKKTETDEAPRLFTFDPNAIERGADICDLFETDEVEEDFNDVVDLDDE
RVEEVVSQLVDS
```

#### L_var_05606_testis- 7-pass transmembrane protein (GPCR)

```
##interproscan-version 5.45-80.0
##sequence-region VIRT-186776:53 1 419
VIRT-186776:53 .	polypeptide	1	419	.	+	.	ID=VIRT-
186776:53;md5=63f0d8cc70b48d3e31c87c1ee0a2d3c8
VIRT-186776:53 Phobius	protein_match	28	50	.	+	.	date=23-06-2020;Target=VIRT-186776:53
28 50;ID=match$1_28_50;signature_desc=Region of a membrane-bound protein predicted to be embedded in the
membrane.;Name=TRANSMEMBRANE;status=T
VIRT-186776:53 Phobius	protein_match	235	258	.	+	.	date=23-06-2020;Target=VIRT-186776:53
235 258;ID=match$2_235_258;signature_desc=Region of a membrane-bound protein predicted to be embedded in the
membrane.;Name=TRANSMEMBRANE;status=T
VIRT-186776:53 Phobius	protein_match	123	193	.	+	.	date=23-06-2020;Target=VIRT-186776:53
123 193;ID=match$3_123_193;signature_desc=Region of a membrane-bound protein predicted to be outside the
membrane, in the cytoplasm.;Name=CYTOPLASMIC_DOMAIN;status=T
VIRT-186776:53 Phobius	protein_match	329	346	.	+	.	date=23-06-2020;Target=VIRT-186776:53
329 346;ID=match$4_329_346;signature_desc=Region of a membrane-bound protein predicted to be embedded in the
membrane.;Name=TRANSMEMBRANE;status=T
VIRT-186776:53 TMHMM	protein_match	21	43	.	+	.	date=23-06-2020;Target=VIRT-186776:53
21 43;ID=match$5_21_43;signature_desc=Region of a membrane-bound protein predicted to be embedded in the
membrane.;Name=TMhelix;status=T
VIRT-186776:53 TMHMM	protein_match	100	122	.	+	.	date=23-06-2020;Target=VIRT-186776:53
100 122;ID=match$6_100_122;signature_desc=Region of a membrane-bound protein predicted to be embedded in the
membrane.;Name=TMhelix;status=T
VIRT-186776:53 Phobius	protein_match	347	419	.	+	.	date=23-06-2020;Target=VIRT-186776:53
347 419;ID=match$7_347_419;signature_desc=Region of a membrane-bound protein predicted to be outside the
membrane, in the extracellular region.;Name=NON_CYTOPLASMIC_DOMAIN;status=T
VIRT-186776:53 Phobius	protein_match	194	215	.	+	.	date=23-06-2020;Target=VIRT-186776:53
194 215;ID=match$8_194_215;signature_desc=Region of a membrane-bound protein predicted to be embedded in the
membrane.;Name=TRANSMEMBRANE;status=T
VIRT-186776:53 MobiDBLite	protein_match	54	95	.	+	.	date=23-06-2020;Target=VIRT-
186776:53 54 95;ID=match$9_54_95;signature_desc=consensus disorder prediction;Name=mobidb-lite;status=T
VIRT-186776:53 Phobius	protein_match	216	234	.	+	.	date=23-06-2020;Target=VIRT-186776:53
216 234;ID=match$10_216_234;signature_desc=Region of a membrane-bound protein predicted to be outside the
membrane, in the extracellular region.;Name=NON_CYTOPLASMIC_DOMAIN;status=T
```

```
VIRT-186776:53 Phobius protein_match  1       27      .       +       .       date=23-06-2020;Target=VIRT-186776:53
1 27;ID=match$11_1_27;signature_desc=Region of a membrane-bound protein predicted to be outside the membrane, in
the cytoplasm.;Name=CYTOPLASMIC_DOMAIN;status=T
VIRT-186776:53 TMHMM   protein_match  239     258     .       +       .       date=23-06-2020;Target=VIRT-186776:53
239 258;ID=match$12_239_258;signature_desc=Region of a membrane-bound protein predicted to be embedded in the
membrane.;Name=TMhelix;status=T
VIRT-186776:53 Phobius protein_match  259     328     .       +       .       date=23-06-2020;Target=VIRT-186776:53
259 328;ID=match$13_259_328;signature_desc=Region of a membrane-bound protein predicted to be outside the
membrane, in the cytoplasm.;Name=CYTOPLASMIC_DOMAIN;status=T
VIRT-186776:53 TMHMM   protein_match  329     351     .       +       .       date=23-06-2020;Target=VIRT-186776:53
329 351;ID=match$14_329_351;signature_desc=Region of a membrane-bound protein predicted to be embedded in the
membrane.;Name=TMhelix;status=T
VIRT-186776:53 Phobius protein_match  102     122     .       +       .       date=23-06-2020;Target=VIRT-186776:53
102 122;ID=match$15_102_122;signature_desc=Region of a membrane-bound protein predicted to be embedded in the
membrane.;Name=TRANSMEMBRANE;status=T
VIRT-186776:53 Phobius protein_match  51      101     .       +       .       date=23-06-2020;Target=VIRT-186776:53
51 101;ID=match$16_51_101;signature_desc=Region of a membrane-bound protein predicted to be outside the
membrane, in the extracellular region.;Name=NON_CYTOPLASMIC_DOMAIN;status=T
VIRT-186776:53 TMHMM   protein_match  197     219     .       +       .       date=23-06-2020;Target=VIRT-186776:53
197 219;ID=match$17_197_219;signature_desc=Region of a membrane-bound protein predicted to be embedded in the
membrane.;Name=TMhelix;status=T
##FASTA
>VIRT-186776:53
MFLNKSLNRCCNHGIRWRLRHLLIRLDAWKLLIATPLIVFLILDTLPVAIASTPEPTSEP
VAEATSEPENEPTAEAEGPTAEPEDEPQSYSEAGGTPSGAEIVIPPFILMLVIMTGLGVL
MIKAQLRRDRYAERKLTEDYFTQPSLRDLSARFGGAEMAGVQEMREAAKYAFLSMSDIDT
IHSMELFFNRPLPLCRIALPALLSLAFCPLVLYIYSPVARFFWPPGDFPEGVPDINDAIA
CFLMPAGLVYATSFGFSYQSAMEKQRSLVSIVSGEVSKLDQILVMTSHLTYITLKEKALD
SLMDHVVGLNSDSSERYEVMMTRIHPLKWLFLEALGFFAFFGVLLLDVNLSALPDIVDKA
SFFYERAIEGKHVDGPADPGPLTTENSIESAGDLGMVKKTSSHSINEVVLESYDHDHAV
>match$1_28_50
AWKLLIATPLIVFLILDTLPVAI
>match$2_235_258
INDAIACFLMPAGLVYATSFGFSY
>match$3_123_193
KAQLRRDRYAERKLTEDYFTQPSLRDLSARFGGAEMAGVQEMREAAKYAFLSMSDIDTIH
SMELFFNRPLP
>match$4_329_346
WLFLEALGFFAFFGVLLL
>match$5_21_43
HLLIRLDAWKLLIATPLIVFLIL
>match$6_100_122
AEIVIPPFILMLVIMTGLGVLMI
>match$7_347_419
DVNLSALPDIVDKASFFYERAIEGKHVDGPADPGPLTTENSIESAGDLGMVKKTSSHSIN
EVVLESYDHDHAV
>match$8_194_215
LCRIALPALLSLAFCPLVLYIY
>match$9_54_95
PEPTSEPVAEATSEPENEPTAEAEGPTAEPEDEPQSYSEAGG
>match$10_216_234
SPVARFFWPPGDFPEGVPD
>match$11_1_27
MFLNKSLNRCCNHGIRWRLRHLLIRLD
>match$12_239_258
IACFLMPAGLVYATSFGFSY
>match$13_259_328
QSAMEKQRSLVSIVSGEVSKLDQILVMTSHLTYITLKEKALDSLMDHVVGLNSDSSERYE
VMMTRIHPLK
>match$14_329_351
WLFLEALGFFAFFGVLLLDVNLS
>match$15_102_122
IVIPPFILMLVIMTGLGVLMI
>match$16_51_101
ASTPEPTSEPVAEATSEPENEPTAEAEGPTAEPEDEPQSYSEAGGTPSGAE
>match$17_197_219
IALPALLSLAFCPLVLYIYSPVA
```

### L_var_05720_testis- implicated in ALS-2-like ortholog

```
##interproscan-version 5.45-80.0
##sequence-region VIRT-164151:53 1 236
VIRT-164151:53 .       polypeptide    1       236     .       +       .       ID=VIRT-
164151:53;md5=bc7721c26657395fe624f9428319c7c8
```

```
VIRT-164151:53 Pfam    protein_match  75      161     9.0E-12 +       .       date=24-06-2020;Target=VIRT-164151:53
75 161;ID=match$1_75_161;signature_desc=Amyotrophic lateral sclerosis 2 candidate
11;Name=PF15729;status=T;Dbxref="InterPro:IPR031462"
##FASTA
>VIRT-164151:53
MSEDVTLLEEEDAEEDGYNANSAQYVTAPPMVVAEPELYLRIRVGANSKCTKTFNWTAGS
KYVVVHEIKHFPLMTIAALDFEICFAYGIFGYGYSHQLQNRQKKLEDIVSHSYLYRSEPP
MNRKQSDNTTMAPVPIEHPDLISFAQKVQIGSPAVREMLAPPGTTVLPDIDQVAAPEELV
YGEDDDHTCVTDVPELSSFLANAPGNFMVSFANKLRLWKSKGAPAKLPNTGPVTSE
>match$1_75_161
TIAALDFEICFAYGIFGYGYSHQLQNRQKKLEDIVSHSYLYRSEPPMNRKQSDNTTMAPV
PIEHPDLISFAQKVQIGSPAVREMLAP
```

### L_var_12731_testis- disordered protein

```
##interproscan-version 5.45-80.0
##sequence-region SPU_023211.3a 1 418
SPU_023211.3a  .       polypeptide    1       418     .       +       .
       ID=SPU_023211.3a;md5=f151280ddaabdabef7a1469ab6b282f9
SPU_023211.3a  PANTHER protein_match  32      232     1.2E-15 +       .       date=24-06-2020;Target=SPU_023211.3a
32
232;Ontology_term="GO:0005524","GO:0006139","GO:0019205";ID=match$1_32_232;Name=PTHR23359;status=T;Dbxref="Inter
Pro:IPR000850","KEGG:00230+2.7.4.3","KEGG:00730+2.7.4.3","MetaCyc:PWY-7219","Reactome:R-HSA-499943"
SPU_023211.3a  MobiDBLite     protein_match  179     199     .       +       .       date=24-06-
2020;Target=SPU_023211.3a 179 199;ID=match$2_179_199;signature_desc=consensus disorder prediction;Name=mobidb-
lite;status=T
SPU_023211.3a  Gene3D  protein_match  37      258     4.8E-20 +       .       date=24-06-2020;Target=SPU_023211.3a
37 258;ID=match$3_37_258;Name=G3DSA:3.40.50.300;status=T
SPU_023211.3a  PANTHER protein_match  32      232     1.2E-15 +       .       date=24-06-2020;Target=SPU_023211.3a
32 232;ID=match$4_32_232;Name=PTHR23359:SF88;status=T
##FASTA
>SPU_023211.3a
MGCGNSTDGKGEDGEKGFFHRRPKVSIRVGNSVKLSTDETTIVFIFGGPGSRKGRIVDDM
LHMYGFKVINVEDILLRELPNKANISAAGSIRETMSIVSENAGLINLEWIFEMISDEIER
APKYIYLVDIVPNLRFLLRCGALTADPEYELKKFTQKYPSYFALNLAIHENKILEDAKAH
GAKHPDKMKEGGQSDEADSSRLARRNVLYQNSVKAWLGYFRNDHRLVTVDVSCGVSDLIW
HRVCDFFGHDLECLPRKTVNTVVLFAFNGYDYSAIDMQRYHMELIHLKDILPKPDMSADA
ALHTLSKHIDKTAPTAESFCVDLSSTTITKDMERLDHSKREFRREMRFIEWHDTFLDKYI
FTSSEKKERGRKSSLFIQTFKVIATTENEVCLFPPETDVGLCQDLAFSFGHHRHPSAR
>match$1_32_232
SVKLSTDETTIVFIFGGPGSRKGRIVDDMLHMYGFKVINVEDILLRELPNKANISAAGSI
RETMSIVSENAGLINLEWIFEMISDEIERAPKYIYLVDIVPNLRFLLRCGALTADPEYEL
KKFTQKYPSYFALNLAIHENKILEDAKAHGAKHPDKMKEGGQSDEADSSRLARRNVLYQN
SVKAWLGYFRNDHRLVTVDVS
>match$2_179_199
AHGAKHPDKMKEGGQSDEADS
>match$3_37_258
TDETTIVFIFGGPGSRKGRIVDDMLHMYGFKVINVEDILLRELPNKANISAAGSIRETMS
IVSENAGLINLEWIFEMISDEIERAPKYIYLVDIVPNLRFLLRCGALTADPEYELKKFTQ
KYPSYFALNLAIHENKILEDAKAHGAKHPDKMKEGGQSDEADSSRLARRNVLYQNSVKAW
LGYFRNDHRLVTVDVSCGVSDLIWHRVCDFFGHDLECLPRKT
>match$4_32_232
SVKLSTDETTIVFIFGGPGSRKGRIVDDMLHMYGFKVINVEDILLRELPNKANISAAGSI
RETMSIVSENAGLINLEWIFEMISDEIERAPKYIYLVDIVPNLRFLLRCGALTADPEYEL
KKFTQKYPSYFALNLAIHENKILEDAKAHGAKHPDKMKEGGQSDEADSSRLARRNVLYQN
SVKAWLGYFRNDHRLVTVDVS
```

### L_var_13026_testis- 7-pass transmembrane protein (GPCR)

```
##interproscan-version 5.45-80.0
##sequence-region VIRT-162979:53 1 119
VIRT-162979:53 .       polypeptide    1       119     .       +       .       ID=VIRT-
162979:53;md5=a46af7c1151d7b38f37aec90e74ee7a1
VIRT-162979:53 Phobius protein_match  73      83      .       +       .       date=24-06-2020;Target=VIRT-162979:53
73 83;ID=match$1_73_83;signature_desc=Region of a membrane-bound protein predicted to be outside the membrane,
in the cytoplasm.;Name=CYTOPLASMIC_DOMAIN;status=T
VIRT-162979:53 TMHMM   protein_match  85      107     .       +       .       date=24-06-2020;Target=VIRT-162979:53
85 107;ID=match$2_85_107;signature_desc=Region of a membrane-bound protein predicted to be embedded in the
membrane.;Name=TMhelix;status=T
VIRT-162979:53 Phobius protein_match  110     119     .       +       .       date=24-06-2020;Target=VIRT-162979:53
110 119;ID=match$3_110_119;signature_desc=Region of a membrane-bound protein predicted to be outside the
membrane, in the extracellular region.;Name=NON_CYTOPLASMIC_DOMAIN;status=T
VIRT-162979:53 Phobius protein_match  14      35      .       +       .       date=24-06-2020;Target=VIRT-162979:53
14 35;ID=match$4_14_35;signature_desc=Region of a membrane-bound protein predicted to be embedded in the
membrane.;Name=TRANSMEMBRANE;status=T
```

```
VIRT-162979:53 Phobius protein_match  36      54      .       +       .       date=24-06-2020;Target=VIRT-162979:53 36 54;ID=match$5_36_54;signature_desc=Region of a membrane-bound protein predicted to be outside the membrane, in the extracellular region.;Name=NON_CYTOPLASMIC_DOMAIN;status=T
VIRT-162979:53 TMHMM   protein_match  13      35      .       +       .       date=24-06-2020;Target=VIRT-162979:53 13 35;ID=match$6_13_35;signature_desc=Region of a membrane-bound protein predicted to be embedded in the membrane.;Name=TMhelix;status=T
VIRT-162979:53 Phobius protein_match  55      72      .       +       .       date=24-06-2020;Target=VIRT-162979:53 55 72;ID=match$7_55_72;signature_desc=Region of a membrane-bound protein predicted to be embedded in the membrane.;Name=TRANSMEMBRANE;status=T
VIRT-162979:53 Phobius protein_match  84      109     .       +       .       date=24-06-2020;Target=VIRT-162979:53 84 109;ID=match$8_84_109;signature_desc=Region of a membrane-bound protein predicted to be embedded in the membrane.;Name=TRANSMEMBRANE;status=T
VIRT-162979:53 Phobius protein_match  1       13      .       +       .       date=24-06-2020;Target=VIRT-162979:53 1 13;ID=match$9_1_13;signature_desc=Region of a membrane-bound protein predicted to be outside the membrane, in the cytoplasm.;Name=CYTOPLASMIC_DOMAIN;status=T
VIRT-162979:53 TMHMM   protein_match  50      72      .       +       .       date=24-06-2020;Target=VIRT-162979:53 50 72;ID=match$10_50_72;signature_desc=Region of a membrane-bound protein predicted to be embedded in the membrane.;Name=TMhelix;status=T
##FASTA
>VIRT-162979:53
MAFNIWALTQKKTLYVYTSSFSVLVLAFIIVFDLLQYTVFERADMTSQGFTESLLTIHVL
SYYTALLGFVYNTKAGKRLGAAKLEWISVVATFLLSIMRVGIEISFLSFRKQAFKNFNE
>match$1_73_83
TKAGKRLGAAK
>match$2_85_107
EWISVVATFLLSIMRVGIEISFL
>match$3_110_119
RKQAFKNFNE
>match$4_14_35
LYVYTSSFSVLVLAFIIVFDLL
>match$5_36_54
QYTVFERADMTSQGFTESL
>match$6_13_35
TLYVYTSSFSVLVLAFIIVFDLL
>match$7_55_72
LTIHVLSYYTALLGFVYN
>match$8_84_109
LEWISVVATFLLSIMRVGIEISFLSF
>match$9_1_13
MAFNIWALTQKKT
>match$10_50_72
FTESLLTIHVLSYYTALLGFVYN
```

### L_var_18063_testis- 7-pass transmembrane protein (GPCR)

```
##interproscan-version 5.45-80.0
##sequence-region VIRT-75805:53 1 282
VIRT-75805:53  .       polypeptide    1       282     .       +       .       ID=VIRT-75805:53;md5=282afbe4486d0ad63faff2a1c28590ea
VIRT-75805:53  MobiDBLite     protein_match  22      55      .       +       .       date=23-06-2020;Target=VIRT-75805:53 22 55;ID=match$1_22_55;signature_desc=consensus disorder prediction;Name=mobidb-lite;status=T
VIRT-75805:53  MobiDBLite     protein_match  236     282     .       +       .       date=23-06-2020;Target=VIRT-75805:53 236 282;ID=match$2_236_282;signature_desc=consensus disorder prediction;Name=mobidb-lite;status=T
VIRT-75805:53  MobiDBLite     protein_match  200     221     .       +       .       date=23-06-2020;Target=VIRT-75805:53 200 221;ID=match$3_200_221;signature_desc=consensus disorder prediction;Name=mobidb-lite;status=T
VIRT-75805:53  Phobius protein_match  100     121     .       +       .       date=23-06-2020;Target=VIRT-75805:53 100 121;ID=match$4_100_121;signature_desc=Region of a membrane-bound protein predicted to be embedded in the membrane.;Name=TRANSMEMBRANE;status=T
VIRT-75805:53  TMHMM   protein_match  99      121     .       +       .       date=23-06-2020;Target=VIRT-75805:53 99 121;ID=match$5_99_121;signature_desc=Region of a membrane-bound protein predicted to be embedded in the membrane.;Name=TMhelix;status=T
VIRT-75805:53  Phobius protein_match  71      94      .       +       .       date=23-06-2020;Target=VIRT-75805:53 71 94;ID=match$6_71_94;signature_desc=Region of a membrane-bound protein predicted to be embedded in the membrane.;Name=TRANSMEMBRANE;status=T
VIRT-75805:53  Phobius protein_match  122     282     .       +       .       date=23-06-2020;Target=VIRT-75805:53 122 282;ID=match$7_122_282;signature_desc=Region of a membrane-bound protein predicted to be outside the membrane, in the cytoplasm.;Name=CYTOPLASMIC_DOMAIN;status=T
VIRT-75805:53  TMHMM   protein_match  73      95      .       +       .       date=23-06-2020;Target=VIRT-75805:53 73 95;ID=match$8_73_95;signature_desc=Region of a membrane-bound protein predicted to be embedded in the membrane.;Name=TMhelix;status=T
VIRT-75805:53  MobiDBLite     protein_match  204     221     .       +       .       date=23-06-2020;Target=VIRT-75805:53 204 221;ID=match$9_204_221;signature_desc=consensus disorder prediction;Name=mobidb-lite;status=T
VIRT-75805:53  Phobius protein_match  1       70      .       +       .       date=23-06-2020;Target=VIRT-75805:53 1 70;ID=match$10_1_70;signature_desc=Region of a membrane-bound protein predicted to be outside the membrane, in the cytoplasm.;Name=CYTOPLASMIC_DOMAIN;status=T
```

VIRT-75805:53 Phobius protein_match 95 99 . + . date=23-06-2020;Target=VIRT-75805:53 95 99;ID=match$11_95_99;signature_desc=Region of a membrane-bound protein predicted to be outside the membrane, in the extracellular region.;Name=NON_CYTOPLASMIC_DOMAIN;status=T
##FASTA
>VIRT-75805:53
MSVKGHDESSLRGNHQPHFLNPTIQSGSHQGGASGSHSPALTATYTGSSSNRTSGSYGGL
VLRSGRTISPAVLRIIYFGSIFCFISGVLCGNLGGWFDSVVGFLGMFLITVGVITTLFVA
YHRYHQRHSPGGAGTNMTIRPTVTLQQEVTNPSGDGGTTSYVIETRRVQTAAGMPIGPQG
FNTQYGFPQGTATQTFQDHMQNVPGPTQPPPANTPSGGYPQPIPTDPTGTFMTSQGATPQ
FNYPVMHPPQIRGTHLGVTQPESVENDLPPPPSYQDVFSKSY
>match$1_22_55
PTIQSGSHQGGASGSHSPALTATYTGSSSNRTSG
>match$2_236_282
GATPQFNYPVMHPPQIRGTHLGVTQPESVENDLPPPPSYQDVFSKSY
>match$3_200_221
MQNVPGPTQPPPANTPSGGYPQ
>match$4_100_121
VVGFLGMFLITVGVITTLFVAY
>match$5_99_121
SVVGFLGMFLITVGVITTLFVAY
>match$6_71_94
AVLRIIYFGSIFCFISGVLCGNLG
>match$7_122_282
HRYHQRHSPGGAGTNMTIRPTVTLQQEVTNPSGDGGTTSYVIETRRVQTAAGMPIGPQGF
NTQYGFPQGTATQTFQDHMQNVPGPTQPPPANTPSGGYPQPIPTDPTGTFMTSQGATPQF
NYPVMHPPQIRGTHLGVTQPESVENDLPPPPSYQDVFSKSY
>match$8_73_95
LRIIYFGSIFCFISGVLCGNLGG
>match$9_204_221
PGPTQPPPANTPSGGYPQ
>match$10_1_70
MSVKGHDESSLRGNHQPHFLNPTIQSGSHQGGASGSHSPALTATYTGSSSNRTSGSYGGL
VLRSGRTISP
>match$11_95_99
GWFDS

### L_var_22946_testis- very large highly disordered protein

##interproscan-version 5.45-80.0
##sequence-region EEA38816.1 1 1104
EEA38816.1 . polypeptide 1 1104 . + . ID=EEA38816.1;md5=c02134d4fc71a6a7bb741e0989079391
EEA38816.1 MobiDBLite protein_match 330 390 . + . date=23-06-2020;Target=EEA38816.1 330 390;ID=match$1_330_390;signature_desc=consensus disorder prediction;Name=mobidb-lite;status=T
EEA38816.1 MobiDBLite protein_match 409 599 . + . date=23-06-2020;Target=EEA38816.1 409 599;ID=match$2_409_599;signature_desc=consensus disorder prediction;Name=mobidb-lite;status=T
EEA38816.1 PANTHER protein_match 155 296 1.2E-242 + . date=23-06-2020;Target=EEA38816.1 155 296;ID=match$3_155_296;Name=PTHR22192:SF17;status=T
EEA38816.1 PANTHER protein_match 793 1104 1.2E-242 + . date=23-06-2020;Target=EEA38816.1 793 1104;ID=match$4_793_1104;Name=PTHR22192;status=T;Dbxref="InterPro:IPR026715"
EEA38816.1 PANTHER protein_match 295 689 1.2E-242 + . date=23-06-2020;Target=EEA38816.1 295 689;ID=match$4_295_689;Name=PTHR22192;status=T;Dbxref="InterPro:IPR026715"
EEA38816.1 MobiDBLite protein_match 456 568 . + . date=23-06-2020;Target=EEA38816.1 456 568;ID=match$5_456_568;signature_desc=consensus disorder prediction;Name=mobidb-lite;status=T
EEA38816.1 MobiDBLite protein_match 133 159 . + . date=23-06-2020;Target=EEA38816.1 133 159;ID=match$6_133_159;signature_desc=consensus disorder prediction;Name=mobidb-lite;status=T
EEA38816.1 MobiDBLite protein_match 583 599 . + . date=23-06-2020;Target=EEA38816.1 583 599;ID=match$7_583_599;signature_desc=consensus disorder prediction;Name=mobidb-lite;status=T
EEA38816.1 MobiDBLite protein_match 220 248 . + . date=23-06-2020;Target=EEA38816.1 220 248;ID=match$8_220_248;signature_desc=consensus disorder prediction;Name=mobidb-lite;status=T
EEA38816.1 PANTHER protein_match 793 1104 1.2E-242 + . date=23-06-2020;Target=EEA38816.1 793 1104;ID=match$9_793_1104;Name=PTHR22192:SF17;status=T
EEA38816.1 PANTHER protein_match 295 689 1.2E-242 + . date=23-06-2020;Target=EEA38816.1 295 689;ID=match$9_295_689;Name=PTHR22192:SF17;status=T
EEA38816.1 MobiDBLite protein_match 105 130 . + . date=23-06-2020;Target=EEA38816.1 105 130;ID=match$10_105_130;signature_desc=consensus disorder prediction;Name=mobidb-lite;status=T
EEA38816.1 MobiDBLite protein_match 719 735 . + . date=23-06-2020;Target=EEA38816.1 719 735;ID=match$11_719_735;signature_desc=consensus disorder prediction;Name=mobidb-lite;status=T

EEA38816.1 MobiDBLite protein_match 1 390 . + . date=23-06-2020;Target=EEA38816.1 1 390;ID=match$12_1_390;signature_desc=consensus disorder prediction;Name=mobidb-lite;status=T
EEA38816.1 MobiDBLite protein_match 746 761 . + . date=23-06-2020;Target=EEA38816.1 746 761;ID=match$13_746_761;signature_desc=consensus disorder prediction;Name=mobidb-lite;status=T
EEA38816.1 MobiDBLite protein_match 409 444 . + . date=23-06-2020;Target=EEA38816.1 409 444;ID=match$14_409_444;signature_desc=consensus disorder prediction;Name=mobidb-lite;status=T
EEA38816.1 PANTHER protein_match 155 296 1.2E-242 + . date=23-06-2020;Target=EEA38816.1 155 296;ID=match$15_155_296;Name=PTHR22192;status=T;Dbxref="InterPro:IPR026715"
EEA38816.1 MobiDBLite protein_match 249 279 . + . date=23-06-2020;Target=EEA38816.1 249 279;ID=match$16_249_279;signature_desc=consensus disorder prediction;Name=mobidb-lite;status=T
EEA38816.1 Pfam protein_match 938 1051 8.2E-12 + . date=23-06-2020;Target=EEA38816.1 938 1051;ID=match$17_938_1051;signature_desc=Speriolin C-terminus;Name=PF15059;status=T;Dbxref="InterPro:IPR029384"
EEA38816.1 Pfam protein_match 1060 1104 1.3E-7 + . date=23-06-2020;Target=EEA38816.1 1060 1104;ID=match$17_1060_1104;signature_desc=Speriolin C-terminus;Name=PF15059;status=T;Dbxref="InterPro:IPR029384"
EEA38816.1 MobiDBLite protein_match 160 212 . + . date=23-06-2020;Target=EEA38816.1 160 212;ID=match$18_160_212;signature_desc=consensus disorder prediction;Name=mobidb-lite;status=T
EEA38816.1 MobiDBLite protein_match 857 929 . + . date=23-06-2020;Target=EEA38816.1 857 929;ID=match$19_857_929;signature_desc=consensus disorder prediction;Name=mobidb-lite;status=T
EEA38816.1 MobiDBLite protein_match 688 762 . + . date=23-06-2020;Target=EEA38816.1 688 762;ID=match$20_688_762;signature_desc=consensus disorder prediction;Name=mobidb-lite;status=T
EEA38816.1 MobiDBLite protein_match 1 36 . + . date=23-06-2020;Target=EEA38816.1 1 36;ID=match$21_1_36;signature_desc=consensus disorder prediction;Name=mobidb-lite;status=T
EEA38816.1 Coils protein_match 601 628 . + . date=23-06-2020;Target=EEA38816.1 601 628;ID=match$22_601_628;Name=Coil;status=T
EEA38816.1 Coils protein_match 360 380 . + . date=23-06-2020;Target=EEA38816.1 360 380;ID=match$23_360_380;Name=Coil;status=T
EEA38816.1 MobiDBLite protein_match 310 329 . + . date=23-06-2020;Target=EEA38816.1 310 329;ID=match$24_310_329;signature_desc=consensus disorder prediction;Name=mobidb-lite;status=T
EEA38816.1 Coils protein_match 170 190 . + . date=23-06-2020;Target=EEA38816.1 170 190;ID=match$25_170_190;Name=Coil;status=T
EEA38816.1 MobiDBLite protein_match 280 309 . + . date=23-06-2020;Target=EEA38816.1 280 309;ID=match$26_280_309;signature_desc=consensus disorder prediction;Name=mobidb-lite;status=T
EEA38816.1 MobiDBLite protein_match 45 94 . + . date=23-06-2020;Target=EEA38816.1 45 94;ID=match$27_45_94;signature_desc=consensus disorder prediction;Name=mobidb-lite;status=T
##FASTA
>EEA38816.1
MASPLEPGSSLSTISPSGTPQVTPSVSRQSSRLTEGGEESASRRPSKVMFTSTPKSERSR
GASSVQLCQDSSGSVLSDITQLRRNSSTLSDGEQLHRDSGVLSDRRDSSNLSTQSDQLRR
NSSVLSDLRDSGVPSDQLRRDSTISDREQLYRDSSVLSDRRDSSNISTQCNQLRQDSSNL
STQSDQLRRDSSVLSGQLHRDSSVMSDQLRRDSSVLSNRDQLRRDSSVLSGQLRRDSTTM
STKTDQPRRDSSVLSRQLHRNDNMQSDSEQLRRDSGVLSDQVHGESSVMSGQLRQDNSTM
SNQSDQVRRDSSTLPDRDQLRRDSSVLSDQPRRDSSVLSVQLHRDSSMQSGQLRQDSSML
AQLRRDSRTMSTQSDQLRRDSSTLSDRYPSTLCTNDLEAWDSLPRGYLTSSSSGLLSPQK
SQEKGDVLTRESTPASSRIFAGSTERSRDTFARDFPSMPSLQSQSSHQSRDSTAPLGRVS
SGPSELTSQEKSSDMDTSLSNKTSSGSTTKLFSAPSQESAASSPQRVQEPSVGTTPVLSP
RKSYESSAEALGSSRSTSKTSTRESSSLLGGVRHKATAMTRGSGPSGAESSSGSPDDSEI
LRRLTNMNERMCNENDLLRKQLKLFKRNAELKQISLISLVIAWHQGLISADECESLSELS
QLCPEDGSGGASVVSGRRLSRCLQELASLRPPRGAATRRREREASGPDSVSQESSERSPR
SAVTRRREREASDPDAVSQESSGRSAREASTRRRTRKQSLPDPILEDISVRLSRDSVTRR
RARDASGPDPLWLESSVREIKRVLHHLERTLIDLDSTVRDHQYLWRDVRRRGTLPLPAPG
SEEGVSAPRLYIPRPHAEQQGTADSGEGVSAPRLYTPRPHAEHAGTAVQFEKKEAAQPND
SGLGGSFSERATFPGGEQAMDSRARRTSETHDVREARLLGEVCHQLEQRVLVSVFPRQKI
DHSSFSLRTIGDLVRLERDPDEKEFLTRKLGRVTATLRLYGFDFQRHPFFMTDVILRFGK
FPTDWDKYNFVHQKGYHDPELLREMVGRLVTIVTRVLPPGYHDPELLREAVGRLVPEPFR
YDMLVVLDCLLAMAREDGCSIFLW
>match$1_330_390
QPRRDSSVLSVQLHRDSSMQSGQLRQDSSMLAQLRRDSRTMSTQSDQLRRDSSTLSDRYP
S
>match$2_409_599
TSSSSGLLSPQKSQEKGDVLTRESTPASSRIFAGSTERSRDTFARDFPSMPSLQSQSSHQ
SRDSTAPLGRVSSGPSELTSQEKSSDMDTSLSNKTSSGSTTKLFSAPSQESAASSPQRVQ
EPSVGTTPVLSPRKSYESSAEALGSSRSTSKTSTRESSSLLGGVRHKATAMTRGSGPSGA
ESSSGSPDDSE

>match$3_155_296
SVLSDRRDSSNISTQCNQLRQDSSNLSTQSDQLRRDSSVLSGQLHRDSSVMSDQLRRDSS
VLSNRDQLRRDSSVLSGQLRRDSTTMSTKTDQPRRDSSVLSRQLHRNDNMQSDSEQLRRD
SGVLSDQVHGESSVMSGQLRQD
>match$4_295_689
QDNSTMSNQSDQVRRDSSTLPDRDQLRRDSSVLSDQPRRDSSVLSVQLHRDSSMQSGQLR
QDSSMLAQLRRDSRTMSTQSDQLRRDSSTLSDRYPSTLCTNDLEAWDSLPRGYLTSSSSG
LLSPQKSQEKGDVLTRESTPASSRIFAGSTERSRDTFARDFPSMPSLQSQSSHQSRDSTA
PLGRVSSGPSELTSQEKSSDMDTSLSNKTSSGSTTKLFSAPSQESAASSPQRVQEPSVGT
TPVLSPRKSYESSAEALGSSRSTSKTSTRESSSLLGGVRHKATAMTRGSGPSGAESSSGS
PDDSEILRRLTNMNERMCNENDLLRKQLKLFKRNAELKQISLISLVIAWHQGLISADECE
SLSELSQLCPEDGSGGASVVSGRRLSRCLQELASL
>match$4_793_1104
LESSVREIKRVLHHLERTLIDLDSTVRDHQYLWRDVRRRGTLPLPAPGSEEGVSAPRLYI
PRPHAEQQGTADSGEGVSAPRLYTPRPHAEHAGTAVQFEKKEAAQPNDSGLGGSFSERAT
FPGGEQAMDSRARRTSETHDVREARLLGEVCHQLEQRVLVSVFPRQKIDHSSFSLRTIGD
LVRLERDPDEKEFLTRKLGRVTATLRLYGFDFQRHPFFMTDVILRFGKFPTDWDKYNFVH
QKGYHDPELLREMVGRLVTIVTRVLPPGYHDPELLREAVGRLVPEPFRYDMLVVLDCLLA
MAREDGCSIFLW
>match$5_456_568
PSMPSLQSQSSHQSRDSTAPLGRVSSGPSELTSQEKSSDMDTSLSNKTSSGSTTKLFSAP
SQESAASSPQRVQEPSVGTTPVLSPRKSYESSAEALGSSRSTSKTSTRESSSL
>match$6_133_159
VPSDQLRRDSTISDREQLYRDSSVLSD
>match$7_583_599
SGPSGAESSSGSPDDSE
>match$8_220_248
DQLRRDSSVLSGQLRRDSTTMSTKTDQPR
>match$9_295_689
QDNSTMSNQSDQVRRDSSTLPDRDQLRRDSSVLSDQPRRDSSVLSVQLHRDSSMQSGQLR
QDSSMLAQLRRDSRTMSTQSDQLRRDSSTLSDRYPSTLCTNDLEAWDSLPRGYLTSSSSG
LLSPQKSQEKGDVLTRESTPASSRIFAGSTERSRDTFARDFPSMPSLQSQSSHQSRDSTA
PLGRVSSGPSELTSQEKSSDMDTSLSNKTSSGSTTKLFSAPSQESAASSPQRVQEPSVGT
TPVLSPRKSYESSAEALGSSRSTSKTSTRESSSLLGGVRHKATAMTRGSGPSGAESSSGS
PDDSEILRRLTNMNERMCNENDLLRKQLKLFKRNAELKQISLISLVIAWHQGLISADECE
SLSELSQLCPEDGSGGASVVSGRRLSRCLQELASL
>match$9_793_1104
LESSVREIKRVLHHLERTLIDLDSTVRDHQYLWRDVRRRGTLPLPAPGSEEGVSAPRLYI
PRPHAEQQGTADSGEGVSAPRLYTPRPHAEHAGTAVQFEKKEAAQPNDSGLGGSFSERAT
FPGGEQAMDSRARRTSETHDVREARLLGEVCHQLEQRVLVSVFPRQKIDHSSFSLRTIGD
LVRLERDPDEKEFLTRKLGRVTATLRLYGFDFQRHPFFMTDVILRFGKFPTDWDKYNFVH
QKGYHDPELLREMVGRLVTIVTRVLPPGYHDPELLREAVGRLVPEPFRYDMLVVLDCLLA
MAREDGCSIFLW
>match$10_105_130
RRDSSNLSTQSDQLRRNSSVLSDLRD
>match$11_719_735
PRSAVTRRREREASDPD
>match$12_1_390
MASPLEPGSSLSTISPSGTPQVTPSVSRQSSRLTEGGEESASRRPSKVMFTSTPKSERSR
GASSVQLCQDSSGSVLSDITQLRRNSSTLSDGEQLHRDSGVLSDRRDSSNLSTQSDQLRR
NSSVLSDLRDSGVPSDQLRRDSTISDREQLYRDSSVLSDRRDSSNISTQCNQLRQDSSNL
STQSDQLRRDSSVLSGQLHRDSSVMSDQLRRDSSVLSNRDQLRRDSSVLSGQLRRDSTTM
STKTDQPRRDSSVLSRQLHRNDNMQSDSEQLRRDSGVLSDQVHGESSVMSGQLRQDNSTM
SNQSDQVRRDSSTLPDRDQLRRDSSVLSDQPRRDSSVLSVQLHRDSSMQSGQLRQDSSML
AQLRRDSRTMSTQSDQLRRDSSTLSDRYPS
>match$13_746_761
AREASTRRRTRKQSLP
>match$14_409_444
TSSSSGLLSPQKSQEKGDVLTRESTPASSRIFAGST
>match$15_155_296
SVLSDRRDSSNISTQCNQLRQDSSNLSTQSDQLRRDSSVLSGQLHRDSSVMSDQLRRDSS
VLSNRDQLRRDSSVLSGQLRRDSTTMSTKTDQPRRDSSVLSRQLHRNDNMQSDSEQLRRD
SGVLSDQVHGESSVMSGQLRQD
>match$16_249_279
RDSSVLSRQLHRNDNMQSDSEQLRRDSGVLS
>match$17_938_1051
LLGEVCHQLEQRVLVSVFPRQKIDHSSFSLRTIGDLVRLERDPDEKEFLTRKLGRVTATL
RLYGFDFQRHPFFMTDVILRFGKFPTDWDKYNFVHQKGYHDPELLREMVGRLVT
>match$17_1060_1104
GYHDPELLREAVGRLVPEPFRYDMLVVLDCLLAMAREDGCSIFLW
>match$18_160_212
RRDSSNISTQCNQLRQDSSNLSTQSDQLRRDSSVLSGQLHRDSSVMSDQLRRD
>match$19_857_929
AEQQGTADSGEGVSAPRLYTPRPHAEHAGTAVQFEKKEAAQPNDSGLGGSFSERATFPGG
EQAMDSRARRTSE

```
>match$20_688_762
SLRPPRGAATRRREREASGPDSVSQESSERSPRSAVTRRREREASDPDAVSQESSGRSAR
EASTRRRTRKQSLPD
>match$21_1_36
MASPLEPGSSLSTISPSGTPQVTPSVSRQSSRLTEG
>match$22_601_628
LRRLTNMNERMCNENDLLRKQLKLFKRN
>match$23_360_380
LAQLRRDSRTMSTQSDQLRRD
>match$24_310_329
DSSTLPDRDQLRRDSSVLSD
>match$25_170_190
CNQLRQDSSNLSTQSDQLRRD
>match$26_280_309
DQVHGESSVMSGQLRQDNSTMSNQSDQVRR
>match$27_45_94
PSKVMFTSTPKSERSRGASSVQLCQDSSGSVLSDITQLRRNSSTLSDGEQ
```

#### L_var_24202_testis- very small coiled coil protein

```
##interproscan-version 5.45-80.0
##sequence-region VIRT-173056:53 1 80
VIRT-173056:53 .	polypeptide	1	80	.	+	.	ID=VIRT-
173056:53;md5=1063a2f9588863c67190775061236b39
VIRT-173056:53 Coils	protein_match	1	28	.	+	.	date=23-06-2020;Target=VIRT-173056:53
1 28;ID=match$1_1_28;Name=Coil;status=T
##FASTA
>VIRT-173056:53
MRKYESQLEVLRAKNESLQRSLQSALDSQGSLGVQINQWQACIDSAGQQLCRVTQEVHES
KSREKHLRHLLGEERRIKVN
>match$1_1_28
MRKYESQLEVLRAKNESLQRSLQSALDS
```

#### L_var_27142_testis- another highly disordered protein

```
##interproscan-version 5.45-80.0
##sequence-region VIRT-153210:53 1 755
VIRT-153210:53 .	polypeptide	1	755	.	+	.	ID=VIRT-
153210:53;md5=13ed02428b11466e27d0311041ac6c48
VIRT-153210:53 Coils	protein_match	335	369	.	+	.	date=23-06-2020;Target=VIRT-153210:53
335 369;ID=match$1_335_369;Name=Coil;status=T
VIRT-153210:53 Coils	protein_match	402	422	.	+	.	date=23-06-2020;Target=VIRT-153210:53
402 422;ID=match$2_402_422;Name=Coil;status=T
VIRT-153210:53 MobiDBLite	protein_match	710	755	.	+	.	date=23-06-2020;Target=VIRT-
153210:53 710 755;ID=match$3_710_755;signature_desc=consensus disorder prediction;Name=mobidb-lite;status=T
VIRT-153210:53 MobiDBLite	protein_match	150	186	.	+	.	date=23-06-2020;Target=VIRT-
153210:53 150 186;ID=match$4_150_186;signature_desc=consensus disorder prediction;Name=mobidb-lite;status=T
VIRT-153210:53 Coils	protein_match	275	295	.	+	.	date=23-06-2020;Target=VIRT-153210:53
275 295;ID=match$5_275_295;Name=Coil;status=T
VIRT-153210:53 MobiDBLite	protein_match	126	249	.	+	.	date=23-06-2020;Target=VIRT-
153210:53 126 249;ID=match$6_126_249;signature_desc=consensus disorder prediction;Name=mobidb-lite;status=T
VIRT-153210:53 Coils	protein_match	430	471	.	+	.	date=23-06-2020;Target=VIRT-153210:53
430 471;ID=match$7_430_471;Name=Coil;status=T
VIRT-153210:53 MobiDBLite	protein_match	518	550	.	+	.	date=23-06-2020;Target=VIRT-
153210:53 518 550;ID=match$8_518_550;signature_desc=consensus disorder prediction;Name=mobidb-lite;status=T
VIRT-153210:53 MobiDBLite	protein_match	592	621	.	+	.	date=23-06-2020;Target=VIRT-
153210:53 592 621;ID=match$9_592_621;signature_desc=consensus disorder prediction;Name=mobidb-lite;status=T
VIRT-153210:53 MobiDBLite	protein_match	506	636	.	+	.	date=23-06-2020;Target=VIRT-
153210:53 506 636;ID=match$10_506_636;signature_desc=consensus disorder prediction;Name=mobidb-lite;status=T
VIRT-153210:53 MobiDBLite	protein_match	722	743	.	+	.	date=23-06-2020;Target=VIRT-
153210:53 722 743;ID=match$11_722_743;signature_desc=consensus disorder prediction;Name=mobidb-lite;status=T
##FASTA
>VIRT-153210:53
MTDERALESNFDRNGRIVQFQEKILNKNVSNRPTPLFIDDRQHKRFDVAKLSAAGDQAKV
ETVVDAGLDKFYENDAKCEDTREPLGSEDTLSASIPCPLLVDHSPMDESESTEDEVDDDV
GIDVTYKNNRQCDNPSSLVVIGRGIRLHTPPRDGPIEKEEGSQRPQPEGEESDASFPARE
DVQRDPEIFLPSPSTPPETKRAGSSRRPTLSPVIEEDVREYMQTWRYEDEQDEREDDDDS
GAGDSTSNVVESVINVGSDDDYANEMSRVLSSAAVDAALAEIDNEEEKIRQPKNQLTGCN
EDLDDSCDDIVIILPEAQELNYYNVPVGCSLEDSLQDAANARAQLDAQADFLENSLNCLE
ITLDRMSLNEAVMPDGSVVFLRDQLRKAKTAELETPLAQIQRELYQEQIKSYNAANEEMR
TAVTQATLQEQALRGQARQVQQTYNGLEQNIANLSNELVGLQNRQHHLQNAMMEERVAKE
AAQREHERIYRRSSSDVIDANNDILRESHIRRPRQPEWISDQSASTSSSEDIPTSGPYQH
DGGQRWWSQRGSRQREYDYIQPPPWRRGYRPSGGRRFGFGQRPNWRSNHGNWRWSETRDG
PRPWERRRPPPPPHVHRPFVDRPRPPPPAPFTPPFKRETEVMLTQYFHEMGRVENLLRER
GFAKGPAATAPVTPYRPHPTRYQRYSGCHRGGWDQRMGSRSLPSMRGYGFPYRERSQRST
```

```
DWFDTSRSQTNNQMSQRSSEEEMPATTVVHAAASA
>match$1_335_369
LQDAANARAQLDAQADFLENSLNCLEITLDRMSLN
>match$2_402_422
RELYQEQIKSYNAANEEMRTA
>match$3_710_755
FPYRERSQRSTDWFDTSRSQTNNQMSQRSSEEEMPATTVVHAAASA
>match$4_150_186
PPRDGPIEKEEGSQRPQPEGEESDASFPAREDVQRDP
>match$5_275_295
VDAALAEIDNEEEKIRQPKNQ
>match$6_126_249
YKNNRQCDNPSSLVVIGRGIRLHTPPRDGPIEKEEGSQRPQPEGEESDASFPAREDVQRD
PEIFLPSPSTPPETKRAGSSRRPTLSPVIEEDVREYMQTWRYEDEQDEREDDDDSGAGDS
TSNV
>match$7_430_471
EQALRGQARQVQQTYNGLEQNIANLSNELVGLQNRQHHLQNA
>match$8_518_550
WISDQSASTSSSEDIPTSGPYQHDGGQRWWSQR
>match$9_592_621
WRWSETRDGPRPWERRRPPPPPHVHRPFVD
>match$10_506_636
RESHIRRPRQPEWISDQSASTSSSEDIPTSGPYQHDGGQRWWSQRGSRQREYDYIQPPPW
RRGYRPSGGRRFGFGQRPNWRSNHGNWRWSETRDGPRPWERRRPPPPPHVHRPFVDRPRP
PPPAPFTPPFK
>match$11_722_743
WFDTSRSQTNNQMSQRSSEEEM
```

### Novel Genes Ovary

### L_var_09132_ovary- transmembrane protein with glycosyltransferase domain

```
##interproscan-version 5.45-80.0
##sequence-region SPU_012963.3a 1 480
SPU_012963.3a	.	polypeptide	1	480	.	+	.
	ID=SPU_012963.3a;md5=d43fc63e7c6bf775a4fb0ab62061ac65
SPU_012963.3a	Phobius	protein_match	45	480	.	+	.	date=24-06-2020;Target=SPU_012963.3a 45 480;ID=match$1_45_480;signature_desc=Region of a membrane-bound protein predicted to be outside the membrane, in the extracellular region.;Name=NON_CYTOPLASMIC_DOMAIN;status=T
SPU_012963.3a	PANTHER	protein_match	50	474	3.9E-139	+	.	date=24-06-2020;Target=SPU_012963.3a 50 474;ID=match$2_50_474;Name=PTHR21461;status=T
SPU_012963.3a	TMHMM	protein_match	23	45	.	+	.	date=24-06-2020;Target=SPU_012963.3a 23 45;ID=match$3_23_45;signature_desc=Region of a membrane-bound protein predicted to be embedded in the membrane.;Name=TMhelix;status=T
SPU_012963.3a	PANTHER	protein_match	50	474	3.9E-139	+	.	date=24-06-2020;Target=SPU_012963.3a 50 474;ID=match$4_50_474;Name=PTHR21461:SF0;status=T
SPU_012963.3a	Phobius	protein_match	1	22	.	+	.	date=24-06-2020;Target=SPU_012963.3a 1 22;ID=match$5_1_22;signature_desc=Region of a membrane-bound protein predicted to be outside the membrane, in the cytoplasm.;Name=CYTOPLASMIC_DOMAIN;status=T
SPU_012963.3a	Phobius	protein_match	23	44	.	+	.	date=24-06-2020;Target=SPU_012963.3a 23 44;ID=match$6_23_44;signature_desc=Region of a membrane-bound protein predicted to be embedded in the membrane.;Name=TRANSMEMBRANE;status=T
SPU_012963.3a	Pfam	protein_match	225	435	3.1E-23	+	.	date=24-06-2020;Target=SPU_012963.3a 225 435;ID=match$7_225_435;signature_desc=Glycosyltransferase family 92;Name=PF01697;status=T;Dbxref="InterPro:IPR008166"
##FASTA
>SPU_012963.3a
MLSRAARLGALRIGRILSCKRRFWPVLAVVITIIYLIASLSFMIQDKSLSSLGFVHNSDE
QLEQLVGRTLPRVKDQPPSSCKANIEVVTHRTLGDGCFFKDRIQCEGMPTINSAFFDVVD
RNVVFIGTLFQNESWINETFICEFPDSNVSLVDPVVIDDRSMGTQPQYVFVMTCPIPDIY
FTSFLPHNLPDRLHVNLLENSTLFQSYRDVPLCKAGWNKVHYLAICTMVNNVDEYFDDWL
LYYRYMGIDHVYVYDNSKDGTLLGVLDRFISLGFVTVIPWSHTFTPTKTYLEVQIAHEND
CLWRHRHDTDWILKIDVDEFMQPMFETETRLVDFLHEFEQTLPPSIGIVRVRNWFFSRPS
SNVSKEIMSGKSVIERNPWRSPEPTREGRGREKCFIRPQWVHYFKIHAMKLGGDSVTLDP
QTEIRLVHYRSENPRHRNFRIHKFIPDNSMVNLWKAINRWVLFTKDRVVMYGYREHLYKT
>match$1_45_480
QDKSLSSLGFVHNSDEQLEQLVGRTLPRVKDQPPSSCKANIEVVTHRTLGDGCFFKDRIQ
CEGMPTINSAFFDVVDRNVVFIGTLFQNESWINETFICEFPDSNVSLVDPVVIDDRSMGT
QPQYVFVMTCPIPDIYFTSFLPHNLPDRLHVNLLENSTLFQSYRDVPLCKAGWNKVHYLA
ICTMVNNVDEYFDDWLLYYRYMGIDHVYVYDNSKDGTLLGVLDRFISLGFVTVIPWSHTF
TPTKTYLEVQIAHENDCLWRHRHDTDWILKIDVDEFMQPMFETETRLVDFLHEFEQTLPP
SIGIVRVRNWFFSRPSSNVSKEIMSGKSVIERNPWRSPEPTREGRGREKCFIRPQWVHYF
```

```
KIHAMKLGGDSVTLDPQTEIRLVHYRSENPRHRNFRIHKFIPDNSMVNLWKAINRWVLFT
KDRVVMYGYREHLYKT
>match$2_50_474
SSLGFVHNSDEQLEQLVGRTLPRVKDQPPSSCKANIEVVTHRTLGDGCFFKDRIQCEGMP
TINSAFFDVVDRNVVFIGTLFQNESWINETFICEFPDSNVSLVDPVVIDDRSMGTQPQYV
FVMTCPIPDIYFTSFLPHNLPDRLHVNLLENSTLFQSYRDVPLCKAGWNKVHYLAICTMV
NNVDEYFDDWLLYYRYMGIDHVYVYDNSKDGTLLGVLDRFISLGFVTVIPWSHTFTPTKT
YLEVQIAHENDCLWRHRHDTDWILKIDVDEFMQPMFETETRLVDFLHEFEQTLPPSIGIV
RVRNWFFSRPSSNVSKEIMSGKSVIERNPWRSPEPTREGRGREKCFIRPQWVHYFKIHAM
KLGGDSVTLDPQTEIRLVHYRSENPRHRNFRIHKFIPDNSMVNLWKAINRWVLFTKDRVV
MYGYR
>match$3_23_45
FWPVLAVVITIIYLIASLSFMIQ
>match$4_50_474
SSLGFVHNSDEQLEQLVGRTLPRVKDQPPSSCKANIEVVTHRTLGDGCFFKDRIQCEGMP
TINSAFFDVVDRNVVFIGTLFQNESWINETFICEFPDSNVSLVDPVVIDDRSMGTQPQYV
FVMTCPIPDIYFTSFLPHNLPDRLHVNLLENSTLFQSYRDVPLCKAGWNKVHYLAICTMV
NNVDEYFDDWLLYYRYMGIDHVYVYDNSKDGTLLGVLDRFISLGFVTVIPWSHTFTPTKT
YLEVQIAHENDCLWRHRHDTDWILKIDVDEFMQPMFETETRLVDFLHEFEQTLPPSIGIV
RVRNWFFSRPSSNVSKEIMSGKSVIERNPWRSPEPTREGRGREKCFIRPQWVHYFKIHAM
KLGGDSVTLDPQTEIRLVHYRSENPRHRNFRIHKFIPDNSMVNLWKAINRWVLFTKDRVV
MYGYR
>match$5_1_22
MLSRAARLGALRIGRILSCKRR
>match$6_23_44
FWPVLAVVITIIYLIASLSFMI
>match$7_225_435
ICTMVNNVDEYFDDWLLYYRYMGIDHVYVYDNSKDGTLLGVLDRFISLGFVTVIPWSHTF
TPTKTYLEVQIAHENDCLWRHRHDTDWILKIDVDEFMQPMFETETRLVDFLHEFEQTLPP
SIGIVRVRNWFFSRPSSNVSKEIMSGKSVIERNPWRSPEPTREGRGREKCFIRPQWVHYF
KIHAMKLGGDSVTLDPQTEIRLVHYRSENPR
```

### L_var_11906_ovary- 7-pass transmembrane (GPCR)

```
##interproscan-version 5.45-80.0
##sequence-region SPU_007031.3a 1 182
SPU_007031.3a	.	polypeptide	1	182	.	+	.
	ID=SPU_007031.3a;md5=867e649d6ea72b557afd91907ed367b3
SPU_007031.3a	TMHMM	protein_match	92	114	.	+	.	date=24-06-2020;Target=SPU_007031.3a 92 114;ID=match$1_92_114;signature_desc=Region of a membrane-bound protein predicted to be embedded in the membrane.;Name=TMhelix;status=T
SPU_007031.3a	Phobius	protein_match	148	182	.	+	.	date=24-06-2020;Target=SPU_007031.3a 148 182;ID=match$2_148_182;signature_desc=Region of a membrane-bound protein predicted to be outside the membrane, in the cytoplasm.;Name=CYTOPLASMIC_DOMAIN;status=T
SPU_007031.3a	TMHMM	protein_match	124	146	.	+	.	date=24-06-2020;Target=SPU_007031.3a 124 146;ID=match$3_124_146;signature_desc=Region of a membrane-bound protein predicted to be embedded in the membrane.;Name=TMhelix;status=T
SPU_007031.3a	Phobius	protein_match	7	35	.	+	.	date=24-06-2020;Target=SPU_007031.3a 7 35;ID=match$4_7_35;signature_desc=Region of a membrane-bound protein predicted to be embedded in the membrane.;Name=TRANSMEMBRANE;status=T
SPU_007031.3a	TMHMM	protein_match	50	72	.	+	.	date=24-06-2020;Target=SPU_007031.3a 50 72;ID=match$5_50_72;signature_desc=Region of a membrane-bound protein predicted to be embedded in the membrane.;Name=TMhelix;status=T
SPU_007031.3a	Phobius	protein_match	87	117	.	+	.	date=24-06-2020;Target=SPU_007031.3a 87 117;ID=match$6_87_117;signature_desc=Region of a membrane-bound protein predicted to be embedded in the membrane.;Name=TRANSMEMBRANE;status=T
SPU_007031.3a	Phobius	protein_match	118	122	.	+	.	date=24-06-2020;Target=SPU_007031.3a 118 122;ID=match$7_118_122;signature_desc=Region of a membrane-bound protein predicted to be outside the membrane, in the extracellular region.;Name=NON_CYTOPLASMIC_DOMAIN;status=T
SPU_007031.3a	Phobius	protein_match	55	75	.	+	.	date=24-06-2020;Target=SPU_007031.3a 55 75;ID=match$8_55_75;signature_desc=Region of a membrane-bound protein predicted to be embedded in the membrane.;Name=TRANSMEMBRANE;status=T
SPU_007031.3a	Phobius	protein_match	123	147	.	+	.	date=24-06-2020;Target=SPU_007031.3a 123 147;ID=match$9_123_147;signature_desc=Region of a membrane-bound protein predicted to be embedded in the membrane.;Name=TRANSMEMBRANE;status=T
SPU_007031.3a	Phobius	protein_match	76	86	.	+	.	date=24-06-2020;Target=SPU_007031.3a 76 86;ID=match$10_76_86;signature_desc=Region of a membrane-bound protein predicted to be outside the membrane, in the cytoplasm.;Name=CYTOPLASMIC_DOMAIN;status=T
SPU_007031.3a	TMHMM	protein_match	13	35	.	+	.	date=24-06-2020;Target=SPU_007031.3a 13 35;ID=match$11_13_35;signature_desc=Region of a membrane-bound protein predicted to be embedded in the membrane.;Name=TMhelix;status=T
SPU_007031.3a	Phobius	protein_match	1	6	.	+	.	date=24-06-2020;Target=SPU_007031.3a 1 6;ID=match$12_1_6;signature_desc=Region of a membrane-bound protein predicted to be outside the membrane, in the cytoplasm.;Name=CYTOPLASMIC_DOMAIN;status=T
```

```
SPU_007031.3a  Phobius protein_match  36      54      .       +       .       date=24-06-2020;Target=SPU_007031.3a
36 54;ID=match$13_36_54;signature_desc=Region of a membrane-bound protein predicted to be outside the membrane,
in the extracellular region.;Name=NON_CYTOPLASMIC_DOMAIN;status=T
##FASTA
>SPU_007031.3a
MGIRRQFIGFTGAVQVLLTLVMFAMGATLSDIAIVQGVVQGSADDSSIDSMAIGVPLWGG
IVILMCGAGNIVEAFTARSSRKRKNDLLPCSIGAFLANLIAFVVSGIIIGIFSWSIYEVA
STFIIAVSTVLLSASIIFLMSMLAMFVDCMTVCLSTGPSQPRPYIVDYDHGPPRGPVHVY
KA
>match$1_92_114
IGAFLANLIAFVVSGIIIGIFSW
>match$2_148_182
DCMTVCLSTGPSQPRPYIVDYDHGPPRGPVHVYKA
>match$3_124_146
IIAVSTVLLSASIIFLMSMLAMF
>match$4_7_35
FIGFTGAVQVLLTLVMFAMGATLSDIAIV
>match$5_50_72
SMAIGVPLWGGIVILMCGAGNIV
>match$6_87_117
LLPCSIGAFLANLIAFVVSGIIIGIFSWSIY
>match$7_118_122
EVAST
>match$8_55_75
VPLWGGIVILMCGAGNIVEAF
>match$9_123_147
FIIAVSTVLLSASIIFLMSMLAMFV
>match$10_76_86
TARSSRKRKND
>match$11_13_35
AVQVLLTLVMFAMGATLSDIAIV
>match$12_1_6
MGIRRQ
>match$13_36_54
QGVVQGSADDSSIDSMAIG
```

### L_var_17102_ovary- transmembrane protein similar to GPCR

```
##interproscan-version 5.45-80.0
##sequence-region VIRT-60199:53 1 72
VIRT-60199:53  .       polypeptide    1       72      .       +       .       ID=VIRT-
60199:53;md5=b5c87b0bf56fd3369992d9dd879105f6
VIRT-60199:53  Phobius protein_match  31      48      .       +       .       date=24-06-2020;Target=VIRT-60199:53
31 48;ID=match$1_31_48;signature_desc=Region of a membrane-bound protein predicted to be embedded in the
membrane.;Name=TRANSMEMBRANE;status=T
VIRT-60199:53  Phobius protein_match  26      30      .       +       .       date=24-06-2020;Target=VIRT-60199:53
26 30;ID=match$2_26_30;signature_desc=Region of a membrane-bound protein predicted to be outside the membrane,
in the extracellular region.;Name=NON_CYTOPLASMIC_DOMAIN;status=T
VIRT-60199:53  TMHMM   protein_match  30      47      .       +       .       date=24-06-2020;Target=VIRT-60199:53
30 47;ID=match$3_30_47;signature_desc=Region of a membrane-bound protein predicted to be embedded in the
membrane.;Name=TMhelix;status=T
VIRT-60199:53  Phobius protein_match  49      72      .       +       .       date=24-06-2020;Target=VIRT-60199:53
49 72;ID=match$4_49_72;signature_desc=Region of a membrane-bound protein predicted to be outside the membrane,
in the cytoplasm.;Name=CYTOPLASMIC_DOMAIN;status=T
VIRT-60199:53  Phobius protein_match  1       6       .       +       .       date=24-06-2020;Target=VIRT-60199:53
1 6;ID=match$5_1_6;signature_desc=Region of a membrane-bound protein predicted to be outside the membrane, in
the cytoplasm.;Name=CYTOPLASMIC_DOMAIN;status=T
VIRT-60199:53  Phobius protein_match  7       25      .       +       .       date=24-06-2020;Target=VIRT-60199:53
7 25;ID=match$6_7_25;signature_desc=Region of a membrane-bound protein predicted to be embedded in the
membrane.;Name=TRANSMEMBRANE;status=T
##FASTA
>VIRT-60199:53
MESEETHRYSLVMSKLFFLFIHQSFDACSVYFQHASLTSGLFIVYLLAERPCRYVDKVFD
DLKPMRTWWDHI
>match$1_31_48
YFQHASLTSGLFIVYLLA
>match$2_26_30
DACSV
>match$3_30_47
VYFQHASLTSGLFIVYLL
>match$4_49_72
ERPCRYVDKVFDDLKPMRTWWDHI
>match$5_1_6
```

```
MESEET
>match$6_7_25
HRYSLVMSKLFFLFIHQSF
```

### L_var_18001_ovary- membrane bound signaling peptide

```
##interproscan-version 5.45-80.0
##sequence-region SPU_009951.3a 1 147
SPU_009951.3a	.	polypeptide	1	147	.	+	.	ID=SPU_009951.3a;md5=b8297ea0e716e90960fb2f584b206605
SPU_009951.3a	Phobius	protein_match	16	20	.	+	.	date=24-06-2020;Target=SPU_009951.3a 16 20;ID=match$1_16_20;signature_desc=C-terminal region of a signal peptide.;Name=SIGNAL_PEPTIDE_C_REGION;status=T
SPU_009951.3a	Phobius	protein_match	21	147	.	+	.	date=24-06-2020;Target=SPU_009951.3a 21 147;ID=match$2_21_147;signature_desc=Region of a membrane-bound protein predicted to be outside the membrane, in the extracellular region.;Name=NON_CYTOPLASMIC_DOMAIN;status=T
SPU_009951.3a	SignalP_GRAM_NEGATIVE	protein_match	1	19	.	+	.	date=24-06-2020;Target=SPU_009951.3a 1 19;ID=match$3_1_19;Name=SignalP-noTM;status=T
SPU_009951.3a	Phobius	protein_match	1	20	.	+	.	date=24-06-2020;Target=SPU_009951.3a 1 20;ID=match$4_1_20;signature_desc=Signal peptide region;Name=SIGNAL_PEPTIDE;status=T
SPU_009951.3a	Phobius	protein_match	1	3	.	+	.	date=24-06-2020;Target=SPU_009951.3a 1 3;ID=match$5_1_3;signature_desc=N-terminal region of a signal peptide.;Name=SIGNAL_PEPTIDE_N_REGION;status=T
SPU_009951.3a	Phobius	protein_match	4	15	.	+	.	date=24-06-2020;Target=SPU_009951.3a 4 15;ID=match$6_4_15;signature_desc=Hydrophobic region of a signal peptide.;Name=SIGNAL_PEPTIDE_H_REGION;status=T
SPU_009951.3a	SignalP_GRAM_POSITIVE	protein_match	1	21	.	+	.	date=24-06-2020;Target=SPU_009951.3a 1 21;ID=match$7_1_21;Name=SignalP-TM;status=T
SPU_009951.3a	SignalP_EUK	protein_match	1	20	.	+	.	date=24-06-2020;Target=SPU_009951.3a 1 20;ID=match$8_1_20;Name=SignalP-noTM;status=T
##FASTA
>SPU_009951.3a
MARVAFFVVFVIVCAVSVASSPTEREISEKQIKEKGRMLLANFRQKARTLLTMKVAEWKT
ADQIETNVDILERFQHVEKEAEYEIKQHIVDAINGNEPVVDPDIPPHSEEIATLFGEALF
ATTYAEQSTKEDSMIQILKDILENKLH
>match$1_16_20
VSVAS
>match$2_21_147
SPTEREISEKQIKEKGRMLLANFRQKARTLLTMKVAEWKTADQIETNVDILERFQHVEKE
AEYEIKQHIVDAINGNEPVVDPDIPPHSEEIATLFGEALFATTYAEQSTKEDSMIQILKD
ILENKLH
>match$3_1_19
MARVAFFVVFVIVCAVSVA
>match$4_1_20
MARVAFFVVFVIVCAVSVAS
>match$5_1_3
MAR
>match$6_4_15
VAFFVVFVIVCA
>match$7_1_21
MARVAFFVVFVIVCAVSVASS
>match$8_1_20
MARVAFFVVFVIVCAVSVAS
```

### L_var_18708_ovary 7-pass transmembrane (GPCR)

```
##interproscan-version 5.45-80.0
##sequence-region SPU_010643.1 1 262
SPU_010643.1	.	polypeptide	1	262	.	+	.	ID=SPU_010643.1;md5=6610b5b723df12faba160a6bd63a4320
SPU_010643.1	Phobius	protein_match	16	20	.	+	.	date=24-06-2020;Target=SPU_010643.1 16 20;ID=match$1_16_20;signature_desc=C-terminal region of a signal peptide.;Name=SIGNAL_PEPTIDE_C_REGION;status=T
SPU_010643.1	Phobius	protein_match	1	20	.	+	.	date=24-06-2020;Target=SPU_010643.1 1 20;ID=match$2_1_20;signature_desc=Signal peptide region;Name=SIGNAL_PEPTIDE;status=T
SPU_010643.1	Phobius	protein_match	215	241	.	+	.	date=24-06-2020;Target=SPU_010643.1 215 241;ID=match$3_215_241;signature_desc=Region of a membrane-bound protein predicted to be embedded in the membrane.;Name=TRANSMEMBRANE;status=T
SPU_010643.1	Phobius	protein_match	242	262	.	+	.	date=24-06-2020;Target=SPU_010643.1 242 262;ID=match$4_242_262;signature_desc=Region of a membrane-bound protein predicted to be outside the membrane, in the cytoplasm.;Name=CYTOPLASMIC_DOMAIN;status=T
SPU_010643.1	Gene3D	protein_match	21	125	1.6E-30	+	.	date=24-06-2020;Target=SPU_010643.1 21 125;ID=match$5_21_125;Name=G3DSA:2.60.40.2140;status=T;Dbxref="InterPro:IPR043030"
SPU_010643.1	SignalP_EUK	protein_match	1	20	.	+	.	date=24-06-2020;Target=SPU_010643.1 1 20;ID=match$6_1_20;Name=SignalP-noTM;status=T
```

```
SPU_010643.1	Pfam	protein_match	24	125	2.8E-24	+	.	date=24-06-2020;Target=SPU_010643.1 24 125;Ontology_term="GO:0030246";ID=match$7_24_125;signature_desc=Carbohydrate binding domain (family 32);Name=PF15886;status=T;Dbxref="InterPro:IPR031756"
SPU_010643.1	Phobius	protein_match	7	15	.	+	.	date=24-06-2020;Target=SPU_010643.1 7 15;ID=match$8_7_15;signature_desc=Hydrophobic region of a signal peptide.;Name=SIGNAL_PEPTIDE_H_REGION;status=T
SPU_010643.1	Phobius	protein_match	21	214	.	+	.	date=24-06-2020;Target=SPU_010643.1 21 214;ID=match$9_21_214;signature_desc=Region of a membrane-bound protein predicted to be outside the membrane, in the extracellular region.;Name=NON_CYTOPLASMIC_DOMAIN;status=T
SPU_010643.1	TMHMM	protein_match	215	237	.	+	.	date=24-06-2020;Target=SPU_010643.1 215 237;ID=match$10_215_237;signature_desc=Region of a membrane-bound protein predicted to be embedded in the membrane.;Name=TMhelix;status=T
SPU_010643.1	MobiDBLite	protein_match	137	166	.	+	.	date=24-06-2020;Target=SPU_010643.1 137 166;ID=match$11_137_166;signature_desc=consensus disorder prediction;Name=mobidb-lite;status=T
SPU_010643.1	Phobius	protein_match	1	6	.	+	.	date=24-06-2020;Target=SPU_010643.1 1 6;ID=match$12_1_6;signature_desc=N-terminal region of a signal peptide.;Name=SIGNAL_PEPTIDE_N_REGION;status=T
SPU_010643.1	MobiDBLite	protein_match	167	181	.	+	.	date=24-06-2020;Target=SPU_010643.1 167 181;ID=match$13_167_181;signature_desc=consensus disorder prediction;Name=mobidb-lite;status=T
SPU_010643.1	MobiDBLite	protein_match	120	198	.	+	.	date=24-06-2020;Target=SPU_010643.1 120 198;ID=match$14_120_198;signature_desc=consensus disorder prediction;Name=mobidb-lite;status=T
##FASTA
>SPU_010643.1
MVRLRPVVALLPLILVSINAYDMKNPEISLLTTGGIRFAYPDEPGITLVAFHYSINTPLS
GVNVGQYNYDVTTKTGAYFVHENTEVDVKKGDVVNYWVYVNYYGPGYQLLEQSWTASEAP
ATVSPASNPPASNPPASNRPATESPATEPPATNPRASNRPATNPPATEPRATNPPGVTNP
PKPVTATIEMKSNRNHQSKNDLISDEKTMRISSSYWLAFGLVVVVTIIDAYGAAILTTLV
LTMGNDYGNADPVVGEELRKSR
>match$1_16_20
VSINA
>match$2_1_20
MVRLRPVVALLPLILVSINA
>match$3_215_241
YWLAFGLVVVVTIIDAYGAAILTTLVL
>match$4_242_262
TMGNDYGNADPVVGEELRKSR
>match$5_21_125
YDMKNPEISLLTTGGIRFAYPDEPGITLVAFHYSINTPLSGVNVGQYNYDVTTKTGAYFV
HENTEVDVKKGDVVNYWVYVNYYGPGYQLLEQSWTASEAPATVSP
>match$6_1_20
MVRLRPVVALLPLILVSINA
>match$7_24_125
KNPEISLLTTGGIRFAYPDEPGITLVAFHYSINTPLSGVNVGQYNYDVTTKTGAYFVHEN
TEVDVKKGDVVNYWVYVNYYGPGYQLLEQSWTASEAPATVSP
>match$8_7_15
VVALLPLIL
>match$9_21_214
YDMKNPEISLLTTGGIRFAYPDEPGITLVAFHYSINTPLSGVNVGQYNYDVTTKTGAYFV
HENTEVDVKKGDVVNYWVYVNYYGPGYQLLEQSWTASEAPATVSPASNPPASNPPASNRP
ATESPATEPPATNPRASNRPATNPPATEPRATNPPGVTNPPKPVTATIEMKSNRNHQSKN
DLISDEKTMRISSS
>match$10_215_237
YWLAFGLVVVVTIIDAYGAAILT
>match$11_137_166
SNRPATESPATEPPATNPRASNRPATNPPA
>match$12_1_6
MVRLRP
>match$13_167_181
TEPRATNPPGVTNPP
>match$14_120_198
PATVSPASNPPASNPPASNRPATESPATEPPATNPRASNRPATNPPATEPRATNPPGVTN
PPKPVTATIEMKSNRNHQS
```

### L_var_19833_ovary- transmembrane protein with prokaryotic lipoprotein attachment domain

```
##interproscan-version 5.45-80.0
##sequence-region VIRT-169181:53 1 85
VIRT-169181:53	.	polypeptide	1	85	.	+	.	ID=VIRT-169181:53;md5=96bf3833f26647f0c321d62e89b11667
```

VIRT-169181:53 Phobius protein_match 20 40 . + . date=24-06-2020;Target=VIRT-169181:53 20 40;ID=match$1_20_40;signature_desc=Region of a membrane-bound protein predicted to be embedded in the membrane.;Name=TRANSMEMBRANE;status=T
VIRT-169181:53 TMHMM protein_match 10 32 . + . date=24-06-2020;Target=VIRT-169181:53 10 32;ID=match$2_10_32;signature_desc=Region of a membrane-bound protein predicted to be embedded in the membrane.;Name=TMhelix;status=T
VIRT-169181:53 Phobius protein_match 1 19 . + . date=24-06-2020;Target=VIRT-169181:53 1 19;ID=match$3_1_19;signature_desc=Region of a membrane-bound protein predicted to be outside the membrane, in the extracellular region.;Name=NON_CYTOPLASMIC_DOMAIN;status=T
VIRT-169181:53 Phobius protein_match 41 85 . + . date=24-06-2020;Target=VIRT-169181:53 41 85;ID=match$4_41_85;signature_desc=Region of a membrane-bound protein predicted to be outside the membrane, in the cytoplasm.;Name=CYTOPLASMIC_DOMAIN;status=T
VIRT-169181:53 ProSiteProfiles protein_match 1 19 5.0 + . date=24-06-2020;Target=VIRT-169181:53 1 19;ID=match$5_1_19;signature_desc=Prokaryotic membrane lipoprotein lipid attachment site profile.;Name=PS51257;status=T
##FASTA
>VIRT-169181:53
MKCGYLCFSSHFGSASYSCWYFFSPFIHLLGYQFTSFMLFTRCSFDKFLFLLTYSRNRYE
EHNYCHLAVYSNAVWHAGTYRSQRI
>match$1_20_40
WYFFSPFIHLLGYQFTSFMLF
>match$2_10_32
SHFGSASYSCWYFFSPFIHLLGY
>match$3_1_19
MKCGYLCFSSHFGSASYSC
>match$4_41_85
TRCSFDKFLFLLTYSRNRYEEHNYCHLAVYSNAVWHAGTYRSQRI
>match$5_1_19
MKCGYLCFSSHFGSASYSC

### L_var_24614_ovary- sensor for hormone

##interproscan-version 5.45-80.0
##sequence-region SPU_012915.3a 1 192
SPU_012915.3a . polypeptide 1 192 . + . ID=SPU_012915.3a;md5=a5d608018603ff2a0554fb5b93428ac5
SPU_012915.3a PANTHER protein_match 11 180 1.1E-42 + . date=24-06-2020;Target=SPU_012915.3a 11 180;ID=match$1_11_180;Name=PTHR14315;status=T;Dbxref="InterPro:IPR009786","Reactome:R-HSA-200425"
SPU_012915.3a PANTHER protein_match 11 180 1.1E-42 + . date=24-06-2020;Target=SPU_012915.3a 11 180;ID=match$2_11_180;Name=PTHR14315:SF17;status=T
SPU_012915.3a Pfam protein_match 25 179 1.2E-23 + . date=24-06-2020;Target=SPU_012915.3a 25 179;ID=match$3_25_179;signature_desc=Thyroid hormone-inducible hepatic protein Spot 14;Name=PF07084;status=T;Dbxref="InterPro:IPR009786","Reactome:R-HSA-200425"
##FASTA
>SPU_012915.3a
MSDFEFTTLCHSMMMELQQRDNEKAPPSQSILGIMKNFIDSVNEMDETVLIPSRLMDINM
DNCGSMLMDNDDSTRTSMSVIPTSTSALPSSTKEVSNDQVSLHTYYSMLKAVKRELARGP
VNESELSELEDETLDEESARLQAETARAFREHLRGLFSILSHLTDVSKHLTTIYQQETGD
HQSCLKPKSFHV
>match$1_11_180
HSMMMELQQRDNEKAPPSQSILGIMKNFIDSVNEMDETVLIPSRLMDINMDNCGSMLMDN
DDSTRTSMSVIPTSTSALPSSTKEVSNDQVSLHTYYSMLKAVKRELARGPVNESELSELE
DETLDEESARLQAETARAFREHLRGLFSILSHLTDVSKHLTTIYQQETGD
>match$2_11_180
HSMMMELQQRDNEKAPPSQSILGIMKNFIDSVNEMDETVLIPSRLMDINMDNCGSMLMDN
DDSTRTSMSVIPTSTSALPSSTKEVSNDQVSLHTYYSMLKAVKRELARGPVNESELSELE
DETLDEESARLQAETARAFREHLRGLFSILSHLTDVSKHLTTIYQQETGD
>match$3_25_179
APPSQSILGIMKNFIDSVNEMDETVLIPSRLMDINMDNCGSMLMDNDDSTRTSMSVIPTS
TSALPSSTKEVSNDQVSLHTYYSMLKAVKRELARGPVNESELSELEDETLDEESARLQAE
TARAFREHLRGLFSILSHLTDVSKHLTTIYQQETG

### L_var_24625_ovary- novel methyltransferase

##interproscan-version 5.45-80.0
##sequence-region SPU_027593.3a 1 222
SPU_027593.3a . polypeptide 1 222 . + . ID=SPU_027593.3a;md5=aaab5605c92f041489929aaac9cb4170
SPU_027593.3a PANTHER protein_match 21 182 2.2E-36 + . date=24-06-2020;Target=SPU_027593.3a 21 182;ID=match$1_21_182;Name=PTHR43591:SF29;status=T
SPU_027593.3a Pfam protein_match 71 161 1.1E-10 + . date=24-06-2020;Target=SPU_027593.3a 71 161;ID=match$2_71_161;signature_desc=Methyltransferase domain;Name=PF13649;status=T;Dbxref="InterPro:IPR041698"

```
SPU_027593.3a	PANTHER	protein_match	21	182	2.2E-36	+	.	date=24-06-2020;Target=SPU_027593.3a 21 182;ID=match$3_21_182;Name=PTHR43591;status=T
SPU_027593.3a	Gene3D	protein_match	24	207	6.7E-21	+	.	date=24-06-2020;Target=SPU_027593.3a 24 207;ID=match$4_24_207;Name=G3DSA:3.40.50.150;status=T
SPU_027593.3a	CDD	protein_match	70	167	4.67545E-9	+	.	date=24-06-2020;Target=SPU_027593.3a 70 167;ID=match$5_70_167;signature_desc=AdoMet_MTases;Name=cd02440;status=T
SPU_027593.3a	Coils	protein_match	27	47	.	+	.	date=24-06-2020;Target=SPU_027593.3a 27 47;ID=match$6_27_47;Name=Coil;status=T
SPU_027593.3a	SUPERFAMILY	protein_match	19	171	7.47E-24	+	.	date=24-06-2020;Target=SPU_027593.3a 19 171;ID=match$7_19_171;Name=SSF53335;status=T;Dbxref="InterPro:IPR029063"
##FASTA
>SPU_027593.3a
MSMTTGTSEDDLHWKNMYSASSGDLDLKRLEQAYKGWSETYDEDNEQMLYKGPHLAAQKL
SKLMPDKSKKILDVACGTGLVGKELHSQGYVNIDGVDLVQDMLTHAEQTGVYSRLEACDV
IGQGLSCQDGTYEAIVCVGSFNQGCVTQAVFPELLRVAKKGCIILIVMREAFVYTCPAFK
NSQLDTDILMRAQNEIWKYILRETVPKYIGELTGLLYAMVVQ
>match$1_21_182
SSGDLDLKRLEQAYKGWSETYDEDNEQMLYKGPHLAAQKLSKLMPDKSKKILDVACGTGL
VGKELHSQGYVNIDGVDLVQDMLTHAEQTGVYSRLEACDVIGQGLSCQDGTYEAIVCVGS
FNQGCVTQAVFPELLRVAKKGCIILIVMREAFVYTCPAFKNS
>match$2_71_161
ILDVACGTGLVGKELHSQGYVNIDGVDLVQDMLTHAEQTGVYSRLEACDVIGQGLSCQDG
TYEAIVCVGSFNQGCVTQAVFPELLRVAKKG
>match$3_21_182
SSGDLDLKRLEQAYKGWSETYDEDNEQMLYKGPHLAAQKLSKLMPDKSKKILDVACGTGL
VGKELHSQGYVNIDGVDLVQDMLTHAEQTGVYSRLEACDVIGQGLSCQDGTYEAIVCVGS
FNQGCVTQAVFPELLRVAKKGCIILIVMREAFVYTCPAFKNS
>match$4_24_207
DLDLKRLEQAYKGWSETYDEDNEQMLYKGPHLAAQKLSKLMPDKSKKILDVACGTGLVGK
ELHSQGYVNIDGVDLVQDMLTHAEQTGVYSRLEACDVIGQGLSCQDGTYEAIVCVGSFNQ
GCVTQAVFPELLRVAKKGCIILIVMREAFVYTCPAFKNSQLDTDILMRAQNEIWKYILRE
TVPK
>match$5_70_167
KILDVACGTGLVGKELHSQGYVNIDGVDLVQDMLTHAEQTGVYSRLEACDVIGQGLSCQD
GTYEAIVCVGSFNQGCVTQAVFPELLRVAKKGCIILIV
>match$6_27_47
LKRLEQAYKGWSETYDEDNEQ
>match$7_19_171
SASSGDLDLKRLEQAYKGWSETYDEDNEQMLYKGPHLAAQKLSKLMPDKSKKILDVACGT
GLVGKELHSQGYVNIDGVDLVQDMLTHAEQTGVYSRLEACDVIGQGLSCQDGTYEAIVCV
GSFNQGCVTQAVFPELLRVAKKGCIILIVMREA
```

#### Supplemental Data Part 2: GAPDH Transcripts

| **LV_ID** | **Gene** | **SPU** | **Expression** |
|---|---|---|---|
| **L_var_19892** | **Sp-Gapdh** | **SPU_007155** | **Ovary** |
| **L_var_19893** | **none** | **SPU_027041** | **Ovary** |
| **L_var_19894** | **Sp-Gapdh** | **SPU_007155** | **Ovary only** |
| **L_var_22529** | **Sp-Gapdh** | **SPU_007155** | **Ovary and testis** |

```
GAPDH transcript alignments:

L_var_19893      ----------ATGAAACACGTATGCAAAG--------AGTACACAA--------------   28
L_var_22529      ------------------------------------------------------------   0
L_var_19892      GGCGGTTGTTCTAAAACATGGAAGCAATTCATTCAAGCGTTCAGGCTGTGCTTTCATCTT   60
L_var_19894      ------------------------------------------------------------   0

L_var_19893      -------CGGATAC----------------------------------------------   35
L_var_22529      ---------------------------------ATGGATATTTGGCAGACTTTCTCGG--   25
```

```
L_var_19892      TCAAGCACGCCGGACTATTCTTTGTAATGGCAAGAAAATAATGAAACGTCTTGTTCGCTC   120
L_var_19894      ------------------------------------------------------------   0

L_var_19893      ------------------------------------------------------------   35
L_var_22529      --GTTGGAGTATCTCGGGTGCAGAATGTCCTGGTCCAAAGGACAGACTGTTCAAGGACTC   83
L_var_19892      CCGCCATGTTAACCCGCGCAAATTAAATT-TACACCCTAGTCCTTCCACCCTGTAAAATC   179
L_var_19894      ------------------------------------------------------------   0

L_var_19893      ------------------------------------------------------------   35
L_var_22529      GTCTAAGGACAGGACCCTCAAA---------------------CAGGATGATCACCACCC   122
L_var_19892      ATCCAAGGTGATGACCAACAGTAGTTTTATTTTTAAGATCGTCTACGTTCCTCCACTGAC   239
L_var_19894      -------------------------------ATGTGGCCATTCGACGGTCGTGAGCAAAA   29

L_var_19893      ------------------------------------------------------------   35
L_var_22529      CATCACTTACTCTCATAACCCACAAACCAAAGCTACTCATTCACCATGGTTATCAAGATT   182
L_var_19892      TAGCTAGGCATAAAACGACTTACCAACCATCATGGTGATTAAAATTGGAATCAACGGGTT   299
L_var_19894      ACTCATCAAATTTTTCCACGACTTTCTCCGGGACCTTCTGGGGGATCACCTTGGCGGGTT   89

L_var_19893      ------------------------------------------------------------   35
L_var_22529      TGGCCGTATCGGTCGTCTGACTTTGAGGGCAGC------CCTCAAGAACCCCAGCGTACA   236
L_var_19892      TGGACGTATAGGGCGTCTGGCCCTAAGAGCTGTTCTTCGAAACGGGGCCTCCAATGTCCA   359
L_var_19894      TGGACGTATAGGGCGTCTGGCCCTAAGAGCTGTTCTTCGAAACGGGGCCTCCAACGTCCA   149

L_var_19893      ------------------------------------------------------------   35
L_var_22529      GGTTGTCGCGATCAATGATCCTTTCATTAATCCTGATTACATGGTGTACATGTTCAAGTA   296
L_var_19892      GGTAGTGGCAATCAATGATCCTTATATTGAACTTGAACACATGGCCTATCTGTTCAGGTA   419
L_var_19894      GGTAGTGGCAATCAATGATCCTTATATTGAACTTGAACACATGGCCTATCTGTTCAGGTA   209

L_var_19893      ------------------------------------------------------------   35
L_var_22529      TGACTCCACACACGGAAAGTTTGAGGGAGACGTGAGCGGTGACAATG-------------   343
L_var_19892      TGATTCTACGCATGGAAGATTTAAAGGTGATATACGCTACGACAAATCAGCAAACGTCCT   479
L_var_19894      TGATTCTACGCATGGAAGATTTAAAGGCGATATACGCTACGACAAATCAGCAAACGTCCT   269
```

### Supplemental Data Part 3: Doublesex Genes

#### LV_Dmrt1

mRNA
Best match:
**Chromosome 11**
**15,788,119-15,789,036**
**36,215,758**

>L_var_18666-RA transcript offset:0 AED:0.02 eAED:0.02 QI:0|-1|0|1|-1|1|1|0|305
ATGAGAGAAATGAAAGAAGACATAGAACGTGAGTCTCCTACAACAAGTGTCCCAGAGCTA
CCGAAGAAGATCGTGGAGAAGCATGAACCACCAGAACCCTCTGAGAAACCAACCGAAGAC
GGTGAGGAACCGAAGAAAAAGGCTAAGACTTCTGGACCTTTGGACACCTTCAATGGCATG
GCATCCAGCTTCGTGTCGCCGTTCCGTCCAGGACGGTTATCGCCGACGGAGATCCTGACA
CGGTTGTTCCCCCACCAACGCAAAGCTGTTCTGGAGTTGGTCCTGCAAGGATGCAGCGGT
GATTTGGTCAAAGCCATTGAGCACTTCCTCAGCGCCGGGGAATCCGCAAAGAACAACGGC
AGTGCCTCCTCGCGATCGGAACACGCCTCGCACTCTTCAGAAAAGGACCAGAACGAAGCG
TATCACATCCCGTCTGGACTTCCTACGGTTCCAGGACTTGGTTCTTCGATCTTCCGCACA
CCAATGCACGCTGACAAGCTAGGTGTCGGTAGCATGAAGTCGGCTTTTACTCCCCTCCCA
CCTGCTAGCTCAGCGCCTCTTCCTCTTCTCTTTTCGCACAGACCGCCAAATCCTTTCCAA
GCAGACGCTCTCCTTGGTCGTACCCCTATATTCGCCCCTTCTAGTCTGAGTGACCTACAC
GTTGGTGCTGCTGGACCTGGTCGTTTCGTATTCCCAGCCATGCACCCGCTCAATCTCTCT
GGAAAGTTAGCGGCGGCAGCCAGTGAGGGATATCCCAGGTATGTCTTTGCACCCTATGGA
ACATGTCCGCCAGACTACACGCAGTACCCGAGCCTCCATAGCACCAGAGCTCCAGCCGGA
AGCACCGGAAGTGATTCCGAAAAGAGTCCGGGTGCCATTGCTGACCTTTCGGTCACGAGC
AACGTTGATTCAGATTGA

> VIRT-4999:5'3' Frame 1, start_pos=0
MREMKEDIERESPTTSVPELPKKIVEKHEPPEPSEKPTEDGEEPKKKAKT
SGPLDTFNGMASSFVSPFRPGRLSPTEILTRLFPHQRKAVLELVLQGCSG
DLVKAIEHFLSAGESAKNNGSASSRSEHASHSSEKDQNEAYHIPSGLPTV
PGLGSSIFRTPMHADKLGVGSMKSAFTPLPPASSAPLPLLFSHRPPNPFQ
ADALLGRTPIFAPSSLSDLHVGAAGPGRFVFPAMHPLNLSGKLAAAASEG
YPRYVFAPYGTCPPDYTQYPSLHSTRAPAGSTGSDSEKSPGAIADLSVTS
NVDSD-

**>chr14**
**Length = 33,697,906**
**18,315,702-18,316,509**

Second best match:
>L_var_22210-RA transcript offset:0 AED:0.02 eAED:0.05 QI:0|0|0|1|1|1|2|0|348
TTCTCAACCGGGTCACCGAGCATGAAGGTTGGGGAGTTACCATTCACTCTCCATCGGAGA
AGCTGTCCGTGGAGCCATTCAACCACGTCCGCGACGACGACAACGGACCAAAACAGCAAC
AATCAACCATCACCTACACATCAACGGCCGTCGCCACTCACGGCGTTACTCTTCGAAGCA
AACGAGTCGTCATCGTCGCCGACGCCGTCAACGGGTCCACCGACGCCCCCCTCATCGCCA
CCCGATCCTGCACCCACATCAACGGCCACGACAACTACGCCACAGATTTTCGCGTCGCAA
CGATGTTCATGCCCGTGCAGCGACTGCGTCAAGCTAGTCCGTGACTCGACGTGTGGTATG
GATATGGACTTGGCGCGTTGGTTGACGAAATGTGCCGAATTCGGGACAGGACATACAAGC
TTAGCAAACATTAATGTACACGATAGGCAATTACTGGTATGCTCTGCCTGGAAAGACATA
GTTATCCTGTACATGGCACAGAATAAGTTCAATTTCGATGTAGTCTACTCGAGCATGCAC
GAATACCTTCAACATGTACCCATTCACCAATATGTCCAAACGTACACGACGCATCATACG
GTGGACGCTCAGTTCCAGGCGGATTTTATCGCGAAGACTGGACTAGTTTCTCAGACCTAC
GTGCATCCTCTAAGGAGAGGTAGCGACTTTGGGATACCAACCATGAGAACTGTTAACCAG
CTCCAATTCACATTATCAAAGCTCAACCAGATGCAACTTGACTCAAAGGAATATGCCGTG
TTGAGGATACTGCTCCTCTTGAATTCCGATCTGCATTCCTTGACGGATAGCCGAGCGATA
GAGGTTGCCAGCGAGAAGGTCCACACGGCCCTGATGGAGTATGAGACCCTCAAATACCCC
GATGATCCTCTTCGGCTGTCCCACATGCTCTTGCTCCTACCGGGTGTCCGAGGGTTCAGC
GCCGCCGTCTTTGAGAACCTTTTCTATCATCACCTCATCGGAGACACCAGCGTCACGTCC
GTTCTCCGGGAACTCATGGTCCGGTAG

>L_var_22210
MKVGELPFTLHRRSCPWSHSTTSATTTTDQNSNNQPSPTHQRPSPLTALL
FEANESSSSPTPSTGPPTPPSSPPDPAPTSTATTTTPQIFASQRCSCPCS
DCVKLVRDSTCGMDMDLARWLTKCAEFGTGHTSLANINVHDRQLLVCSAW
KDIVILYMAQNKFNFDVVYSSMHEYLQHVPIHQYVQTYTTHHTVDAQFQA
DFIAKTGLVSQTYVHPLRRGSDFGIPTMRTVNQLQFTLSKLNQMQLDSKE
YAVLRILLLLNSDLHSLTDSRAIEVASEKVHTALMEYETLKYPDDPLRLS
HMLLLLPGVRGFSAAVFENLFYHHLIGDTSVTSVLRELMVR-

> VIRT-34411:5'3' Frame 1, start_pos=7
MKVGELPFTLHRRSCPWSHSTTSATTTTDQNSNNQPSPTHQRPSPLTALL
FEANESSSSPTPSTGPPTPPSSPPDPAPTSTATTTTPQIFASQRCSCPCS
DCVKLVRDSTCGMDMDLARWLTKCAEFGTGHTSLANINVHDRQLLVCSAW
KDIVILYMAQNKFNFDVVYSSMHEYLQHVPIHQYVQTYTTHHTVDAQFQA
DFIAKTGLVSQTYVHPLRRGSDFGIPTMRTVNQLQFTLSKLNQMQLDSKE
YAVLRILLLLNSDLHSLTDSRAIEVASEKVHTALMEYETLKYPDDPLRLS

HMLLLLPGVRGFSAAVFENLFYHHLIGDTSVTSVLRELMVR

**LV Dmrt2**
**>chr11**
**Length = 36215758**

**Score = 1899 bits (958), Expect = 0.0**
**Identities = 958/958 (100%)**
**Strand = Plus / Plus**

**3879607-3880564**

```
>L_var_18272-RA transcript offset:0 AED:0.05 eAED:0.08 QI:0|0|0|1|1|1|2|0|522
ATGGCTGAGGGCAAGGAAACTAATGTTACGACGAGGAAAATAAGTGAATCGCTGAATACC
GAAATAATTTCTGATCCATCACTAAAGAAAAACATGACTTCTGGTCTTACAACCGAGAGA
AACAAGCACTTGGATAAGAAATCACTGCCCAAACACCAAGCATCGGTACGTCGAGTTGTT
CGGACACCTAAGTGCGCCCGATGCCGAAACCACGGGGTCGTGTCTTGCCTGAAGGGTCAC
AAGCGGTTTTGCCGCTGGAGGGATTGCCGATGCACAAATTGTCTCTTGGTGGTGGAGCGA
CAACGGGTGATGGCCGCTCAAGTCGCGCTGAGGCGACAGCAATCCAACGACCCGTCGGCT
AGCGGGAATGGAGCAGTGAAGACGCCGGGAAGCGGCTCGGCGTCGGGCGGGGAAAGAGAA
GGAGGAAACGTCAACGGCTGTAGGAAAAAGAGTGCCAAGGAGTTGGTTTCGGCTGAAGAG
CTGTCGGACAGTAAACGGAAACTGGCCGAGATAGGTGCCGAGGCTAGCCGATTGAAAGAG
AGAGTTAAGAGACTGAGTTCGACAGCAGCCAGGGGGAGAATAAATGGATCTATTGCACGG
GACATACTCGAAGGTCAACGTGGCAAGCCAGGTCGACTGTCATCTCGCGTTCCAAGTCGC
CCAGTCATTTTCCTACCCCCGCCAGTCAGCGAGAGGATGCGCAAACGACGCGCTTTCGCC
GATAAAGAACTAGAGACAACAATGCTTCAACGCGAATGCCAGTGGACGCTGATGTTGGCG
TACGCAGCCGGTGACAAGACGCTAGCGCCTTTCGCGTCACATCCCTTCTCCTCACATTTC
GACGCAGGTCGTCACACCGGTCAAGATACGGATAAAGAACCGTTACGGAATATTCGAACC
GGATATGGAGGTCTTAATCATGTGAGTGGAGAGACGAGGTCGAATGAATCTGAACTGTTG
TGGACAGGATCGCCGTGCAATGATGTGCGACTGCAGCTAGGTGGCGGCGGTGAGGTGCCG
ACATTGTCCCCGAGCGTCGCAGTGCCTTGCATGAGAACACCGAATGCCTCGATGGTTTCA
ATTCCGTCGACGCTCATCAAACACGGTCACGGAGCTGGAGTTGGACTTGGACAATTTTCG
GGAAATCTCCACCGGAACCCGGCACTAACAACCATTGTAACGAATAAAGAACATCGTGAA
AGATCACATGAAGACACCTATCTGCATGATCCGGCAAAACTTCCGTTGTCTAGGTGCAAA
AGACTAAAAGGTGACGAAAGGTTAGGCAGTTCTTCGCAGAATAGTAGGTTTGGTGGAGAG
TTCGACAGCCGCCCAGAAGAAGCAGACTTTAGGGGTTCAGGTTCGATGGACAGAGATACT
GAACATGCTAAGTGGAGGGTTGTGGGTCGTGATCATGACATGCATGGACATGGAAAACAA
CTTGGTCAGGATAGCGCGGAAATTGGACGCGACGGATCAGAGTTTGAAGCGCAGGATTTC
TGTGAAAAGACAATACAAGCGTCAACAAATAATCTAGCATTTTCCATTGAGAGCTTGTTA
AAGAAATGA

> VIRT-145156:5'3' Frame 1, start_pos=0
MAEGKETNVTTRKISESLNTEIISDPSLKKNMTSGLTTERNKHLDKKSLP
KHQASVRRVVRTPKCARCRNHGVVSCLKGHKRFCRWRDCRCTNCLLVVER
QRVMAAQVALRRQQSNDPSASGNGAVKTPGSGSASGGEREGGNVNGCRKK
SAKELVSAEELSDSKRKLAEIGAEASRLKERVKRLSSTAARGRINGSIAR
DILEGQRGKPGRLSSRVPSRPVIFLPPPVSERMRKRRAFADKELETTMLQ
RECQWTLMLAYAAGDKTLAPFASHPFSSHFDAGRHTGQDTDKEPLRNIRT
GYGGLNHVSGETRSNESELLWTGSPCNDVRLQLGGGGEVPTLSPSVAVPC
MRTPNASMVSIPSTLIKHGHGAGVGLGQFSGNLHRNPALTTIVTNKEHRE
RSHEDTYLHDPAKLPLSRCKRLKGDERLGSSSQNSRFGGEFDSRPEEADF
RGSGSMDRDTEHAKWRVVGRDHDMHGHGKQLGQDSAEIGRDGSEFEAQDF
CEKTIQASTNNLAFSIESLLKK-
```

LV_DsxL

**>chr7**
**Length = 45571165**
**24440607-24442803**

```
>L_var_13715-RA transcript offset:0 AED:0.01 eAED:0.08 QI:0|0|0|1|1|0.5|2|0|780
GGCGCAGATGTGAAAGGTCGGGAAGATGAGTCACAAAGATGTCTCCCCCTTAAGAAAAGG
CATCCAACCTGCGCTCGATGCCGAAACCACGGACTCGTCGTGGACCTCAAAGGCCACAAG
CATTTATGCGAGTACCGTGACTGCCTCTGCACTCGATGTAACCTTGTGATTCAACGACGG
GTAGTGATGGCGAAGCAGGTAGCCCTAAGTCGAGAGCAGTTGAAGCAACAGAGACAAGAT
CACTGTACATCTCAACAGCTACAGGTCAGTCAAGGTTCCTCACAGGGTATACCATCGTCT
CCAACAGTTATTGATCTCATATCTGAAGGTACAAGGTCAGAGGAGAGGGCGAATTCCGTC
GTTCAACTTTCACCGGATTCGACAAATGTCGAATACAATGGTCCATATTCTTCTTGGGAC
TCGCCTCCGATGAATTTGTCACAGCCGACACAAATGACGTTGACTCCTCAGAATGTTTCA
AGCCAAGCTTCGCGGTCATCTCCCACATCTTTCCAAGCAGTTCAGGAAAGCGAAACGTTT
AGACCAGGAATGGTCTTCTTCACAAGCCCTTCGAATAGATCGTTACACCCCGTGGTACCT
TACCAATCTAGTGCCCCGAGAATTGTCTCTAACAACGACAATATCCCCGGAGCTTCGGCC
TTTATCAACTATGCGTCGTCTCAAGGCAATTTTGACCACGCGCTAGAGTACTCTCCCACG
TCATATCACACGCCTCAATTAAATGAAATGTATCGACCCCAGCCTTTAGTCGGTGCCAAC
CAAATGCCAAGACCATTCGTACCAAGTCCCGCCATACAGATGGCATCTAACGAATCTATC
AACCAACATATTGGATATATTCCCAGTCAAACGGCAACCTATTCCGCCAACCTTTCGTCA
GCTCATGGTGTTACAGGACAAACGTCAGAGTATCCCGCCAACCATTTGTTCGCTCATGGT
GTTACAGGACAAATAGCAGCCTCGTCCGCCAACCATTTGTTAGCTGAGGGTGTTTCAGGA
CAAGCATCAGAGTATTCCGCCAACCATATGTTAGCTCATAGTGTTACAGGACAAACGACA
GCCTATACCGCCAACCATTTGTTAGCTCATGGTGTTACAGGACAAACGGCAGCCTATTCC
GCCAACCATTTGGCTCAGGGTGTTACAGGACAAGCATCTGAGTATTTCGCCAACCATTCG
TCAGCTCATGGTGTTTCAAGAGAAATAGAGTTTCATCCAAGATCAACATTCCATTCATCA
CTGGTAGATGAGACGTACCGGTGCCAAGTATCGATAACTTCAAATTATCCAACTGGACCA
GCTTTCCATTGTCCCGTGGTGCCACATCAACCATTTGACGCAGTCTCTGCATATCGCGAT
AGTTACATGTTGGCAAACTCACGGGTGGCTTCTAGCTCCGCCATGCATATCGATTCAATG
TCGGAGACCCAACCGCCTTCAGAGGAAGCAACAGACCTGTCTAATATTCCGTACTGTCAA
GTTAAAGAGCATGCGGACCTGCTGCATCAGACACCGCTTTCAAGTTCGTTACGGGTATCA
TCATTCGTAAACCAATCAGCACAAAACTTCATGCCTGCCCAAGAATATTACAGCAATCCA
GATAGATATCCATCATGCTCTTTGCCAGCAGATTCGGCAACAGCAGCGTATTTATTTCAA
ACAGGAGGTAGAGCTATGTTATCTTCTGAGAGTGATAACTCATTCCAGGGTGCCGACTTT
ACGAGACGGCCCCAAATCGATACAACACCACTGCAACCAGAAGACACCCCTCTGAATCAC
GCTGACTTTTCTTCTAGGTCAGTACCTTTAACTACAACACAGCAAGAAATCGCGGTTAAT
ATTTCAAAAGAACCAACCTACAAATCTCGTCGACCTTTTCCTCTTCGGATCACCATCCCA
TCCCTGAATCTATCAGATAGATTCAACGACAGTCTGTCTCCAAGTCCCATGGACCCATCG
CCAACTTTCACACCATCAAACTTAAGCATTCTCACTAACACCTCTCTGCCGACAAGTGAT
ATTCCGTCTCCTGGATTCTTTACTGCGTCTCTCCCTTCATTCAACTACCCATCTCCGACG
ATGCCCACTCAGAGCTCTCCCGTCTTTAGTTACCCATCTCCTTACACCAACCCCTCTAAG
CATGCAGGATTGCCGTCTTTCCAAAGTCTGACGGAATCGATTTTACCTGTGACGCAAAGC
AGCCTGTTGCCTTCAAATTTCGCGGAGAAGGTTCGGATCCCGCCGAAATATGACGAAGAC
TTTGAAAATGATTACGAGAGTGACGAAATTGACGTTTGCTCCTTGAACCATTATGCTCAG
TGA
```

```
> VIRT-142159:5'3' Frame 1, start_pos=62
MAKQVALSREQLKQQRQDHCTSQQLQVSQGSSQGIPSSPTVIDLISEGTR
SEERANSVVQLSPDSTNVEYNGPYSSWDSPPMNLSQPTQMTLTPQNVSSQ
ASRSSPTSFQAVQESETFRPGMVFFTSPSNRSLHPVVPYQSSAPRIVSNN
DNIPGASAFINYASSQGNFDHALEYSPTSYHTPQLNEMYRPQPLVGANQM
PRPFVPSPAIQMASNESINQHIGYIPSQTATYSANLSSAHGVTGQTSEYP
ANHLFAHGVTGQIAASSANHLLAEGVSGQASEYSANHMLAHSVTGQTTAY
TANHLLAHGVTGQTAAYSANHLAQGVTGQASEYFANHSSAHGVSREIEFH
```

PRSTFHSSLVDETYRCQVSITSNYPTGPAFHCPVVPHQPFDAVSAYRDSY
MLANSRVASSSAMHIDSMSETQPPSEEATDLSNIPYCQVKEHADLLHQTP
LSSSLRVSSFVNQSAQNFMPAQEYYSNPDRYPSCSLPADSATAAYLFQTG
GRAMLSSESDNSFQGADFTRRPQIDTTPLQPEDTPLNHADFSSRSVPLTT
TQQEIAVNISKEPTYKSRRPFPLRITIPSLNLSDRFNDSLSPSPMDPSPT
FTPSNLSILTNTSLPTSDIPSPGFFTASLPSFNYPSPTMPTQSSPVFSYP
SPYTNPSKHAGLPSFQSLTESILPVTQSSLLPSNFAEKVRIPPKYDEDFE
NDYESDEIDVCSLNHYAQ-

Lv Doublesex Primers

| | fw1 | rev1 | fw2 | rev2 |
|---|---|---|---|---|
| **LV_Dmrt1T7** | AAGTGTCCCAGAGCTACCGA | TAATACGACTCACTATAGGGCTCGGGTACTGCGTGTAGTC | AAACCAACCGAAGACGGTGA | TAATACGACTCACTATAGGGTTCGGAATCACTTCCGGTGC |
| **LV_DmrtA2T7** | GATCATCCGAGCTCCAGCAA | TAATACGACTCACTATAGGGGCCGAAGTGTAAGGGTTCCA | GTCACTATCGCCAGTGGCTT | TAATACGACTCACTATAGGGCGTTGGGTCTGGTGTTGTCT |
| **LV Dmrt2T7** | GGCTGAGGGCAAGGAAACTA | TAATACGACTCACTATAGGGATCTTGACCGGTGTGACGAC | TGAAGGGTCACAAGCGGTTT | TAATACGACTCACTATAGGGAAACCATCGAGGCATTCGGT |
| **LV_Dmrt1BT7** | GATCCGGTCGATGTGCTAGA | TAATACGACTCACTATAGGGGCCCGGAGACCAAGAATGAT | TGCCCGTACACTCATCAACC | TAATACGACTCACTATAGGGGAAGGCGGCAATTCGACATC |
| **LV_DsxLT7** | GAAAAGGCATCCAACCTGCG | TAATACGACTCACTATAGGGCGGGACTTGGTACGAATGGT | GCAATTTTGACCACGCGCTA | TAATACGACTCACTATAGGGCGAATCTGCTGGCAAAGAGC |

### Supplemental Data Part 4: Nanos Orthologs

>L_var_05060
ATGAGAGGAGGTACGGCAGCCCGGCCACCGTCATGGTGCGTGTTTTGTAAAAACAATGGT
GAAAGTGAGTTGGTCTACGCCAGCCACAAGTTGAAGTCCGAGGATGGCATCACTACCTGC
CCAATACTAAGGGCGTACACCTGTCCGCTCTGTGGGACGAATGGGGACCGCGCCCACACC
ATCAAATACTGCCCGGTCAATAAGGCCAAGGAGGATGTCGGACCAGGTACCATCGGACAG
ACGTCGCAATATCGTACGCCACGCACATCAACCGGTCGAAGACGAACCTCCAGCATCAGT
AATGCCAGCACCAGCGGCTTCAACTAG

>L_var_05060
MRGGTAARPPSWCVFCKNNGESELVYASHKLKSEDGITTCPILRAYTCPL
CGTNGDRAHTIKYCPVNKAKEDVGPGTIGQTSQYRTPRTSTGRRRTSSIS
NASTSGFN

>L_var_08463
ATGAGACTAGTGCTGAGACTAGCTCTGGTCAAGAGGTTAAGCTCTGGCTGGTCACCTCTG
ACCGTCCACATACCACATGGATGTGATATGAAGCATATAAGTAAGGTTCTAGGGAAAGTA
ACCTGCCCTGTTCTACAGGCTTACGCATGCCCCATCTGCGAAACAAATGGCGACAATGCC
CATACAATCAAATACTGCCCCTTGGAAACGGACAAGTTCAGAGAATCTTCCTACATTAGG
AGTAAGGTTCTAGGGAAAGTAACCTGCCCTGTTCTACAGGCTTACGCATGCCCCATCTGC
GAAACAAATGGCGACAATGCCCATACAATCAAATACTGCCCCTTGGAAACGGACAAGTTC
AGAGAATCTTCCTACATTAGGAGTAAGGTTCTAGGGAAAGTAACCTGCCCTGTTCTACAG
GCTTACGCATGCCCCATCTGCGAAACAAATGGCGACAATGCCCATACAGTCAAATACTGC
CCCTTGGAAACGGACAAGTTCAGAGAATCTTCCTACATTAGGAGTAGGGCCAGCCTCTCC
TCTGCAGAAATACTCCTCCTCTGA

>L_var_08463
MRLVLRLALVKRLSSGWSPLTVHIPHGCDMKHISKVLGKVTCPVLQAYAC
PICETNGDNAHTIKYCPLETDKFRESSYIRSKVLGKVTCPVLQAYACPIC
ETNGDNAHTIKYCPLETDKFRESSYIRSKVLGKVTCPVLQAYACPICETN

GDNAHTVKYCPLETDKFRESSYIRSRASLSSAEILLL

>L_var_19332
ATGGGTACTGCAACCCGACCGCCTAGCTGGTGTGTGTTCTGTAAGAACAACGGGGAAAGC
GAGATGGTGTATGCCAGCCACAAGTTGAAGTCCGAGGATGGCATCACTAGCTTTGAGAAC
TCTGGCGATTTTAAGAACTCTATCTGTAGTCATTTGCACAACGATGATTTCAACTCTAAT
GGAGTCGATGTTCGTGTCATCACGAACCCGGCTGTGATGAGCGATCTCAATAACGTCATT
TCTGACCTCCCTGTGTCGACTGCCAGTATAGCGTACTCTGGATTCCTACCACTTAACATC
AATATTACTGAACTTTCGAAAGTGTTGAGAGGAGCGGGTCGACGCTCTAATAACGGTAGA
GGTGGAGCAACGGGTACTGCAACCCGACCGCCTAGCTGGTGTGTGTTCTGCAAGAACAAC
GGGGAAAGCGAGATGGTGTACGCCAGCCACAAGTTGAAGTCCGAGGATGGCATCATTAGC
TTTGAGATTAGACCGAGAAAGTGGAGAGGTGGAGCAACGGGTACTGCAACCCGACCGCCT
AGCTGGTGTGTGTTCTGTAAGACCAACGGGGAAAGCGAGATGGTGTACGCCAGCCACAAG
TTCAAGTTGGAGGATGGCATCACTAGCTTTGAGAAAGAGCACACCACCATGATGAGAGGA
GCGAGTCGACGCTCTAATAACGGTAGAGGTGGAGCAACGGGTACTGCAACCCGACCGCCC
AGCTGGTGTGTGTTCTGTAAAAACAACGGGGAAAGCGAGATGGTGTACCCCAGGCACAAG
TTGAAGTCCGAAGATGGCATTTACTAG

>L_var_19332
MGTATRPPSWCVFCKNNGESEMVYASHKLKSEDGITSFENSGDFKNSICS
HLHNDDFNSNGVDVRVITNPAVMSDLNNVISDLPVSTASIAYSGFLPLNI
NITELSKVLRGAGRRSNNGRGGATGTATRPPSWCVFCKNNGESEMVYASH
KLKSEDGIISFEIRPRKWRGGATGTATRPPSWCVFCKTNGESEMVYASHK
FKLEDGITSFEKEHTTMMRGASRRSNNGRGGATGTATRPPSWCVFCKNNG
ESEMVYPRHKLKSEDGIY

>L_var_19333
ATGAGAGGAGCGAGTCGACGCTCTAATAACGGTAGAGGTGGAGCAACGGGTACTGCAACC
CGACTGCCCAGCTGGTGTGTGTTCTGTAAGAACAACGGGGAAAGCGAGATGGTGTACGCC
TGCCACAAGTTGAAGTCGGAGGATGGCATCACTAGCTTTGAGATGATGAGAGGAGCGAGT
CGACGCTCTAATAACGGTAGAGGTGGAGCAACGGGTACTGCAACCCGACCGCCCAGCTGG
TGTGTGTTCTGTAAAAACAACGGGGAAAGCGAGATGGTGTACCCCAGGCACAAGTTGAAG
TCCGAAGATGGCATTTACTAG

> L_var_19333
MRGASRRSNNGRGGATGTATRLPSWCVFCKNNGESEMVYACHKLKSEDGI
TSFEMMRGASRRSNNGRGGATGTATRPPSWCVFCKNNGESEMVYPRHKLK
SEDGIY

>L_var_19334
ATGGCAGATAATACAATCACAACTAGAGCTGGGAAAGAAAAGAAAGAATCAACAGCCTCT
CTGATCTCTGCTGTCATTAAGGATATTCTACATGACAAGCGATTCCTGGACCTAGTTACA
AACCATGTTGATATTCAACTCAACAAATTTCAATCAGTGCTTGAAGAACAAGAGGCAAAG
ATATTGGAGTTGGAGGTGCAAAATGAAAAGAAATCACATGAACTTGGCCAACTAAAGTCA
AAACTTGATCAACAAAGCGACCAATTACGGAAACTACAACATGAAACCAACAGGCAAGAG
CAGTACAGTCGCCGCAACTGTTTGCGCTTCTTTGGTATAGAAGAATCAACAGATGAGGAT
ACCGACATGAAGATTATTCAGCTGGCAAATGAATCTTTAGGAGTGCCATTGACCAAGGAT
GACCTGGAAAGAAGTCATCGCATTAACTTCAAAGCCAACATGACGCCTGAGGAAAGGAAG
AAACGAGTGCGCCCAATCATAGCGAAGTTTGTTTCTTATCGTAAAAGATCAGAAGTCTTA
AAACACAGAAGAAAACTTGCAGGTCTACGGAAATCTATCCAAGAGGACCTCACAAGTGAA
AATGCCAAACTCTTGCAAAAGACCAGAGATAGTGAAAAGGTCAAATCAGCATGGACTCGC
GATGGTCGAATTGTTGCATTGATTCCAACATCGAACAACCATGCAATAACAAAGATCATC
TCATGTGCTAAAGACCTGGACCTTATGTGA

>L_var_19334
MADNTITTRAGKEKKESTASLISAVIKDILHDKRFLDLVTNHVDIQLNKF
QSVLEEQEAKILELEVQNEKKSHELGQLKSKLDQQSDQLRKLQHETNRQE
QYSRRNCLRFFGIEESTDEDTDMKIIQLANESLGVPLTKDDLERSHRINF
KANMTPEERKKRVRPIIAKFVSYRKRSEVLKHRRKLAGLRKSIQEDLTSE
NAKLLQKTRDSEKVKSAWTRDGRIVALIPTSNNHAITKIISCAKDLDLM

>L_var_19343
ATGAGAGGAGCGAGTCGACGCTCTAATAACGGTAGAGGTGGAGCAACGGGTACTGCAACC
CGACTGCCCAGCTGGTGTGTGTTCTGTAAGAACAACGGGGAAAGCGAGATGGTGTACCCC
AGCCACAAGTTGAAGTCGGAGGATGGCATCACTAGCTTTGAGAACTCTGGCGATTTTAAG
AACTCTATCTGTAGTCATTTGCACAACGACGATTTCATCTCCAATGGAGTCGATGTTCGT

GTCATCACGAACCCGGCTATGATGAGTGATCTCAACAACGTTATGTCCGGTTTCCCCGTG
TCGACTGCCAATACAGCATACTCAGGATTCCTACCACTGAACATTGATATTACTGAACTT
TCGAAAGTGATGAGAGGAGCGAGTCGACGCTCTAATAACGGTAGAGGTGGAGCAACGGGT
ACTGCAACCCGACCGCCCAGCTGGTGTGTGTTCTGTAAGAACAACGGGGAAAGCGAGATG
GTGTACGCCTGCCACAAGTTGAAGTCGGAGGATGGCATCACTAGCTTTGAGAACTCTGGC
GATTTTAAGAACTCTATCTGTAGTCATTTGCACAACGACGATTTCATCTCCAATGGAGTC
GATGTTCGTGTCATCACGAACCCGGCTATGATGAGTGATCTCAACAACGTTATGTCCGGT
TTCCCCGTGTCGACTGCCAATACAGCATACTCAGGATTCCTACCACTGAACGTTGATATT
ACTGAACTTTCGAAAGTGATGAGAGGAGCGAGTCGACGCTCTAATAACGGTAGAGGTGGA
GCAACGGGTACTGCAACCCGACCGCCCAGCTGGTGTGTGTTCTGTAAAAACAACGGGGAA
AGCGAGATGGTGTACCCCAGGCACAAGTTGAAGTCCGAAGATGGCATTTACTAG

>L_var_19343
MRGASRRSNNGRGGATGTATRLPSWCVFCKNNGESEMVYPSHKLKSEDGI
TSFENSGDFKNSICSHLHNDDFISNGVDVRVITNPAMMSDLNNVMSGFPV
STANTAYSGFLPLNIDITELSKVMRGASRRSNNGRGGATGTATRPPSWCV
FCKNNGESEMVYACHKLKSEDGITSFENSGDFKNSICSHLHNDDFISNGV
DVRVITNPAMMSDLNNVMSGFPVSTANTAYSGFLPLNVDITELSKVMRGA
SRRSNNGRGGATGTATRPPSWCVFCKNNGESEMVYPRHKLKSEDGIY

>L_var_19345
ATGGGTACTGCAACCCGACCGCCTAGCTGGTGTGTGTTCTGTAAGAACAACGGGGAAAGC
GAGATGGTGTATGCCAGCCACAAGTTGAAGTCCGAGGATGGCATCACTAGCTTTGAGAAC
TCTGGCGATTTTAAGAACTCTATCTGTAGTCATTTGCACAACGATGATTTCAACTCTAAT
GGAGTCGATGTTCGTGTCATCACGAACCCGGCTGTGATGAGCGATCTCAATAACGTCATT
TCTGACCTCCCTGTGTCGACTGCCAGTATAGCGTACTCTGGATTCCTACCACTTAACATC
AATATTACTGAACTTTCGAAAGTGTTGAGAGGAGCGGGTCGACGCTCTAATAACGGTAGA
GGTGGAGCAACGGGTACTGCAACCCGACCGCCTAGCTGGTGTGTGTTCTGCAAGAACAAC
GGGGAAAGCGAGATGGTGTACGCCAGCCACAAGTTGAAGTCCGAGGATGGCATCATTAGC
TTTGAGAGTGAGGGCACCACCATTAGACCGAGAAAGTGGAGAGGTGGAGCAACGGGTACT
GCAACCCGACCGCCTAGCTGGTGTGTGTTCTGCAAGACCAACGGGGAAAGCGAGATGGTG
TACGCCAGCCACAAGTTCAAGTCGGAGGATGGCATCACTAGCTTTGAGAAAGAGCACACC
ACCAACTCTGGCGATTTTAAGAACTCTATCTGTAGTCATTTGCACAACGATGATTTCAAC
TCTAATGGAGTTGATGTTCGTGTCATCACGAACCCGGCTATGATGAGTGATCTCAATAAC
AACGTCATATCTGACCTCCCCGTGCGACTGCCAATACAGCGTACTCAGGATTCCTACCAC
TGA

> L_var_19345
MGTATRPPSWCVFCKNNGESEMVYASHKLKSEDGITSFENSGDFKNSICS
HLHNDDFNSNGVDVRVITNPAVMSDLNNVISDLPVSTASIAYSGFLPLNI
NITELSKVLRGAGRRSNNGRGGATGTATRPPSWCVFCKNNGESEMVYASH
KLKSEDGIISFESEGTTIRPRKWRGGATGTATRPPSWCVFCKTNGESEMV
YASHKFKSEDGITSFEKEHTTNSGDFKNSICSHLHNDDFNSNGVDVRVIT
NPAMMSDLNNNVISDLPVRLPIQRTQDSYH

GTCTGGTCACCCTGAAAAATTAGGTGGGGCTAAATGGCAATTTAGCCCCACCTATAGTAAGGGGTTAAGGAAATTGTAAA
TAGGGTATTATTCAATTTCCAAACGAGTTTGATCTGAAAGATCCCATCGACCACCGCTCCTTTCACTGAAAGTAAAATGT
TTAGATTTCTAGACTATTTTATCTCCAGCCACACCGTTCATCCCAGAGGTTAGAAGTTGCGTTCGATCTCATTTATCTCT
ATTGTCCAGTCTTGGTCATTTATTTGAGCTTAATCAACGCTTCAGCTGAATGTAAATTTCTCTTAAAATGAGAATAATTG
AATAATTATGTTGCAAATTTTTATAATCATTGATCTTCTTCAAAATGTGATACCATCTCACTGTGCTAGAATCAACTTTG
ATACAAGCAAAATATATTTAATGTTTAGTGCCGCCTGAAATACAACTAAGCAGGGGTTATTCCCCCACCATCGTGAAAGT
TGGGGCGATGAGAAAATTGCGTAATCTCGAACTTTGCTTGACCGCAAAGTGGGGGTTCAGTTTGCAGTATGTAGTTCGCC
ATATCAATTTCGACGCATTGCATACTTTGAGTATCGTCCACTGCACTCGTGTCCTGCTTTCCGAGCTCAAGCTTCAGAAA
GTTTTTAAGATCGTACGATTACTATGGAGACTTGGATGTGTTATTCACGGACGGTGCCGACATCCCCATGCTCCCCGCCA
CAGCCTCAGACTCAGAGTCCGCAATCCCCGCAGTTCTCCTGGCCATTACCTGTAACCCCATCACGCACCGTATCATCACC
GCCACCTGTCCCATCACCACCTCCCTCACCTTCTCCGCATCATGAAAGGAAGAAATATCTGAACGTGTATTGCGTTTTTT
GCAAGAACAATAAAGAGACGTTAAGCTTTTACAGCTCACATGTTTTGAAGGACGATGTAGGGAAGGTAACCTGCCCTGTT
CTTCGGGCGTACACATGCCCCATCTGCGGGGCCAATGGGGACAGTGCCCATACCATAAAATACTGCCCCATGGCATGCAT
GGAATCGGACAAATTTAGGAATAGGAGCAGAAGGGCATCTTTGTCCAAATGGACAGAAAGTTCCTCACCCTGAAGAAACA
AGAATCCAAAGACCATAAGGTAAAGTGTGATATTCGCCACTTCTTTACGAAACGAAGGAAAATTTTAGAAATTGAGCTAA
TTTAGTAATTATTTAATTAATTTTGGAGTTTGGACCCCCTTAGGCCTTACATGTTCTAAGCTTCTGTTTAGAAAATTCGT
AGATTCATAGTTTCTCTTCCGCATCAGACATTGCAGCTGGTGATATTATTTTAAGGTTTTTCCTGCTGTTTAATCGAATC
ATCATATAAATGTACATGGAATATAAACGAAATCGCATCCGAGATTTTCATTATATGTCCTTGATTACATTTGATATGGA
AATGGAATTCTGTTCAAACTCAAGGGGCTAATTCATGAGGGGGTGGCGGGGGGTTGAAATTGTCGACCATAAATCAGACA
ATTAAAAAAAGATAAACATTAAAGTAAAACTAAATGGACAGATCTACCACAAAAGTAAGTTGATTTTAATAAAAAGAGAA
AAATCCAAACAGCAAACTTACGAAAATTCATCAAAATCAGGTGAAAACTAGTTAGAAAGTTGTGACAATTTTAAGTTCGC

TTTAGTTAAATATCACAGAACAGTTTTACATCTACCCATTGTGGGCTGCAAATGAGGGGAGTGATGATTTCACTCACTAT
TTCTCTTGTATTTTATCATCTGAGATATGAATAATTCTAATTTTTCCTCATTGTCCCCTTTAATCCCCCTGAACATGGGT
AATTACCCATTTTAAAAAAAATTAACTGTTCAAGGTTCAGTCATGTTTGTCATTGACAAATATGTAAGAATTGAAATATT
GTACAATTCAAAGAATAAAAAACAAAAGAAAAGGTGAGTGCGGGATATCATCGAATCTCTCATTTGCATGTCTATATTTT
TTTGTATAACTGTTTTCTGAAAAATAAGCGAAACTTTAAAATGTCATAACTTTCTTAAATACATCCGATTTTGATGAAAT
TTTGGGCATAATCATGATGATGATGTTTGTTTAATATCTCTGGGATGGACTTGACCTTTAAATATTCGGCATGTTTATGC
TTAGGTTCCCTTTCTTGTTGTACTTGTGGAATTTAAACCGTCCCCAAAAATGAAGAAGGAAAAAATTACCCATCTTCCTT
TTTTGACCTGGATACACCGTCTGCATCCCCTTCTCCTTTCATTCACGGATTGGAGATTGTTTTATTTCTCTAATCTGCTG
ATCAGGGGAGCGTTTCATGAAAGGATTTTATCCGACAAGTCTTGTTTTATCCGACAATTACCATAGTAACAGAGTACCTG
TCAGCCAATCAGAATCAGGGAAACTTGTCAGATCTGACAACTTGTCGCACAAAAATGTTGATGAAACGCTCCCCAGGGGC
CAGAAAGAGTTGCAATCAATCGCAGCTTTAAAAAAAAATTGCGTAACTTGATTTTCAACCGATCAACGGCGCGCATTTTGG
ACTTGAGATTGATTTTTTGACGGCAATCCTGTATTTTAGGGGACTAAATTATTTTAGGTTGCTAACGTCAGTTCAGGCAA
CAGCTCCACGACCGCTGTTAACGGTAGTTCAGTAGTGCGTACGTACGTGCCGGGTATGGCAGGCCGTTTATTATTTCTTC
ATTCTATAACCCCCCCCCCCCGCCATGACAAGTAACGCTGATTACTGTTGGTTTGATATAAATTACCATTATACTCTTCA

>L_var_08461
ATGCCCATACAATCAAATACTGCCTCATGGAATTGGGGCAGTCCAGAGAATCTTCCTAGA
AGTGGGGCCAGCCTCTCCTCTGCAGAAATACTCCTCCGACACACCTTGAAGAAATACCTG
AACACCTACTGTGTCTTTTGCAAGAACAACAGAGAAACTCTCAGCTTCGACAGCTCCCAT
GTTTTGAAGGATGATCTAGGGAAAGTAACCTGCCCTGTTCTATGGGCTTACACATGCCCC
ATCTGCGGGGCCAATGGTGACAATGCCCATACAATCAAATACTGCCCCTTGGAATTGGAC
AAGTCAAGAGAATCTTCCTAG

>L_var_08461
MPIQSNTASWNWGSPENLPRSGASLSSAEILLRHTLKKYLNTYCVFCKNN
RETLSFDSSHVLKDDLGKVTCPVLWAYTCPICGANGDNAHTIKYCPLELD
KSRESS

>L_var_08462
ATGGGCTTACACATGCCCCATCTGTGGAGCCAATTGAAGAGATACCTGAACACCTACTGT
GTCTTTTGCAAGAACAACAGAGAAACTCTCAGCTTCTACAGCTCCCATGTTTTGAAGGAT
GATCTAGGGAAAGTAACCTGCCCTGTTCTACGGGCTTACACGTGCCCCATCTGCGGGGCC
AATGGTGACAATGCCCATACAATCAAATACTGCCCCTTGGAATTGGACAAGAAGAAATAC
CTGAACACCTACTGTGTCTTTTGCAAGAACAACAGAGAAACTCTCAGCTTCGACAGCTCC
CATGTTTTGAAGGATGATCTAGGGAAAGTAACCTGCCCTGTTCTATGGGCTTACACATGC
CCCATCTGTGGAGCCAATTGTGACAATGCCCATACAATCAAATACTGCCTCATGGAATTG
GGGCAGTCCAGAGAATCTTCCTAG

> L_var_08462
MGLHMPHLWSQLKRYLNTYCVFCKNNRETLSFYSSHVLKDDLGKVTCPVL
RAYTCPICGANGDNAHTIKYCPLELDKKKYLNTYCVFCKNNRETLSFDSS
HVLKDDLGKVTCPVLWAYTCPICGANCDNAHTIKYCLMELGQSRESS

**>L_var_19628**
**ATGTGTTATTCACGGACGGTGCCGACATCCCCATGCTCCCCGCCACAGCCTCAGACTCAG**
**AGTCCGCAATCCCCGCAGTTCTCCTGGCCATTACCTGTAACCCCATCACGCACCGTATCA**
**TCACCGCCACCTGTCCCATCACCACCTCCCTCACCTTCTCCGCATCATGAAAGGAAGAAA**
**TATCTGAACGTGTATTGCGTTTTTTGCAAGAACAATAAAGAGACGTTAAGCTTTTACAGC**
**TCACATGTTTTGAAGGACGATGTAGGGAAGGTAACCTGCCCTGTTCTTCGGGCGTACACA**
**TGCCCCATCTGCGGGGCCAATGGGGACAGTGCCCATACCATAAAATACTGCCCCATGGCA**
**TGCATGGAATCGGACAAATTTAGGAATAGGAGCAGAAGGGCATCTTTGTCCAAATGGACA**
**GAAAGTTCCTCACCCTGA**

**> L_var_19628**
**MCYSRTVPTSPCSPPQPQTQSPQSPQFSWPLPVTPSRTVSSPPPVPSPPP**
**SPSPHHERKKYLNVYCVFCKNNKETLSFYSSHVLKDDVGKVTCPVLRAYT**
**CPICGANGDSAHTIKYCPMACMESDKFRNRSRRASLSKWTESSSP**

Lv Nanos Orthologs Primers qPCR

| | |
|---|---|
| L_var_05060_F | CGAGGATGGCATCACTACCT |
| L_var_05060_R | ATTGACCGGGCAGTATTTGA |

| | |
|---|---|
| L_var_08463_F | TGCTGAGACTAGCTCTGGTCAA |
| L_var_08463_R | CCCTAGAACCTTACTTATATGCTTCA |
| L_var_19332_F | TTCTGACCTCCCTGTGTCG |
| L_var_19332_R | CCGCTCCTCTCAACACTTTC |
| L_var_19333_F | AGGATGGCATCACTAGCTTTG |
| L_var_19333_R | GCCTGGGGTACACCATCT |
| L_var_19334_F | ACTGTTTGCGCTTCTTTGGT |
| L_var_19334_R | TCTTTCCAGGTCATCCTTGG |
| L_var_19345_F | CTGTGTCGACTGCCAGTATAG |
| L_var_19345_R | TGCTCCACCTCTACCGTTATT |
| L_var_08461_F | CCTCATGGAATTGGGGCAGT |
| L_var_08461_R | AGACACAGTAGGTGTTCAGGT |
| L_var_08462_F | GTGGAGCCAATTGAAGAGATACC |
| L_var_08462_R | GCACGTGTAAGCCCGT |
| L_var_19628_F | TGTTATTCACGGACGGTGCC |
| L_var_19628_R | ATGATACGGTGCGTGATGGG |

Nanos Orthologs Primers T7 WMISH

| | |
|---|---|
| L_WMT705060F | GGGCTATTGTCCAACTGCAA |
| L_WMT705060R | TAATACGACTCACTATAGGGCCATTCGTCCCACAGAGC |
| LVT7WM19334_F | ATGACAAGCGATTCCTGGAC |
| LVT7WM19334_R | TAATACGACTCACTATAGGGTTGCATGGTTGTTCGATGTT |
| LVWM08461F | ATGCCCATACAATCAAATAC |
| LVWM08461R | TAATACGACTCACTATAGGGCAAGTCAAGAGAATCTTCCTAG |
| LVWM08461F | |
| LVWM08461R | |
| LVWMT719332F | CTAGCTTTGAGAACTCTGGCGATT |
| LVWMT719332R | TAATACGACTCACTATAGGGACTCGCTCCTCTCATCATGG |
| LVT7WM19333F | |
| LVT7WM19333R | |
| LVWM19345F | TCACTAGCTTTGAGAACTCTGG |
| LVWM19345R | TAATACGACTCACTATAGGGAGTTGGTGGTGTGCTCTTTCTC |
| LVWM19628F | ATGTGTTATTCACGGACGGTGC |
| LVWM19628R | TAATACGACTCACTATAGGGCAAAGATGCCCTTCTGCTCCTA |
